# Genomic inference of a human super bottleneck in Mid-Pleistocene transition

**DOI:** 10.1101/2021.05.16.444351

**Authors:** Wangjie Hu, Ziqian Hao, Pengyuan Du, Fabio Di Vincenzo, Giorgio Manzi, Yi-Hsuan Pan, Haipeng Li

## Abstract

The demographic history is a foundation of human evolutionary studies. However, the ancient demographic history during the Mid-Pleistocene is poorly investigated while it is essential for understanding the early origin of humankind. Here we present the fast infinitesimal time coalescent (FitCoal) process, which allows the analytical calculation of the composite likelihood of a site frequency spectrum and provides the precise inference of demographic history. We apply it to analyze 3,154 present-day human genomic sequences. We find that African populations have passed through a population super bottleneck, a small effective size of approximately 1,280 breeding individuals between 930 and 813 thousand years ago. Further analyses confirm the existence of the super bottleneck on non-African populations although it cannot be directly inferred. This observation, together with simulation results, indicates that confounding factors, such as population structure and selection, are unlikely to affect the inference of the super bottleneck. The time interval of the super bottleneck coincides with a gap in the human fossil record in Africa and possibly marks the origin of *Homo heidelbergensis*. Our results provide new insights into human evolution during the Mid-Pleistocene.

## INTRODUCTION

With African hominid fossils, the origin of anatomically modern humans has been determined to be approximately 200 thousand years (kyr) ago (White et al., 2003). Based on present-day human genomes, the recent demographic history of humans has been intensively studied which reveals the world-wide spread of our ancestors (Li and Durbin, 2011; Liu and Fu, 2015; Manica et al., 2007; Nielsen et al., 2017; Ramachandran et al., 2005; Stoneking and Krause, 2011; Terhorst et al., 2017). However, the ancient demographic history during the Mid-Pleistocene is still poorly investigated while it is essential for understanding the early origin of humankind. It is mainly due to limitations of existed methods since this task requires a precise estimate for the ancient demographic history. Thus a novel approach is needed to improve the inference accuracy of demographic history.

As site frequency spectrum (SFS) plays an essential role in demographic inference (Excoffier et al., 2013; Griffiths and Tavaré, 1996; Gutenkunst et al., 2009; Li and Stephan, 2006; Liu and Fu, 2020; Liu and Fu, 2015; Terhorst et al., 2017), many efforts have been made to derive its analytical formula under a predefined demographic model (Fu, 1995; Jouganous et al., 2017; Zivković and Wiehe, 2008). Therefore, to precisely infer recent and ancient demography, we developed the fast infinitesimal time coalescent (FitCoal) process (Figure 1) that analytically derives expected branch length for each SFS type under arbitrary demographic models. It is effective for a wide range of sample sizes in the analytical calculation of the composite likelihood of a given SFS. FitCoal first maximizes the likelihood with the constant size model and then increases the number of inference time intervals and re-maximizes the likelihood until the best model is found. FitCoal does not need prior information on demography, and its accuracy is confirmed by simulation. The demographic inference of FitCoal is more precise than that of PSMC (Li and Durbin, 2011) and stairway plot (Liu and Fu, 2015), and the effects of positive selection and sequencing error can be easily excluded.

**Figure 1.**
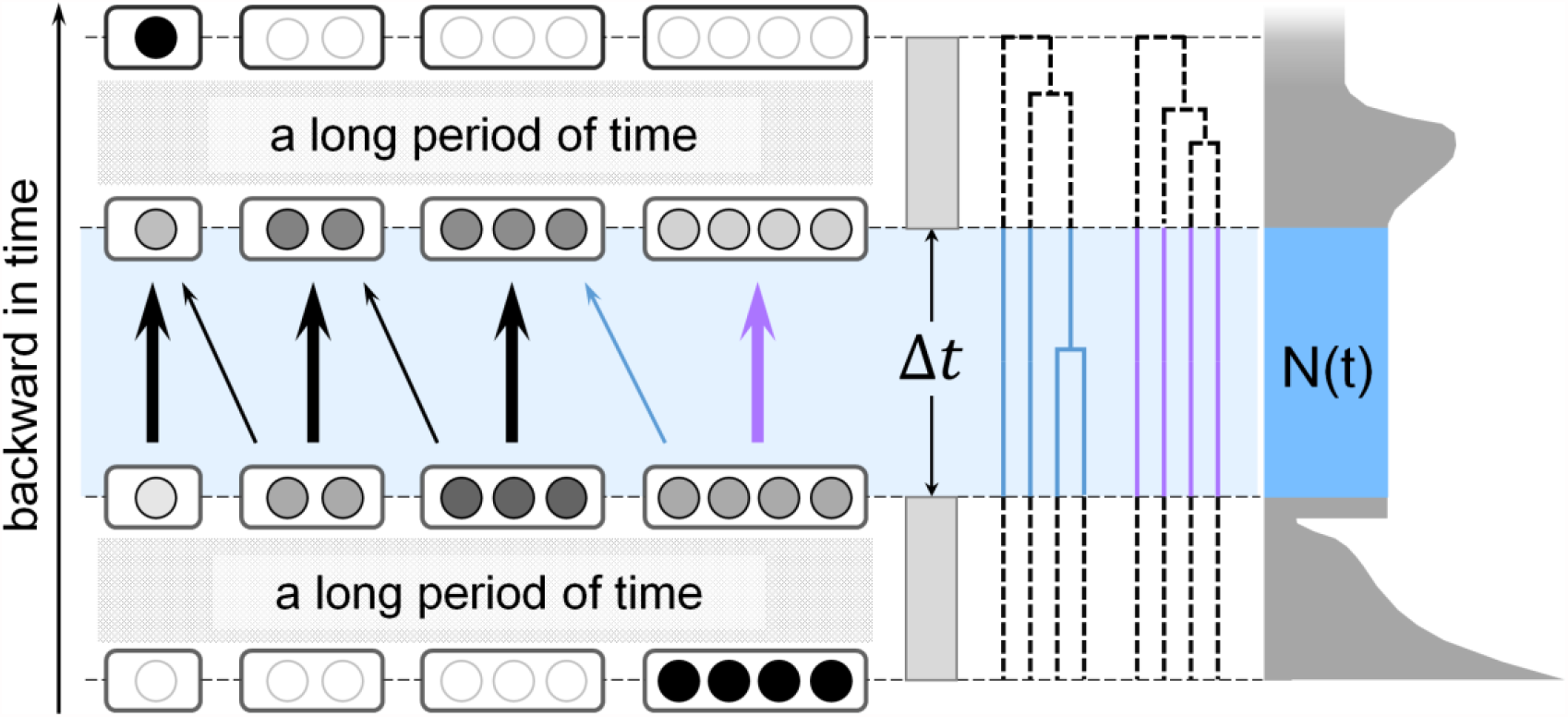
Illustration of the fast infinitesimal time coalescent (FitCoal) process. The left panel shows the backward process in which four lineages coalesce into one after passing through millions of infinitesimal time intervals. The highlighted area shows the backward transformation process of different states with tiny probability changes in an infinitesimal time interval (Δ*t*). Thick arrows indicate high transformation probabilities, and thin arrows indicate low transformation probabilities. Each state is indicated with a rounded rectangle, in which one circle indicates one lineage. The rounded rectangles with black filled circles are the states with probability 1. The rounded rectangles with empty circles are the states with probability 0. The probabilities between 0 and 1 are indicated by grey circles. The middle panel shows branches of different states. The right panel shows the demographic history of a population. The width of shadowed area indicates the effective population size, *i*.*e*., the number of breeding individuals (Harpending et al., 1998). It is assumed that the effective population size remains unchanged within Δ*t*.

We then used FitCoal to analyze large sets of present-day human genomic sequences sampled from 10 African and 40 non-African populations. The inferred recent demographic histories, including recent population size expansion/reduction and the out-of-African bottleneck, are consistent with previous studies (Altshuler et al., 2015; Bergstrom et al., 2020; Li and Durbin, 2011; Prugnolle et al., 2005; Ramachandran et al., 2005; Schiffels and Durbin, 2014; Terhorst et al., 2017). However, we found that our ancestors experienced a super bottleneck and the effective size of our ancestors remained small (about 1,280 breeding individuals) between 930 and 813 thousand years ago. The super bottleneck was directly inferred on African populations but only indirectly detected on non-African populations, which is expected by the coalescent theory. This observation, together with simulation results, indicates that confounding factors, such as population structure and selection, are unlikely to affect the inference of the super bottleneck during the Mid-Pleistocene. The super bottleneck not only explains a gap of the human fossil record in Africa between roughly 900 and 600 kyr ago (Profico et al., 2016), but also may represent a major transition in human evolution, possibly leading to the origin of *H. heidelbergensis*: the alleged ancestral species of modern humans (Profico et al., 2016; Stringer, 2016).

## RESULTS

### Fast Infinitesimal Time Coalescent Process

As analytical result of expected branch length for each SFS type is essential for theoretical population genetics and demographic inference (Excoffier et al., 2013; Fu, 1995; Li and Stephan, 2006; Zivković and Wiehe, 2008), we developed the fast infinitesimal time coalescent (FitCoal) process to accomplish the task (Figure 1). The analytical result of expected branch length for each SFS type was presented in the STAR★METHODS. For FitCoal calculation, each of millions of time intervals Δt was set extremely small, and the population size was assumed to be constant within each infinitesimal time interval. The probabilities of all states were calculated backward in time. During each Δt, the branches were categorized according to their state. For each state, the branch length was multiplied by its probability and population size and then transformed to calculate the expected branch length of each SFS type. Because the expected branch length of a SFS type is equal to the sum of the expected branch length of this type during each time interval, the latter can be rescaled and tabulated, making the calculation of the expected branch lengths extremely fast under arbitrary demographic histories. Hereafter, tabulated FitCoal is referred to as FitCoal for short, unless otherwise indicated.

### FitCoal Demographic Inference

After the expected branch lengths were obtained, the composite likelihood of the SFS observed in a sample was calculated (Excoffier et al., 2013; Hudson, 2001; Li and Stephan, 2006; Liu and Fu, 2015). As each single nucleotide polymorphism (SNP) was treated independently, FitCoal did not need phased haplotype data. When inferring demography, the likelihood was maximized in a wide range of demographic scenarios. The FitCoal likelihood surface is smooth (Figure S1), so it is efficient to maximize the likelihood. FitCoal considered both instantaneous populations size changes (Li and Durbin, 2011; Liu and Fu, 2015; Schiffels and Durbin, 2014) and long-term exponential changes of population in order to generate various demographic scenarios.

### Demographic Inference on Simulated Data

The accuracy of FitCoal was validated by simulation and comparing its demographic inferences with those of PSMC (Li and Durbin, 2011) and stairway plot (Liu and Fu, 2015) (Figure 2). Six demographic models, examined in the former study (Liu and Fu, 2015), were considered by simulating 200 independent data sets under each model. The medians and 95% confidence intervals of demography were then determined by FitCoal with the assumption that a generation time is 24 years (Liu and Fu, 2015; Scally and Durbin, 2012) and the mutation rate is 1.2 × 10^−8^ per site per generation for human populations (Campbell et al., 2012; Conrad et al., 2011; Kong et al., 2012; Liu and Fu, 2015).

**Figure 2.**
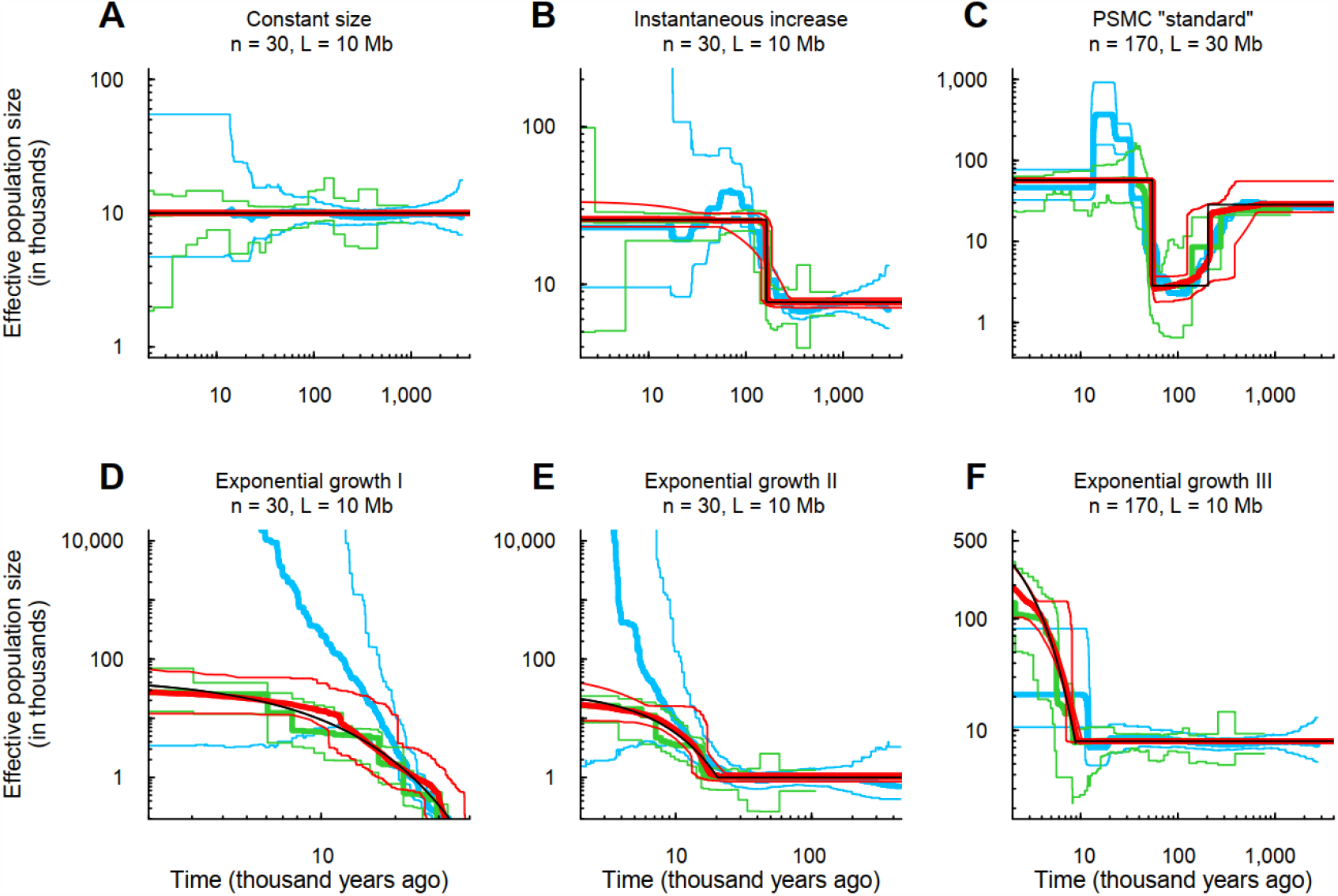
Demographic histories estimated by FitCoal, stairway plot, and PSMC using simulated samples. (**A**) Constant size model. (**B**) Instantaneous increase model. (**C**) PSMC “standard” model. (**D**) Exponential growth I model. (**E**) Exponential growth II model. (**F**) Exponential growth III model. These six models are the same as those of the previous study by Liu and Fu (Liu and Fu, 2015). Thin black lines indicate true models. Thick red lines indicate the medians of FitCoal estimated histories; thin red lines are 2.5 and 97.5 percentiles of FitCoal estimated histories. Green and blue lines indicate the results of stairway plot and PSMC, respectively, of the previous study (Liu and Fu, 2015). The mutation rate is assumed to be 1.2 × 10^−8^ per base per generation, and a generation time is assumed to be 24 years. *n* is the number of simulated sequences, and *L* is the length of simulated sequences.

FitCoal was found to precisely infer demographic histories (Figure 2). In general, the confidence intervals of FitCoal-inferred histories were narrower than those of PSMC and stairway plot, indicating a better FitCoal-demographic inference. The inference accuracy can be improved by increasing sample size and length of sequence (Figure S2). Our results confirmed that SFS allows precise recovery of the demographic history (Bhaskar and Song, 2014). The proportion of the most recent change type inferred from six models above showed that FitCoal can distinguish instantaneous and exponential changes (Table S1).

Since a demographic event may affect every SFS type, demographic history can be inferred using a subset of SFS. Results of simulation confirmed that FitCoal accurately determined demographic history based on truncated SFSs (Figures S3 and S4), thus reducing the impact of other factors, such as positive selection (Figure S5) and sequencing error, on FitCoal analysis.

### Demographic Inference of African Populations

To infer the demographic histories of African populations, seven African populations in the 1000 Genomes Project (1000GP) (Altshuler et al., 2015) were analyzed by FitCoal. Only non-coding regions, defined by GENCODE (Frankish et al., 2019), were used in order to avoid the effect of purifying selection. To avoid the potential effect of positive selection (Fay and Wu, 2000), high-frequency mutations were excluded from the analysis.

Results showed that all seven African populations passed through a super bottleneck around 914 (854–1,003) kyr ago and that this bottleneck was relieved about 793 (772–815) kyr ago (Figures 3A-C and S6; Table S2). The average effective population size of African populations during the bottleneck period was determined to be 1,270 (770–2,030). Although traces of the bottleneck were observed in previous studies, the bottleneck was ignored because its signatures were too weak to be noticed (Altshuler et al., 2015; Bergstrom et al., 2020; Li and Durbin, 2011; Schiffels and Durbin, 2014; Terhorst et al., 2017). After the bottleneck was relieved, the population size was increased to 27,080 (25,300–29,180), a 20-fold increase, around 800 kyr ago. This population size remained relatively constant until the recent expansion.

**Figure 3.**
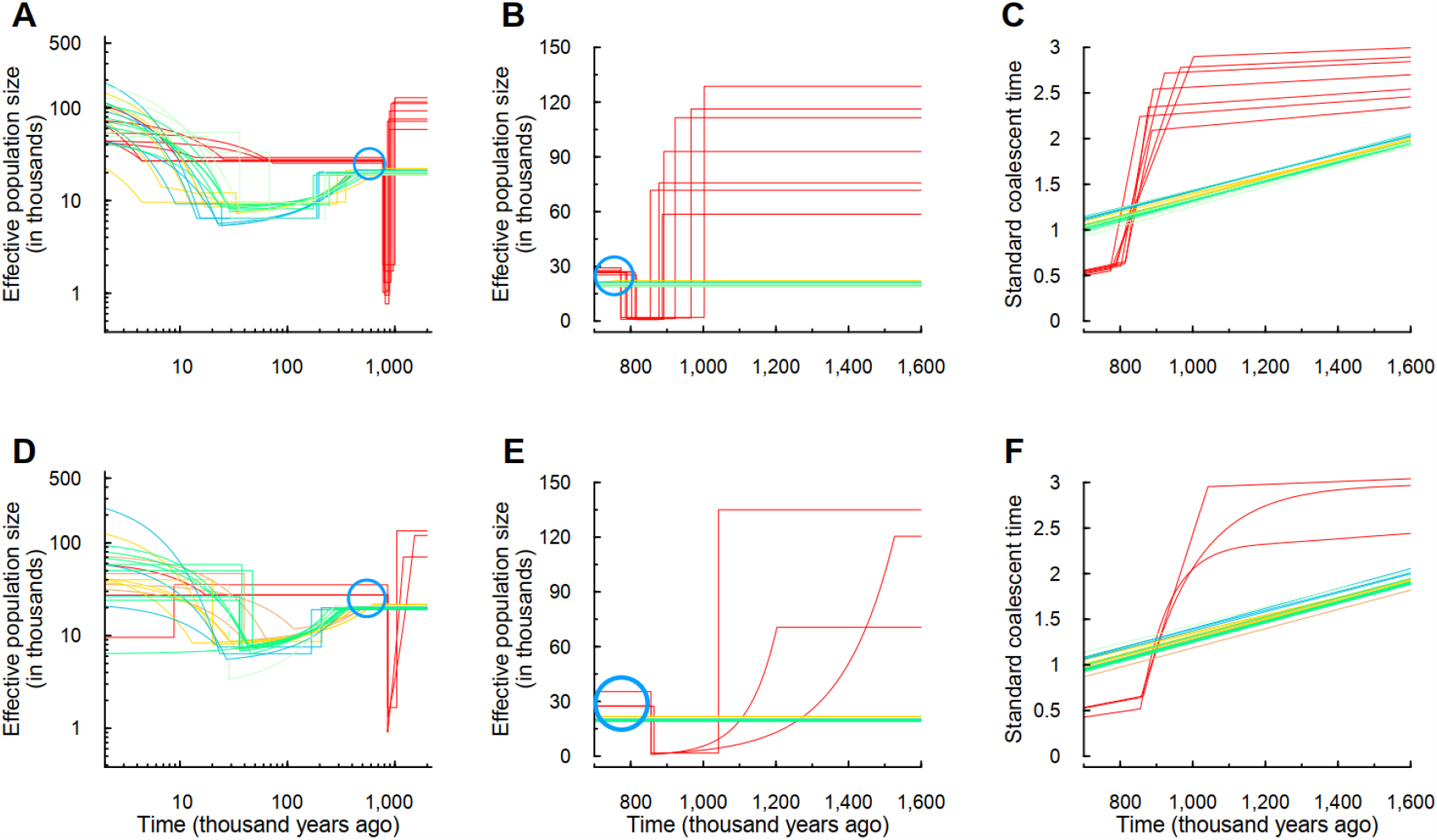
FitCoal estimated histories of human populations using 1000GP and HGPD-CEPH genomic data sets. (**A**) Estimated histories of 26 populations in 1000GP. (**B**) Linear-scaled estimation of histories of 1000GP populations during the super bottleneck period. (**C**) Calendar time *vs* standard coalescent time of estimated histories of 1000GP populations. (**D**) Estimated histories of 24 HGPD-CEPH populations. (**E**) Linear-scaled estimation of histories of HGPD-CEPH populations during the super bottleneck period. (**F**) Calendar time *vs* standard coalescent time of estimated histories of HGPD-CEPH populations. Various color lines indicate the following: red, African populations; yellow, European populations; brown, Middle East populations; blue, East Asian populations; green, Central or South Asian populations; and dark sea green, American populations. Blue circles show the population size gap between the African and non-African populations, indicating the hidden effect of the super bottleneck in non-African populations. The mutation rate is assumed to be 1.2 × 10^−8^ per base per generation, and a generation time is assumed to be 24 years.

To avoid the potential effects of low sequencing depth (∼ 5x) of non-coding regions in the 1000GP on the analysis, the autosomal non-coding genomic polymorphism of Human Genome Diversity Project – Centre d’Etude du Polymorphisme Humain panel (HGDP-CEPH) with high sequencing coverage (∼35x) was analyzed (Bergstrom et al., 2020). Populations with more than 15 individuals each were examined. Results showed that the super bottleneck occurred on all three African populations in HGDP-CEPH between 1,257 (1,042–1,527) and 859 (856–864) kyr ago (Figures 3D-F and S7; Table S3), and the average population size during the bottleneck period was 1,300 (908–1,670). This number was very similar to that (1,270) estimated from the data of 1000GP.

After the bottleneck was relieved, the population sizes of the two HGDP-CEPH agriculturalist populations were increased to 27,300 and 27,570 (Figures 3E and S7; Table S3), consistent with the 1000GP estimate of 27,280. The Biaka, a hunter-gatherer population, had a larger population size of 35,330, suggesting a deep divergence between this and other agriculturalist populations (Hsieh et al., 2016; Schlebusch and Jakobsson, 2018; Skoglund et al., 2017). The Biaka population was found to have a recent population decline (Figures 3D and S7), as previously observed (Bergstrom et al., 2020). These results suggest that hunter-gatherer populations were widely spread and decreased when agriculturalist populations were expanded.

To provide a precise inference of the super bottleneck, the results from the two data sets were combined. After analyzing the inferred time of instantaneous change of 10 populations, the super bottleneck was inferred to last for about 117,000 years, from 930 (854–1,042; s.e.m.: 23.52) to 813 (772–864; s.e.m.: 11.02) kyr ago. The effective size during the bottleneck period was precisely determined to be 1,280 (767–2,031; s.e.m.: 131). A loss of 65.85% in current genetic diversity of human populations was estimated because of the bottleneck.

### Demographic Inference of Non-African Populations

No super bottleneck was directly observed on all 19 non-African populations in 1000GP (Figures 3A-C and S6; Table S4). The ancestral population size of these populations was determined to be 20,260 (18,850–22,220), similar to that determined in previous studies (Bergstrom et al., 2020; Li and Durbin, 2011; Schiffels and Durbin, 2014; Terhorst et al., 2017). The population size of 1000GP non-African populations started to decline around 368 (175–756) kyr ago, suggesting that African and non-African divergence occurred much earlier than the out-of-Africa migration (Altshuler et al., 2015; Bergstrom et al., 2020; Li and Durbin, 2011; Nielsen et al., 2017; Schiffels and Durbin, 2014; Terhorst et al., 2017). European and South Asian populations were found to have a relatively weaker out-of-Africa bottleneck than East Asian populations, and the bottleneck severity was found to correlate with their geographic distance to Africa, consistent with the observed correlation between heterozygosity and geographic distance (Prugnolle et al., 2005; Ramachandran et al., 2005). A weak bottleneck was observed on American populations, probably because of recent admixture (Altshuler et al., 2015). All 1000GP non-African populations were found to increase in size recently.

The super bottleneck was also not directly detected in all 21 HGDP-CEPH non-African populations (Figures 3D-F and S7; Table S5). The ancestral population size of these populations was determined to be 20,030 (19,060–21,850), very similar to that (20,260) estimated from 1000GP. These populations started to decline 367 (167–628) kyr ago. A positive correlation was also observed between the severity of out-of-Africa bottleneck and their geographic distance to Africa. The Middle East populations had the weakest bottleneck, while the Maya, an American population, had the strongest bottleneck. Similar to 1000GP non-African populations, most HGDP-CEPH non-African populations were found to increase in size recently, except an isolated Kalash population, consistent with previous studies (Ayub et al., 2015; Bergstrom et al., 2020).

### Super Bottleneck in the Early Middle Pleistocene

The super bottleneck was directly inferred on all 10 African populations, but not on all 40 non-African populations. To investigate this observation, simulations were performed with three 1000GP demographic models, designated Bottleneck I, II, and III (Figure 4). Bottleneck I simulated the average inferred demographic history of African populations with the super bottleneck, and Bottleneck II and III simulated the demography of non-African populations without and with the super bottleneck. Both Bottleneck I and II were inferred correctly in all simulated data sets (Table S6). However, no super bottleneck was detected in Bottleneck III simulations. The super bottleneck was found to cause a population size gap between the true model and inferred demographic history after the bottleneck was relieved, suggesting a hidden effect of the super bottleneck on non-African populations. Simulations were then extended to HGDP-CEHP populations with Bottleneck models IV–VI, and similar results were obtained (Figure S8; Table S7). When simulations were performed on three artificial models (Bottleneck VII–IX) with various demographic parameters, the population size gap was still detected (Figure S9; Table S8). These results suggest a hidden effect of the super bottleneck on non-African populations.

**Figure 4.**
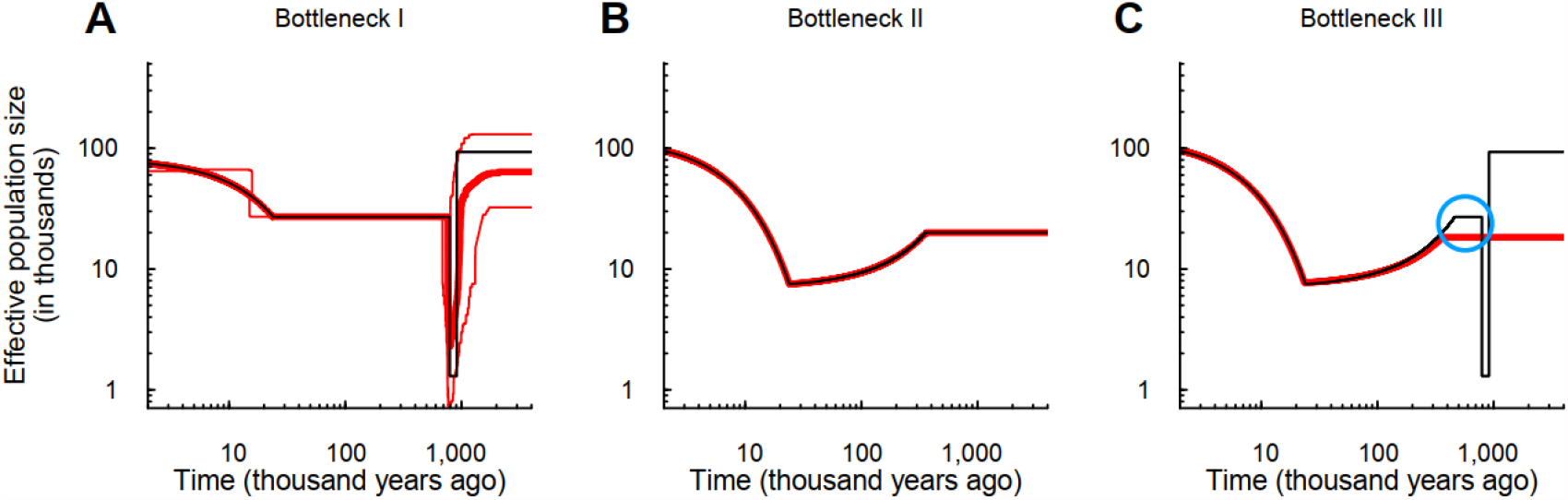
Verification of the super bottleneck. (**A**) Bottleneck I model, mimicking the demography of 1000GP African population and its estimated histories. (**B**) Bottleneck II model, mimicking the estimated demography of 1000GP non-African population and its estimated histories. (**C**) Bottleneck III model, mimicking the true demography of 1000GP non-African population and its estimated histories. Thin black lines indicate models. Thick red lines denote the medians of FitCoal estimated histories; thin red lines represent 2.5 and 97.5 percentiles of FitCoal estimated histories. Blue circle indicates the population size gap, the hidden effect of the super bottleneck in non-African populations. The mutation rate is assumed to be 1.2 × 10^−8^ per base per generation, and a generation time is assumed to be 24 years. The number of simulated sequences is 202 in Bottleneck I and 200 in Bottleneck II and III. The length of simulated sequence is 800 Mb.

The population size gap was found in both 1000GP and HGDP-CEPH data sets (Figure 3A, D). After the bottleneck was relieved, the average population sizes of non-African populations were determined to be 20,260 and 20,030, respectively, while those of African agriculturalist populations were 27,080 and 27,440, respectively in these two data sets. The observed population size gap was 7,020, probably due to the hidden effect of the super bottleneck on non-African populations.

The reasons were then investigated why the super bottleneck had different effects on African and non-African populations. Results showed that non-African populations had the out-of-Africa bottleneck, but African populations lacked such bottleneck. Therefore, the standard coalescent time of non-African populations was larger than that of African populations (Figure 3C, F). As African populations had more coalescent events occurred during the bottleneck period, the bottleneck was more readily inferred. The mathematical proof on this issue was described in the STAR★ METHODS.

## DISCUSSION

In this study, we develop FitCoal, a novel model-flexible method for demographic inference. One key characteristic feature of FitCoal is that the analytical result of expected branch length is obtained for each SFS type under arbitrary demographic models. This enables us to calculate precisely the likelihood. Second, the tabulated FitCoal is used to calculate rapidly the likelihood, making FitCoal economical of inference time. Third, the confounding effects of sequencing error and positive selection can be easily avoided by discarding rare and high-frequency mutations without losing inference accuracy. Fourth, exponential change is allowed within each inference time interval which represents a long-term continuous population change. This feature provides a better approximation to the demographic history of real populations while PSMC (Li and Durbin, 2011) and stairway plot (Liu and Fu, 2015) need multiple instantaneous changes to fit an exponential change. Last but not least, inference time intervals are variable during the demographic inference, leading to a better inference of ancient demographic events. Since coalescent events become rare when tracing backward in time, the length of time interval is usually set to increase progressively (Li and Durbin, 2011; Liu and Fu, 2015; Schiffels and Durbin, 2014; Terhorst et al., 2017). Although this strategy can capture recent demographic events, it may miss ancient ones. Therefore, FitCoal can make a fast and accurate inference for recent and ancient demographic events.

The most important discovery with FitCoal in this study is that human ancestors passed through a super bottleneck during the Mid-Pleistocene. Strikingly, the super bottleneck is inferred on all the 10 African populations while only a hidden effect of the super bottleneck is detected on all the 40 non-African populations. This observation is not only explained by the coalescent theory (see the section above) but also exclude the possibility that the super bottleneck is falsely inferred due to positive selection, population structure, sequencing error, and other confounding factors. If the inferred demographic histories of non-African populations are affected by those confounding factors, the super bottleneck should be falsely inferred on non-African populations. Moreover, large-scale simulations demonstrate that FitCoal did not falsely infer a bottleneck due to the existence of positive selection (Figure S5) and population structure (Figures S35 and S36) in African populations. Therefore, the super bottleneck exists during the Mid-Pleistocene and is shared by African and non-African populations.

The ancient population size reduction around 930 kyr ago was likely to be driven by the climatic changes at the transition between the Early and Middle Pleistocene (Lisiecki and Raymo, 2005). During the transition, low-amplitude 41 kyr obliquity-dominated glacial cycles shifted to quasi-periodic, low frequency 100 kyr periodicity, and climate change became more extreme and unpredictably associated with a longer dry period in Africa and a large faunal turnover in Africa and Eurasia (Head et al., 2008). Coinciding with this date, archaic humans referable to African *Homo erectus* became extinct. Subsequently, from about 900 until 600 kyr ago, there is a gap in the human fossil record in Africa (Figure S10) (Profico et al., 2016). Only few fossil specimens have been found in this time span, such as the cranial fragments from Gombore in Ethiopia and the mandibles from Tighenif in Algeria, all of which show features linked to later *H. heidelbergensis* representatives and represent the evolutionary origin of this species (Stringer, 2016). As a matter of fact, our data suggest that the ancestors of modern humans had a very small effective size of approximately 1,280 breeding individuals during the bottleneck period. This number is comparable in the same magnitude in the effective size of mammals threatened by extinction (Li et al., 2016).

A rapid population recovery was inferred on all 10 African populations with a 20-fold population growth during a short time period around 813 kyr ago. The earliest archaeological evidence for human control of fire was found in Israel 790 kyr ago (Goren-Inbar et al., 2004). As the control of fire profoundly affected social evolution (Foley and Gamble, 2009) and brain size (Melchionna et al., 2020), it may be associated with the big bang in population size at the end of the super bottleneck. However, climatic changes, as the alternative hypothesis, cannot be ruled out. Thus, the driving force of the rapid population recovery needs to be further studied.

The super bottleneck, which started about one million years ago, might represent a speciation event at the origin of *H. heidelbergensis* and should be strongly related to the gap in the African human fossil record. The questions about where the small ancient population dwelt, and how they survived for such a long time, remain to be investigated. Our findings may also shed light on a debate about the divergence time between Neanderthals/Denisovans and modern humans (between 440 and 270 *vs* 1,007 kyr ago) (Green et al., 2010; Ni et al., 2021; Reich et al., 2010; Shao et al., 2021). The two estimates can be verified by detecting whether ancestors of Neanderthals/Denisovans passed through the super bottleneck. In the future, a more detailed picture of human evolution during the Pleistocene may be revealed because more genomic sequences of present populations and those of archaic hominins as well as more advanced population genomics methods will be available.

## ACKNOWLEDGMENTS

We thank Daniel Zivković for sharing his codes to calculate the expected branch length, and Xiaoming Liu for sharing his simulated results. This work was supported by grants from the Strategic Priority Research Program of the Chinese Academy of Sciences (XDB13040800), the National Natural Science Foundation of China (nos. 31100273, 31172073, 91131010), and National Key Research and Development Project (No. 2020YFC0847000).

## AUTHOR CONTRIBUTIONS

W.H., Z.H., Y.H.P., and H.L. conceived and designed the research; W.H., Z.H., and H.L. wrote the code; W.H., Z.H., P.D., F.D.V., G.M., and Y.H.P. analyzed the data; W.H., Z.H., P.D., F.D.V., G.M., Y.H.P., and H.L. wrote the paper.

## DECLARATION OF INTERESTS

The authors declare no competing interests.

## STAR★METHODS

### KEY RESOURCES TABLE

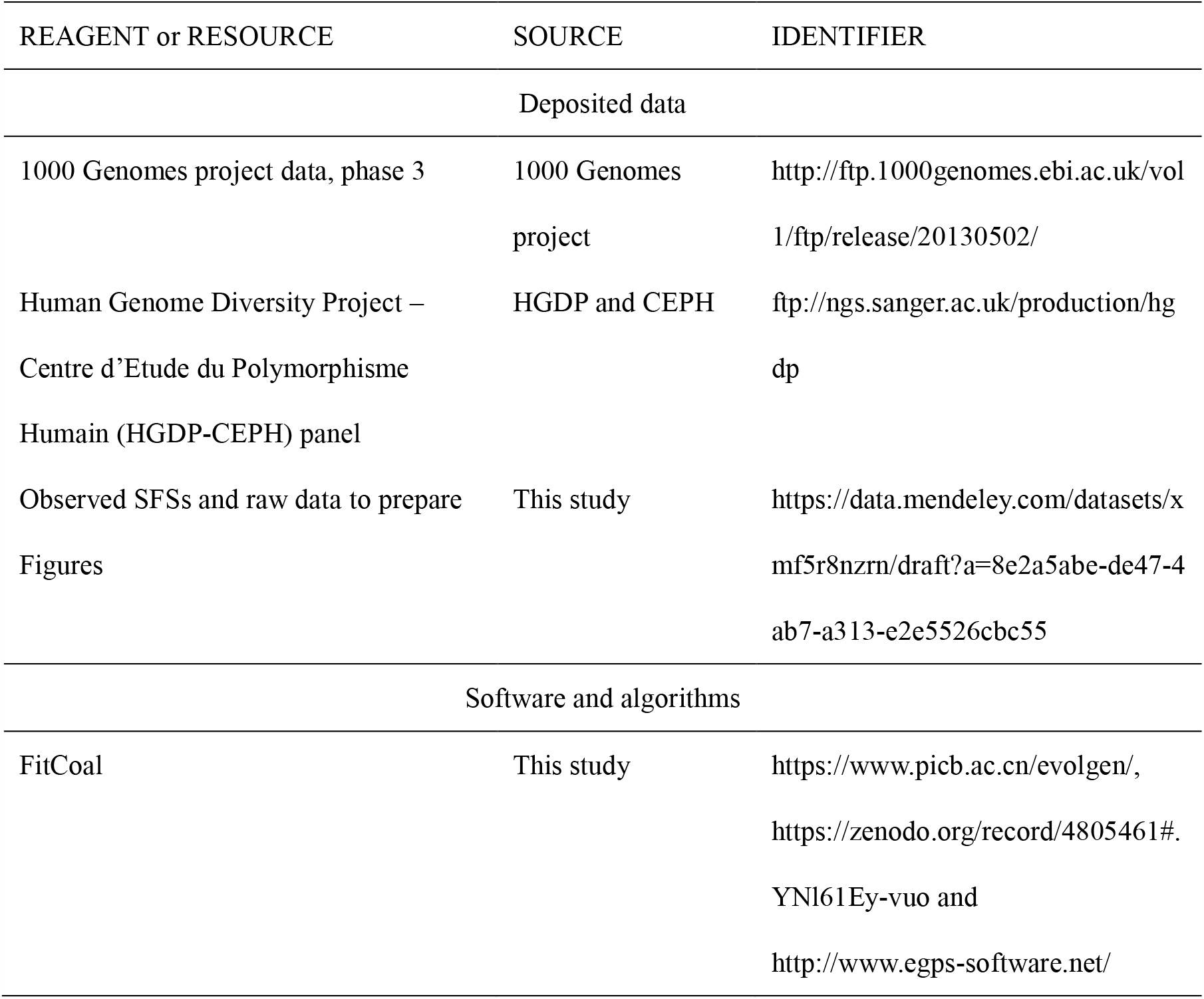

### CONTACT FOR REAGENT AND RESOURCE SHARING

Further information and requests for resource and reagents should be directed to and will be fulfilled by the Lead Contact, Haipeng Li.

## METHOD DETAILS

### Standard coalescent time and time in generations

The population size is denoted *N*(⋅), representing the demographic history. Time τ represents one-point scaled time since the time in a generation is scaled by 2*N*(0). Time *t* is usually scaled by 2*N*(*t*) generations (Bhaskar and Song, 2014; Chen, 2019; Fu, 1995; Myers et al., 2008). To distinguish it from the one-point scaled time τ, time *t* is designated as the standard coalescent time.

### Fast infinitesimal time coalescent (FitCoal) process

The FitCoal calculates the expected branch length for each type of site frequency spectrum (SFS) under arbitrary demographic history *N*(⋅). We assume that a sample is obtained by randomly taken *n* sequences from the population. The sample is designated to be state *l* (*l* = 2, ⋯, *n*) at time *t* if it has exactly *l* ancestral lineages at this time. The probability of state *l* at time *t* is denoted *p*_*l*_(*t*). In a coalescent tree, a branch is designated to be type *i* if it has exactly *i* descendants. We have

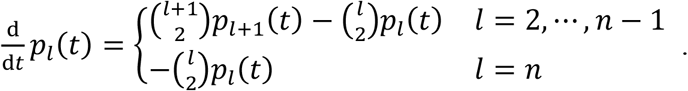

When Δ*t* is extremely small (Figure 1), there is at most one coalescent event during *t* and *t* + Δ*t*, leading to

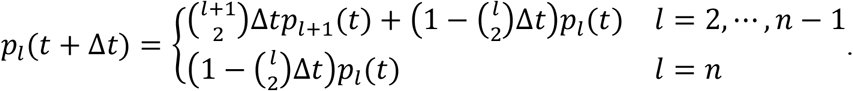

The branch length is in units of generations. The expected branch length of state *l* during *t* and *t* + Δ*t* is calculated as 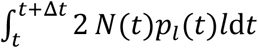. The probability that a branch of state *l* is of type *i* is 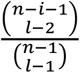 (Fu, 1995). The expected branch length of type *i* of state *l* during *t* and *t* + Δ*t* is 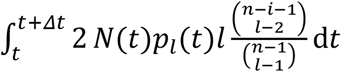. Therefore, the expected branch length *BL*_*i*_(*N*(⋅)) of type *i* is

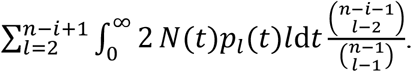

A FitCoal time partition is denoted by {*t*_0_, *t*_1_, ⋯, *t*_*m*_}, where 0 = *t*_0_ < *t*_1_ < ⋯ < *t*_*m*_. We have 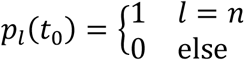. For a large positive number *m*, if *t*_*m*_ is large and (*t*_*k*_ − *t*_*k*−1_) is small for *k* = 1, ⋯, *m*, then

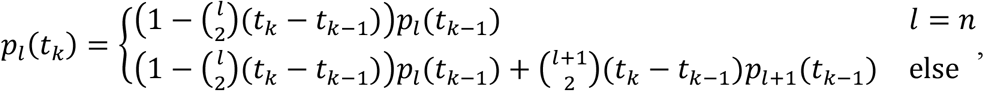

where *k* = 1, ⋯, *m*.

The expected branch length of type *i* is calculated as

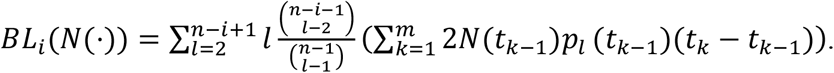

To determine the time partition, we required that the coalescent probability was less than 10^−4^ during *t*_*k*−1_ and *t*_*k*_ (*k* = 1, ⋯, *m*), the probability of common ancestor (*i*.*e*., the probability of state 1) at *t*_*m*_ was larger than (1 − 10^−6^). When the sample size was 10, the number of infinitesimal time intervals was 1,571,200. When the sample size was 200, the number of infinitesimal time intervals was 7,038,398. Thus, each Δ*t* was extremely small for precise calculation of expected branch length, and the time was partitioned to obtain *p*_*l*_(*t*) in order to calculate the expected branch length of type *i*.

### Tabulated FitCoal

The expected branch length of each type can be calculated for arbitrary time intervals according to the procedure described above. Considering another tabulated time partition {*t*_0_, *t*_1_, ⋯, *t*_*m*_} (0 = *t*_0_ < *t*_1_ < ⋯ < *t*_*m*_), the expected branch length of a type is equal to the sum of the expected branch length of this type during each tabulated time interval, thus the latter can be rescaled and tabulated.

The scaled expected branch length *BL*_*i,t*_ of type *i* during 0 and *t* is 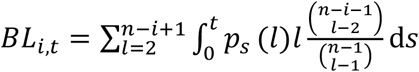, where *i* = 1, ⋯, *n* − 1. For the tabulated time partition 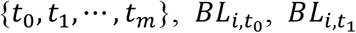, …, and 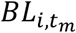 are tabulated. When *n* = 10, *m* = 231. When *n* = 200, *m* = 529.

*BL*_*i,t*_ is used to calculate the expected branch lengths under arbitrary demographic histories. When 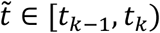,

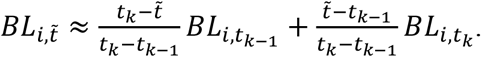

If *N*(*t*) is a piecewise constant, that is, there exists a demographic time partition 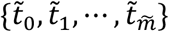, such that *N*(*t*) = *N*_*k*_ for 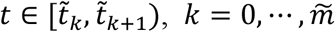. Then, the expected branch length of type *i* is calculated as

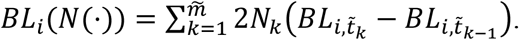

When *N*(*t*) is complex, the population size can be approximated by a piecewise constant function.

### Composite likelihood

The mutation rate per base pair per generation is denoted *μ*, and 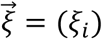 is the observed number of SNPs of *n* sequences with σ base pairs, where *i* = 1, ⋯, *n* − 1. The expected SFS is 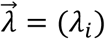, where *λ*_*i*_ = *μ*σ*BL*_*i*_(*N*(⋅)). Following the Poisson probability and previous studies (Li and Stephan, 2006), the composite likelihood is calculated as follows:

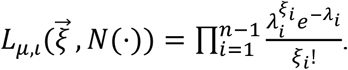

The likelihood is extended to missing data and truncated SFS (see Supplemental Text).

### Demographic inference

The number of demographic time intervals is variable. FitCoal first fits the observed SFS using a constant size model with one demographic time interval, and the number of time intervals is increased by one at a time to generate more complex models. The Local Unimodal Sampling (LUS) algorithm (Pedersen, 2010) is used to maximize the likelihood and estimate demographic parameters. A log-likelihood promotion rate is used to determine the best model to explain the observed SFS, and 20% is used as the threshold.

A series of demography with *m* pieces is denoted by a set *S*(*m*), where *S*(*m*) contains all of the following *m* pieces of population size:

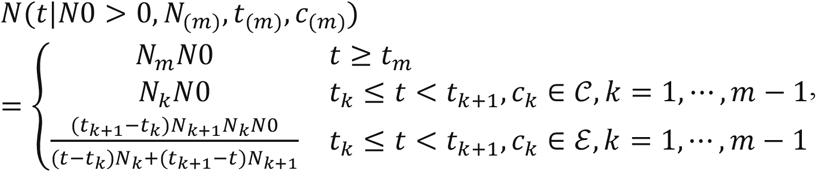

where *N*_(*m*)_ = (*N*_1_, ⋯, *N*_*m*_) ∈ *N*[*m*], *t*_(*m*)_ = (*t*_1_, ⋯, *t*_*m*_) ∈ *t*[*m*], *c*_(*m*)_ = (*c*_1_, ⋯, *c*_*m*_) ∈ *c*[*m*], *N*[*m*] = {(*N*_1_, ⋯, *N*_*m*_)|*N*_1_ = 1, *N*_*i*_ > 0 for *i* > 1}, *t*[*m*] = {(*t*_1_, ⋯, *t*_*m*_)|0 = *t*_1_ <⋅⋅⋅< *t*_*m*_}, *c*[*m*] = {(*c*_1_, ⋯, *c*_*m*_)|*c*_*m*_ ∈ 𝒞, *c*_*i*_ ∈ 𝒞 ∪ ℰ for = 1, ⋯, *m* − 1}, 𝒞 = {constant}, and ℰ = {exponential}.

The set *S*(*m*) was used as the wide-range parameter space to determine the maximum likelihood. To find the best demographic history to explain the observed SFS, the following procedures were used:

1. The number of inference time intervals (or pieces) *m* is initially set to 1, and the maximum likelihood max *L*_1_ is determined with the constant size model (model in *S*(1)).
2. Increase *m* by 1. For each change of type *c*_(*m*)_, parameters *N*_(*m*)_ = (*N*_1_, ⋯, *N*_*m*_) and *t*_(*m*)_ = (*t*_1_ = 0, *t*_2_, ⋯, *t*_*m*_) are searched to maximize the likelihood by LUS algorithm to fit the observed SFS. The maximum likelihood max *L*_*m*_ is calculated with models in S(m) with all possible change types.
3. Repeat step (2) until (1 + threshold) · log(max *L*_*m*_) < log(max *L*_*m*−1_) is obtained. The best model corresponding max *L*_*m*−1_ is determined to explain the observed SFS.
4. To avoid local optima, steps (1) – (3) are repeated *K* times to find the best model. *K* = 10 when analyzing simulated samples, and *K* = 200 when analyzing the observed SFSs of the 1000GP and HGDP-CEPH populations.

To determine the threshold of log-likelihood promotion rate, a large number of simulations were performed (Table S9). For each model, 200 replicates were conducted, and the number of inference time intervals in the estimated demographic history was determined for each replicate. If the estimated number of inference time intervals was larger than the true number of inference time intervals, overfitting was recorded. When the former was smaller than the latter, underfitting was considered. The thresholds of 10%, 20%, and 30% were used. When 10% was used, the maximum overfitting rate was 2%. When 20% was used, all cases examined were inferred correctly. When 30% was used, the underfitting was observed in one of 20 examined models. Therefore, 20% was used as the threshold of log-likelihood promotion rate in subsequent analyses.

### Data simulation

Data were simulated using ms (Hudson, 2002) and MaCS (Chen et al., 2009) software. Unless otherwise specified, a generation time was assumed to be 24 years (Liu and Fu, 2015; Scally and Durbin, 2012), the mutation rate *μ* was set for 1.2 × 10^−8^ per base per generation (Campbell et al., 2012; Conrad et al., 2011; Kong et al., 2012; Liu and Fu, 2015), and the recombination rate was *r* = 0.8*μ*. For each model, 200 SFSs were simulated to calculate the median and 2.5 and 97.5 percentiles. When verifying the inferred demographic histories, 80,000 DNA fragments with the length of 10kb each were used for simulation, taking into the consideration of small fragments split by sequencing mask in 1000GP and HGDP-CEPH data sets. High frequency alleles of SFS (10% mutation types for Bottleneck I, II, III, VII, VIII, IX, and 15% for Bottleneck IV, V, VI) were removed when assessing models to verify the super bottleneck. Detailed simulation command lines and demographic inference are presented in the Supplementary Text.

### 1000 Genomes Project data

Sequences of autosomal SNPs in 1000GP phase 3 (Altshuler et al., 2015) were downloaded from the 1000GP ftp server (ftp://ftp.1000genomes.ebi.ac.uk/vol1/ftp/release/20130502/), and 26 populations were analyzed, including seven African populations (ACB, ASW, ESN, GWD, LWK, MSL, and YRI), five European populations (CEU, FIN, GBR, IBS, and TSI), five East Asian populations (CDX, CHB, CHS, JPT, and KHV), five South Asian populations (BEB, GIH, ITU, PJL, and STU), and four American populations (CLM, MXL, PEL, and PUR). The 1000 GP strict mask was used to exclude artifacts of SNP calling. Noncoding regions except pseudogenes, defined by GENCODE release 35 (Frankish et al., 2019), were examined to avoid potential effects of purifying selection. The number of sites that passed the filtering was 826,649,529 in the human genome. Bi-allelic polymorphic sites with high-confidence ancestral allele inference, according to 1000GP annotations, were used. To avoid the effect of positive selection, high frequency mutations were excluded, and the truncated SFS was used to infer demographic history (Figure S11; Table S10). The average proportion of excluded high-frequency SNPs for all 1000GP populations was 4.40%.

### HGDP-CEPH data

In total, 24 populations were analyzed, including three African populations (Biaka, Mandeka, and Yoruba), five European populations (Adygei, Basque, French, Russian, and Sardinain), four Middle East populations (Bedouin, Druze, Mozabite, and Palestinian), three East Asian populations (Han, Japanese, and Yakut), eight Central and South Asian populations (Balochi, Brahui, Burusho, Hazara, Kalash, Makrani, Pathan, and Sindhi), and an American population (Maya). Only bi-allelic SNPs locating in GENCODE non-coding regions (Frankish et al., 2019) except pseudogenes that passed HGDP-CEPH filtering were used. HGDP-CEPH accessible mask was also used to filter SNPs (Bergstrom et al., 2020). The number of sites that passed the filtering was 791,999,125 in the human genome. Missing data were allowed to avoid artifacts due to imputation. The proportion of sites with two or more missing individuals was less than 3% for all populations (Table S11). Each population had two SFSs, with one calculated from sites with no missing data, and another from sites with one missing individual. Similarly, truncated SFSs were used to avoid the effect of positive selection (Figures S12 and S13; Table S12). The average proportion of excluded high-frequency SNPs for all HGDP-CEPH populations was 7.18%.

### SFS truncation

Denote the SFS of *n* samples by 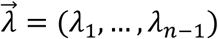. An *m*-dimension vector 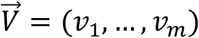 is said to be tail-up if there exist *z* ∈ {1, ⋯, *m* − 1} such that *v*_*z*_ < ⋯ < *v*_*m*_. If 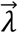 is the expected SFS of a single varying size population, we have *λ*_⌈*n*/2⌉_ > ⋯ > *λ*_*n*−1_. However, the observed SFS 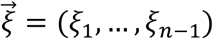 may be tailed up because of some evolutionary factors, such as positive selection and population structure, which could introduce bias to the demographic inference. Therefore, the truncated SFS is recommended.

A simple procedure is implemented to discard the tail-up types of SFS, containing high-frequency mutations. To determine the truncated tail of SFS, a small window slides through the SFS. The cutoff is determined if ξ_*i*_ exceeds its random fluctuation range. Let 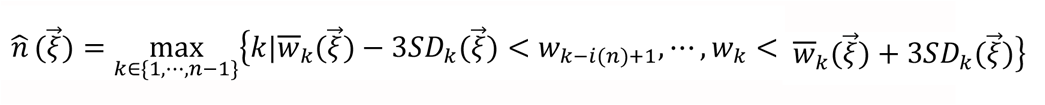, where 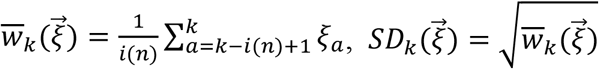, and 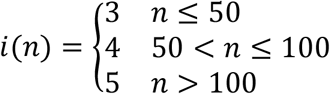. The truncated SFS 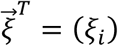, where *i* = 1, ⋯, *k*. In the analysis, we used this strategy to truncate the SFS for each human population. We call (*n* − *k*)/*n* the proportion of truncated SFS types.

When the truncating strategy was applied, the proportion of truncated SFS types was different for different populations (Table S5, S7). Therefore, to verify the effect of this strategy, the same truncating standard (∼10%, the mean proportion) was also used for 1000GP populations (Figure S15). For HGDP-CEPH, because the proportion of considered SNPs without missing samples is larger than 80% for all populations, we used the corresponding SFS to determine the cutoff to truncate both SFSs.

Similarly, the same truncating standard (∼15%, the mean proportion) was used for HGDP-CEPH (Figure S15).

### Composite likelihood

Denote *μ* as the mutation rate per base pair per generation. Denote 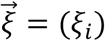 as the observed number of SNPs of *n* sequences with σ base pair, where *i* = 1, ⋯, *n* − 1. The expected SFS 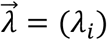, where *λ*_*i*_ = *μ*σ*BL*_*i*_(*N*(⋅)). Following the Poisson probability and the previous studies (Hudson, 2001; Li and Stephan, 2006), the composite likelihood could be written as

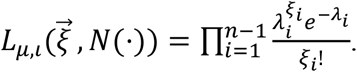

For missing data, we assume that σ^(*n*)^ base pair are sequenced in *n* samples and *S* is the set of all sample sizes. We denote the observed number of SNPs of *n*(∈ *S*) sequences by 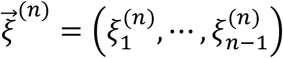. The expected SFS of *n* sequences 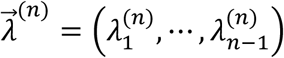, where 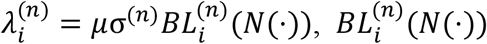 is the expected branch length of type *i* with *n* samples under population size *N*(⋅). Total number of base pair is given by σ(*S*) : = ∑_*n*∈*S*_ σ^(*n*)^. The composite likelihood could be written as

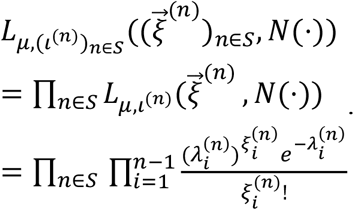

If SFS is tail-up, we use truncated SFS 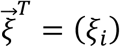, where *i* = 1, ⋯, *k*. The composite likelihood is

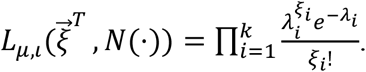

Sequencing errors often affect rare mutations in a sample. Thus singletons and mutations with size (*n* − 1) can be discarded. Although this is unnecessary in this study, as a general method, the composite likelihood of an SFS without those mutations is

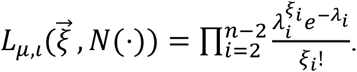

### Loss of genetic diversity due to the super bottleneck

To measure the loss of current human genetic diversity due to the super bottleneck, we calculated the expected tree length of demographic histories with or without the super bottleneck. It was straightforward to ignore a bottleneck with instantaneous size changes, thus we considered seven 1000GP African populations (ACB, ASW, ESN, GWD, LWK, MSL and YRI) and one HGDP-CEPH African population (Yoruba). To remove the bottleneck, we replaced the population size during the super bottleneck with that after the bottleneck. We then compared the expected tree length of inferred demographic history (ω_1_) with that of demographic history without the bottleneck (ω_0_).

The loss of current genetic diversity due to the super bottleneck is (ω_0_ − ω_1_)/ω_0_. When the actual sample size was used for each population, the genetic diversity was measured as Watterson’s *θ*. The genetic diversity loss of these eight populations was 46.22% and the range was 32.17–60.56%.

When *n* = 2, the genetic diversity was measured as π, the pairwise nucleotide diversity. The loss of current genetic diversity in these eight populations was 65.85% and the range was 52.71–73.60%. It was larger than the estimate based on Watterson’s *θ* because the bottleneck was ancient and the recovery rate of Watterson’s *θ* was faster than that of π (Tajima, 1989). These results demonstrate the importance of the super bottleneck in the human evolution.

## QUANTIFICATION AND STATISTICAL ANALYSES

### Validation of FitCoal calculation

We verified the calculation of expected branch lengths in this section. Under the constant size model, when the sample size was small (*n* = 5, where *n* is the number of sequences) or extremely large (*n* = 1,000), FitCoal calculated the expected branch lengths correctly (Fu, 1995) (Figure S14, Table S13). Computational accuracy reaches 10^−8^ or 10^−11^. The high accuracy is important for the precise estimation of demographic history in the following sections.

Moreover, our results were almost the same as the expected branch lengths under three simple models calculated by using the Zivković-Wiehe method (Zivković and Wiehe, 2008) (Table S14). Since Zivković-Wiehe equations can be numerically solved when n < 50, we could not compare our results with theirs when the sample size was large.

For more complex models, the average branch lengths were obtained from extensive coalescent simulations. Although with certain variances, the simulated results were consistent with the FitCoal expected branch lengths under different demographic models (Table S15). Therefore, FitCoal can analytically derive the expected branch length for each SFS type under arbitrary demographic models.

We also compared the results obtained from the tabulated FitCoal and those from the original ones without tabulation. These results were nearly identical with each other (Tables S14 and S15). Since the former was much faster than the latter, the former was used to infer demographic histories. Hereafter, tabulated FitCoal is referred to as FitCoal for short, unless otherwise indicated.

### FitCoal- and simulation-based likelihood surface

In this section, we compared two likelihood surfaces based FitCoal and simulation (Figure S1). We considered an instantaneous growth model. The population size increases from 10,000 (*N*_1_) to 20,000 (*N*_0_) at standard coalescent time 0.2. For simplicity, we obtained a SFS by multiplying the expected branch length by *θl* (= 4N_0_*μ*), where *μl* = 1.0. The number of sequences is 100.

We then compared the FitCoal composite likelihood surface of the SFS and the composite likelihood surface of the SFS based on simulation approach. To draw the likelihood surfaces, we performed a grid search in a parameter space. We considered that the population size increase from *N*_1_ to *N*_0_ at standard coalescent time 0.2, where *N*_0_ ranges from 19,600 to 20,400 and *N*_1_ from 9,800 to 10,200. The coalescent simulations were conducted by the ms software. The number of simulations is 100,000 to calculate the simulation-based likelihood.

The surface of FitCoal likelihood is smooth, but the surface of likelihood based on simulation approach is rugged (Figure S1). Moreover, the FitCoal likelihoods are also larger than those based on simulation approach because the FitCoal expected branch lengths fit the data better than the average branch lengths obtained from simulations.

### Demographic inference on simulated data

It has been shown that FitCoal can precisely estimate the demographic histories under six different demographic models (Figure 2). We then validated the accuracy of FitCoal on more simulated data in this section.

Comparing with the examined cases (Figure 2), the performance of FitCoal can be further improved by providing a priori knowledge. In some circumstances, a slow and continuous change may be more biological relevant than a quick and sudden change and vice versa. FitCoal was then re-performed conditional on either exponential or instantaneous change within each inference time interval (Figures S16 and S17). Our results showed that the FitCoal accuracy was enhanced in the presence of correct priori knowledge. Even if the condition was misspecified, the inferred demographic histories were still similar with the true histories.

FitCoal is a model-flexible method and the number of inference time intervals is dependent on the complexity of true demography. FitCoal has the power to detect more complex population histories (Figure S18). Although FitCoal may omit slight changes of population size occurred in short time periods, it has great ability to detect the major changes in all examined complex histories. When two-population split models are considered (Figure S19), FitCoal is reasonably accurate but with a slightly larger recent population size due to the effects of migration.

### Effects of positive selection

To simulate samples affected by positive selection, we considered a two-locus model (Kim and Stephan, 2002) under a constant size model. We assumed that the effective population size was 27,000, and the number of neutral fragments were 10,000, and 10 or 20% of them were partially linked with selected alleles. The distance between the neutral and the selected loci was 50kb, and recombination rate was 1cM per Mb. The sample size was 202 (the average sample size of 1000GP populations). The selection coefficient (*s* = 0.01 or 0.05) was varied. We assumed a mutation rate of 1.2 × 10^−8^ per base per generation and a generation time of 24 years. To compare among different cases, the fixed number SNPs (5,882,885 SNPs, the average number of SNPs in 1000GP populations) were applied. Under neutrality, it was equivalent to the sequenced length of 771.589 Mb.

All the simulated samples had a tail-up feature because of the excess of high-frequency mutations (Fay and Wu, 2000). Considering the low genetic diversity of selected loci, the contribution of selected loci to the genome-wide diversity was relatively low, thus only a slight excess of rare mutations (Fu and Li, 1993) was observed. The ratio between the number of singletons and doubletons ranged between 2.01 and 2.10 in the simulated samples, only slightly larger than the expected value (2.0) under neutrality.

We then applied FitCoal to estimated demography. When the full SFSs were used, our results showed that the population size remains constant within 2,000 kry (Figure S5A). If the selection strength was greatly strong (*s* = 0.05, where *s* is the selection coefficient), FitCoal estimated a large ancient population ∼240 kyr ago because of the effects of high-frequency mutations. When the high-frequency mutations were removed (*i*.*e*. the truncated SFS), the large ancient population size was reduced (Figure S5B). If *s* = 0.01 and 20% loci were subject to positive selection, a slight population expansion was observed, corresponding to the slight excess of rare mutations due to positive selection. Overall, a correct demographic history was estimated within two million years.

### Verification of inferred human demographic histories

To evaluate the precision of the inferred human demographic histories (Figure 3), we simulated 200 data sets under each demographic history. The SFSs of simulated data fit the observed SFSs perfectly (Figures S20 and S21). The results showed that FitCoal, with truncated SFS, is highly accurate to reveal human demographic history (Figures S22 – S32). Moreover, when high-frequency mutations were discarded, the truncated proportion of SFS was different for different populations. To address the influence of truncated proportions, we inferred the demographic histories by setting the average truncating proportion within each data set (10% for 1000GP and 15% for HGDP-CEPH) (Fig S10). Results were consistent with the ones obtained above. Therefore, the strategy of truncating SFS does not affect our conclusions.

Similar with the log-likelihood ratio test, the number of inference time intervals was determined by the log-likelihood promotion rate when increasing the number of inference time intervals. It is recommended to use 20% as the threshold of log-likelihood promotion rate derived from extensive simulation results (Table S11). When analyzing the human data, the inferred demographic histories are not sensitive to this threshold (Figure S33, S34; Tables S16, S17). For example, the log-likelihood promotion rate for three and four inference time intervals of CEU is 2471.16 and 17.07%, respectively. The number of inference time intervals is three, and the inferred demographic history is highly similar with that with four inference time intervals. Thus, the inferred demographic histories are robust to the threshold of 20%.

### The super bottleneck estimated in Africans

In this section, we explored why the super bottleneck can only be estimated in the African population and provided the mathematical explanation. We proved that the inferred number of intervals before time *t* depends on the dimension of the SFS before time *t*.

Denote the probability of state *l* at time *t* from *n* samples by 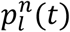, where *l* = 2, ⋯, *n*. And denote the expected brach length of size *i* from *n* samples by 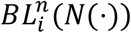, where *i* = 1, ⋯, *n* − 1. There exists an invertible matrix 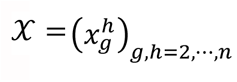 which only depends on *n*, such that 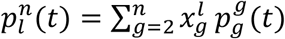 (Bhaskar and Song, 2014; Polanski et al., 2003). If positive numbers *m* < *n*, there exist a matix 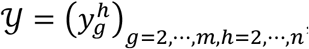, which only depends on *m* and *n*, such that 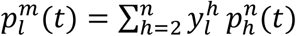. Combined with eq(1), there exist a matrix 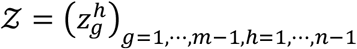, which only depends on *m* and *n*, such that 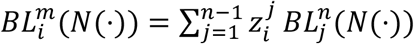.

Define the population size before time *t* by *N*^*t*^(*s*) = *N*(*t* + *s*). Denote the expected branch length of state *l* before time *t* by *B*_*l*_(*t*) = (*b*_1,*l*_(*t*), ⋯, *b*_*l*−1,*l*_(*t*)), where *b*_*i,l*_(*t*) represent the expected branch length of state *l* before time *t* of type *i* at time *t*. We have 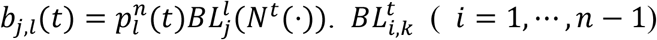 denote the branch length of type *i* whose number of lineages are no more than *k* before time *t*. We have

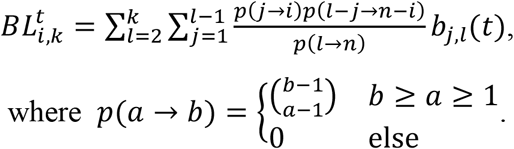

Then,

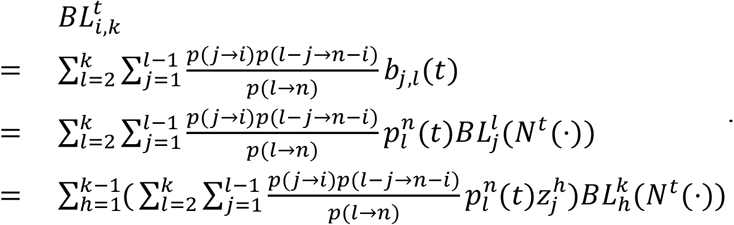

Thus, the space that is generated by 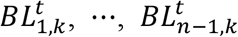 can be generated by 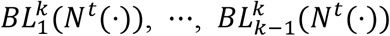. This leads that the dimension of 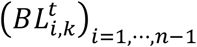 is no more than (*k* − 1).

If the number of ancestral lineages is no more than *k* before a given standard coalescent time *t*, the number of inference time intervals should be no more than (*k* − 1) before time *t* in the inferred demographic history without overfitting. Technically speaking, if a high proportion of the number of ancestral lineages is no more than *k* before a given standard coalescent time *t*, we have the same conclusion because it is an inferred demographic history.

For the non-African populations, when *t* = 1.0, the number of ancestral lineages is no more than three in more than 90% cases (Table S18), indicating the power to infer an constant size model (with one inference time interval), an expansion or contraction (with two inference time intervals) beyond this time point. The end time of the super bottleneck is 813 (772–864) kyr ago and the corresponding standard coalescent time is larger than 1.0 for all non-African populations (Figure 3C, F). Therefore, the super bottleneck cannot be inferred in this case since the bottleneck contains three inference time intervals.

### Confounding factors of bottleneck

African populations have complex population structure (Hsieh et al., 2016; Lopez et al., 2018; Schlebusch and Jakobsson, 2018; Skoglund et al., 2017), and a complex population structure model is proposed for African and European populations (Lopez et al., 2018) (Figure S35). To address the effects of population structure, we simulated data for a western rainforest hunter-gatherer (wRHG) and a western farmer (wARG) population and estimated their demographic histories (Figure S35). Due to frequent migrations, a larger recent population size is estimated for both populations. However, the ancient population size (14,427) is accurately inferred for both populations (14,493 and 14,428). Thus, the super bottleneck is not due to the complex African population structure.

To consider the effects of archaic introgression from ghost populations (Beerli, 2004; Durvasula and Sankararaman, 2020), we examined different models by assuming that introgression happened in different time periods with different migration rates (Figure S36). Results show that archaic introgression does not result in an ancient super bottleneck.

Truncated SFS was used in demography inference in this study. To examine the effects of SFS truncation, the FitCoal inference was re-performed by taking the full SFSs that include high-frequency derived mutations. Again, the super bottleneck is revealed only in the African populations, but not in the non-African populations (Figures S37 and S38). Therefore, the ancient super bottleneck is not due to the effects of SFS truncation.

### Computational performance

We compared the performance of the FitCoal with or without tabulation. We applied them to analyze the data of YRI population by fixing four inference time intervals and allowing instantaneous population size change. The former is much faster than the latter (1 second *vs* 36.2 hours).

## DATA AND SOFTWARE AVAILABILITY

The authors declare that all data are available in the main text and the supplementary materials. FitCoal is a free plug-in of the eGPS software (Yu et al., 2019) and can be downloaded and run as an independent package. FitCoal and its documentation are available via Zenodo at https://zenodo.org/record/4805461#.YNl61Ey-vuo, our institute website at http://www.picb.ac.cn/evolgen/, and eGPS website http://www.egps-software.net/. Raw data were deposited on Mendeley (https://data.mendeley.com/datasets/xmf5r8nzrn/draft?a=8e2a5abe-de47-4ab7-a313-e2e5526cbc55).

## Notes

https://data.mendeley.com/datasets/xmf5r8nzrn/draft?a=8e2a5abe-de47-4ab7-a313-e2e5526cbc55

### Competing Interest Statement

The authors have declared no competing interest.

### Summary of Updates

The abstract, introduction, and discussion has been re-written to make our findings clearer. And the external raw data is also provided.

