## Supplementary Materials for "Genomic inference of a human super bottleneck in Mid-Pleistocene transition"

**List of supplementary figures**

Figure S1. Comparison of likelihood surfaces based on simulations and FitCoal.

Figure S2. Effects of sequence length (A), sample size (B), and recombination rate (C) in the FitCoal inference.

Figure S3. Verification of FitCoal accuracy with truncated SFS.

Figure S4. Verification of FitCoal accuracy with truncated SFS under more complexed models.

Figure S5. Effects of positive selection on demographic inference.

Figure S6. Inferred demographic histories and standard coalescent times of 1000GP populations.

Figure S7. Inferred demographic histories and standard coalescent times of HGPD-CEPH populations.

Figure S8. Verification of the HGDP-CEPH inferred super bottleneck.

Figure S9. Verification of the super bottleneck in artificial models.

Figure S10. Distribution of the human fossil record in Africa from the late Early Pleistocene to present.

Figure S11. The observed SFSs of 1000GP populations.

Figure S12. The observed SFSs without missing data of HGDP-CEPH populations.

Figure S13. The observed SFSs with one missing individual of HGDP-CEPH populations.

Figure S14. Comparison of the expected branch lengths of FitCoal and the average branch lengths of coalescent simulations.

Figure S15. Inferred demographic histories of 1000GP and HGPD-CEPH populations using the same truncating SFS standard for each data set.

Figure S16. Estimated demographic histories of FitCoal conditional on exponential change, stairway plot, and PSMC using simulated samples.

Figure S17. Estimated demographic histories of FitCoal conditional on instantaneous change, stairway plot, and PSMC using simulated samples.

Figure S18. Verification of the accuracy of FitCoal using simulated samples under complex models.

Figure S19. Verification of FitCoal accuracy under three migration models.

Figure S20. The observed SFS and simulated SFS of 1000GP populations.

Figure S21. The observed SFS without missing data and simulated SFS of HGDP-CEPH populations.

Figure S22. 95% confidence intervals of 1000GP African populations.

Figure S23. 95% confidence intervals of 1000GP European populations.

Figure S24. 95% confidence intervals of 1000GP East Asian populations.

Figure S25. 95% confidence intervals of 1000GP South Asian populations.

Figure S26. 95% confidence intervals of 1000GP American populations.

Figure S27. 95% confidence intervals of HGPD-CEPH African populations.

Figure S28. 95% confidence intervals of HGPD-CEPH Middle East populations.

Figure S29. 95% confidence intervals of HGPD-CEPH European populations.

Figure S30. 95% confidence intervals of HGPD-CEPH East Asian populations.

Figure S31. 95% confidence intervals of HGPD-CEPH Central & South Asian populations.

Figure S32. 95% confidence intervals of HGPD-CEPH American population.

Figure S33. Inferred demographic histories with different inference time intervals of 1000GP populations.

Figure S34. Inferred demographic histories with different inference time intervals of HGPD-CEPH American populations.

Figure S35. Verification of inference accuracy under complex population structure model.

Figure S36. Verification of inference accuracy when migration occurred with unknown hominin population.

Figure S37. Estimated demographic histories using the full and the truncated SFSs of 1000GP populations.

Figure S38. Estimated demographic histories using the full and the truncated SFSs of HGDP-CEPH populations.

**List of supplementary tables**

Table S1. Proportion of correctly-inferred change type for the most recent demographic event in six models.

Table S2. Parameters of the super bottleneck in 1000GP African populations.

Table S3. Parameters of the super bottleneck in HGDP-CEPH African populations

Table S4. Parameters of the out-of-Africa bottleneck in 1000GP non-African populations.

Table S5. Parameters of the out-of-Africa bottleneck in HGDP-CEPH non-African populations.

Table S6. The super bottleneck parameters of Bottleneck I model.

Table S7. The super bottleneck parameters of Bottleneck IV model.

Table S8. Super bottleneck parameters of Bottleneck VII model.

Table S9. Influence of different log-likelihood promotion thresholds.

Table S10. Information of truncated SFS of 1000GP populations.

Table S11. Proportion of SNPs without or with missing data of HGDP-CEPH populations.

Table S12. Information of truncated SFS of HGDP-CEPH populations.

Table S13. Comparison of branch length between theoretical values and FitCoal under the constant size model.

Table S14. Comparison of accuracy between FitCoal and Z-W's method.

Table S15. Comparison of accuracy between FitCoal and simulations.

Table S16. Likelihood promotion rate of inferred demographic histories with different inference time intervals of 1000GP populations.

Table S17. Likelihood promotion rate of inferred demographic histories with different inference time intervals of HGDP-CEPH populations.

Table S18. Probabilities of each state (the number of ancestral lineages remained) at time *t*.

### Paleoanthropology

The fossil record of human evolution is rather rich in some passages, whereas it is poor during others. Particularly, at the transition between the Early and the Middle Pleistocene (between 900 and 600 kyr ago, corresponding to the isotope stages MIS 23-16), the human fossil record in Africa become extremely poor or even absent (Fig. S10) (Mounier et al., 2011). The species *Homo ergaster* (or African *H. erectus*) disappears abruptly around 950 kyr ago and there are no longer specimens such as OH 12 (in Tanzania), Olorgesailie (in Kenya), Daka (in Ethiopia) and Buia (in Eritrea), all associated with Mode 2 (Early Stone Age) lithic assemblages (Mori et al., 2020). Fossil evidence becomes relatively abundant again after 600 kyr ago, with fossils – such as Bodo (Ethiopia), Saldanha (South Africa) and, later, Broken Hill (Zambia) – having derived cranial features, such as a more arched frontal profile, a typical supra-orbital torus morphology and enlarged brain volumes of about 1200 ml (the average for *H. erectus* sensu lato is about 1000 ml), also these specimens are associated to Mode 2 stone tools. At the same time humans with similar characteristics, have already spread throughout Africa, as well as in Eurasia, and will be related (in the late Middle Pleistocene) to the origin of our species *H. sapiens*, as well as to those of the Neanderthals and the so-called Denisovans. These new humans are referred by many authors to the species *H. heidelbergensis*.

From an archaeological point of view, a discontinuity in the production of Mode 2 artefacts has been noted in different layers of the Melka Kunture formation in Ethiopia, dating from 1.0-0.85 Ma (Gallotti and Mussi, 2017). Furthermore, according to Shea (Shea, 2020) only a limited number (6 on 27) of archaeological sites of the later Early Stone Age in East Africa overlap the time span in which the super bottleneck occurred (see Table. 5.2 in the reference (Shea, 2020)).

The origin of *H. heidelbergensis* is still poorly known and widely debated. However, the evidence of human presence do not disappear completely from Africa during the fossil gap, although this is reduced to a few archaeological findings and an extremely limited number of fragmentary fossil remains, which point to a possible event of speciation linked to the origin of *H. heidelbergensis* in this time span, ranging between 950 and 700 kyr ago. This fragmentary fossil sample are two cranial fragments, probably belonging to the same individual, from the Gombore II site in the Melka Kunture area of Ethiopia, (about 850 kyr ago), and three isolate mandibles from the Algerian site of Tighenif or Ternifine (more than 700 kyr ago). These specimens were long attributed to the variability of *H. erectus*, but more recent analyses have denied this attribution and indicated affinities with later *H. heidelbergensis* representatives (Profico et al., 2016; Zanolli and Mazurier, 2013). In particular, in the case of Tighenif, the endostructural analysis of the teeth and the enamel-dentine junction revealed derived features linking them with the mandible of Mauer (the holotype of *H. heidelbergensis*). Furthermore, the morphology of the Tighenif mandibles do not show affinities with that of archaic contemporary specimens such as the mandibles from the Gran Dolina sites attributed to *H. antecessor* (Bermúdez de Castro et al., 2007).

**Models and simulation commands for Figure S14**

We simulated trees for 10 sequences with 1,000,000 replications using the ms software (Hudson, 2002) to calculate the average branch length for each SFS type. In this section, time is scaled by $4N_0$ generations and the population size is scaled by $N_0$ when using the ms software.

1. Constant size model (Figure S14A)

The corresponding ms command is

ms 10 1000000 -L -T

2. Exponential growth model (Figure S14B)

We assumed an ancient population size of 0.5, and the population began an exponential growth at time of 0.17328679513998632 to the current size of 1.0. The corresponding ms command is

ms 10 1000000 -G 4.0 -eN 0.17328679513998632 0.5 -L -T

3. Bottleneck model (Figure S14C)

We assumed a population size of 1.0 and a population size of 0.5 during time of 0.1 and 0.25. The corresponding ms command is

ms 10 1000000 -eN 0.1 0.5 -eN 0.25 1.0 -L -T

4. Complex model (Figure S14D)

We assumed an ancient population size of 0.5. The population experienced a bottleneck with population size decreasing to 0.2 from time of 0.11465735902799726 to 0.08465735902799726 and growing exponentially to the current size of 1.0. The corresponding ms command for the simulation is

ms 10 1000000 -G 20.0 -eN 0.03465735902799726 0.5 -eN 0.08465735902799726 0.2 -eN 0.11465735902799726 0.5 -L –T

**Models and simulation commands for Table S15**

To ensure the accuracy of FitCoal, we compared the expected branch length of FitCoal ($n = 10$) with Zivković-Wiehe's method (Zivković and Wiehe, 2008) that is a numerical calculation method for piecewise constant model with at most three inference time intervals. In this section, time is scaled by $4N_0$ generations and the population size is scaled by $N_0$ because of the requirement of Zivković-Wiehe's method.

1. Constant size model

The corresponding input parameters of Zivković-Wiehe's method are

unfoldedfreq[10, 0.0, 0.0, 1.0, 1.0]

2. Instantaneous growth model

We assumed a current population size of 1.0 and an ancient population size of 0.5 before time 0.5. The corresponding input parameters of Zivković-Wiehe's method are

unfoldedfreq[10, 0.5, 0.0, 0.5, 0.5]

3. Bottleneck model

We assumed a population size of 1.0 and a population size of 0.5 during time of 0.1 and 0.25. The corresponding input parameters of Zivković-Wiehe's method are

unfoldedfreq[10, 0.2, 0.6, 0.5, 1]

**Models and simulation commands for verifying the accuracy of FitCoal**

If not specified, we used a default mutation rate $\mu$ of $1.2 \times 10^{-8}$ per base per generation and a recombination rate $r = 0.8\mu$. All models were simulated with 200 replications using ms (Hudson, 2002) or MaCS (Chen et al., 2009). In this section, time is scaled by $4N_0$ generations and the population size is scaled by $N_0$ because of the requirement of simulation softwares.

1. Constant size model (Figure 2A, S3A, S16A and S17A)

We assumed the effective population size of 10,000. And 30 sequences of 10 Mb were simulated. The corresponding ms command is

ms 30 200 -t 4800 -r 3800 10000000

2. Instantaneous increase model (Figure 2B, S3B, S16B and S17B)

We used this model to mimic the demography of an African population (Boyko et al., 2008). We assumed the ancient population size of 7,778 and the current population size of 25,636 and the population began an instantaneous increase at 6,809 generations ago. 30 sequences of 10 Mb were simulated. The corresponding ms command for the simulation is

ms 30 200 -t 12310 -r 9750 10000000 -eN 0.066 0.3

3. PSMC "standard" model (Figure 2C, S3C, S16C and S17C)

This model was based on the "standard simulation" model in PSMC publication (Li and Durbin, 2011). 170 sequences of 30Mb were simulated. The corresponding MaCS command for this simulation is

macs 170 30000000 -i 200 -h 1e3 -t 0.002732 -r 0.002179 -h 1e3 -eN 0.01 0.05 -eN 0.0375 0.5 -eN 1.25 1

4. Exponential growth model I (Figure 2D, S3D, S16D and S17D)

We assumed the current population size of 50,000 and a growth rate $r = 0.004$. 30 sequences of 10 Mb were simulated. The corresponding ms command for the simulation is

ms 30 200 -t 24000 -r 19000 10000000 -G 800

5. Exponential growth model II (Figure 2E, S3E, S16E and S17E)

We used this model to mimic the demography of a European populations (Gutenkunst et al., 2009). We assumed the ancient population size of 1,000 and the current population size of 29,525 and the population began exponential growth at 848 generations ago. 30 sequences of 10 Mb were simulated. The corresponding ms command for the simulation is

ms 30 200 -t 14172 -r 10629 10000000 -G 472.4 -eN 0.00718 0.0339

6. Exponential growth model III (Figure 2F, S3F, S16F and S17F)

We assumed the ancient population size of 8,000. The population experienced an instantaneous decrease to 7,900 and then began exponentially growth to 900,000. 30 sequences of 10 Mb were simulated. The corresponding ms command for the simulation is

ms 30 200 -t 432000 -r 340000 10000000 -G 46368 -eN 0.0001027 0.008889

7. PSMC “sim-YH” model (Figure S4A and S18A)

This model was based on the “sim-YH” model in the PSMC publication (Li and Durbin, 2011). 170 sequences of 30 Mb were simulated. The corresponding MaCS command for the simulation is

macs 170 30000000 -i 200 -h 1e3 -t 0.002171 -r 0.001731 -eN 0.0055 0.0832 -eN 0.0089 0.0489 -eN 0.0130 0.0607 -eN 0.0177 0.1072 -eN 0.0233 0.2093 -eN 0.0299 0.3630 -eN 0.0375 0.5041 -eN 0.0465 0.5870 -eN 0.0571 0.6343 -eN 0.0695 0.6138 -eN 0.0840 0.5292 -eN 0.1010 0.4409 -eN 0.1210 0.3749 -eN 0.1444 0.3313 -eN 0.1718 0.3066 -eN 0.2040 0.2952 -eN 0.2418 0.2915 -eN 0.2860 0.2950 -eN 0.3379 0.3103 -eN 0.3988 0.3458 -eN 0.4701 0.4109 -eN 0.5538 0.5048 -eN 0.6520 0.6520 -eN 0.7671 0.6440 -eN 0.9020 0.6178 -eN 1.0603 0.5345 -eN 1.4635 1.7931

8. PSMC “sim-1” model (Figure S4B and S18B)

This model was based on the “sim-1” model in the PSMC publication (Li and Durbin, 2011). 170 sequences of 30 Mb were simulated. The corresponding MaCS command for the simulation is

macs 170 30000000 -i 200 -h 1e3 -t 0.001 -r 0.0008 -eN 0.01 0.1 -eN 0.06 1 -eN 0.2 0.5 -eN 1 1 -eN 2 2

9. PSMC “sim-2” model (Figure S4C and S18C)

This model was based on the “sim-1” model in the PSMC publication (Li and Durbin, 2011). 170 sequences of 30 Mb were simulated. The corresponding MaCS command for the simulation is

macs 170 30000000 -i 200 -h 1e3 -t 0.0001 -r 0.00008 -eN 0.1 5 -eN 0.6 20 -eN 2 5 -eN 10 10 -eN 20 5

10. PSMC “sim-3” model (Figure S4D and S18D)

This model was based on the "sim-1" model in the PSMC publication (Li and Durbin, 2011). 170 sequences of 30 Mb were simulated. The corresponding MaCS command for the simulation is

macs 170 30000000 -i 200 -h 1e3 -t 0.002 -r 0.0016 -eN 0.01 0.05 -eN 0.0150 0.5 -eN 0.05 0.25 -eN 0.5 0.5

11. Complicated model III (Figure S4E and S18E)

We assumed the ancient population size of 4,167. The population experienced an instantaneous increase to 20,833 at 33,333 generations ago, an instantaneous decrease to 2,083 at 2,500 generations ago and an instantaneous increase to 41,667 at 833 generations ago. 170 sequences of 30 Mb were simulated. The corresponding MaCS command for the simulation is

macs 170 30000000 -i 200 -h 1e3 -t 0.002 -r 0.0016 -eN 0.005 0.05 -eN 0.0150 0.5 -eN 0.2 0.1

12. Complicated model II (Figure S4F and S18F)

We assumed the ancient population size of 1,250. The population experienced an instantaneous increase to 33,333 at 33,333 generations ago, an instantaneous decrease to 12,500 at 16,667 generations ago, an instantaneous increase to 20,833 at 8,333 generations ago, an instantaneous decrease to 8,333 at 4,167 generations ago, an instantaneous decrease to 2,083 at 1,667 generations ago and an instantaneous increase to 41,667 at 833 generations ago. 170 sequences of 30 Mb were simulated. The corresponding MaCS command for the simulation is

macs 170 30000000 -i 200 -h 1e3 -t 0.002 -r 0.0016 -eN 0.005 0.05 -eN 0.01 0.2 -eN 0.0250 0.5 -eN 0.05 0.3 -eN 0.1 0.8 -eN 0.2 0.03

13. Exponential growth model IV (Figure S4G and S18G)

We assumed the ancient population size of 20,000. The population experienced an instantaneous decrease to 1,000 at 4,000 generations ago, and began exponential growth to 20,000 at 2,000 generations ago. 170 sequences of 30 Mb were simulated. The corresponding MaCS command for the simulation is

macs 170 30000000 -i 200 -h 1e3 -t 0.001 -r 0.0008 -G 120 -eG 0.025 0 -eN 0.05 1

14. Exponential growth model V (Figure S4H and S18H

We assumed the ancient population size of 15,000. The population experienced an instantaneous decrease to 6,000 at 4,000 generations ago, and began exponential growth to 30,000 at 2,000 generations ago. 170 sequences of 30 Mb were simulated. The corresponding MaCS command for the simulation is

```
macs 170 30000000 -i 200 -h 1e3 -t 0.00144 -r 0.00115 -G 96.37 -eG 0.0167 0 -eN 0.033 0.5
```

15. Exponential growth model VI (Figure S4I and S18I)

We assumed the ancient population size of 15,000. The population experienced an exponential decrease to 6,000 at 5,000 generations ago, and began exponential growth to 30,000 at 2,000 generations ago. 170 sequences of 30 Mb were simulated. The corresponding ms command for the simulation is

```
ms 170 100 -t 1440 -r 1152 1000000 -G 96.56627474604602 -eG 0.01666666666667 -36.65162927 -eN 0.0416666666667 0.5
```

16. Split model I (Figure S19A)

We used this model to mimic the split demography of African hunter-gatherer and agriculturist populations. We assumed the ancient population size of 20,833. The ancient population splits into two subpopulation at 6,667 generations ago. Population 1 experienced an instantaneous increase to 41,667 at 500 generations ago, and population 2 experienced an instantaneous decline to 8,333 at 1,250 generations ago. 170 sequences of 30 Mb were simulated. The corresponding MaCS command for the simulation is

```
macs 340 30000000 -i 1 -h 1e3 -t 0.002 -r 0.0016 -I 2 170 170 4 -n 2 0.2 -en 0.003 1 0.5 -en 0.0075 2 0.5 -ej 0.04 2 1
```

17. Split model II (Figure S19B)

We used this model to mimic the split demography of African and European populations. We assumed the ancient population size of 20,833. The ancient population split into two subpopulation at 5,000 generations ago and the population size of population 1 became 83,303. Then, population 1 experienced an instantaneous increase to 20,833 at 833 generations ago. Population 2 experienced an instantaneous increase to 41,667 at 500 generations ago. 170 sequences of 30 Mb were simulated. The corresponding MaCS command for the simulation is macs 340 30000000 -i 1 -h 1e3 -t 0.004 -r 0.0032 -I 2 170 170 4 -n 2 0.5 -en 0.0015 2 0.25 -en 0.0025 1 0.1 -ej 0.015 1 2

18. Split model III (Figure S19C)

We assumed the ancient population size of 4,167. The ancient population experienced an instantaneous growth to 20,833 at 8,333 generations ago and splits into two subpopulations at 2,500 generations ago. The population size of population 1 decreased to 2,083 at 1,666 generations ago and increased to 20,833 at 833 generations ago. The population size of population 2 remained constant. 170 sequences of 100 Mb were simulated. The corresponding MaCS command for the simulation is

macs 340 1000000 -i 100 -h 1e3 -t 0.001 -r 0.0008 -I 2 170 170 4 -eN 0 1 -en 0.01 1 0.1 -en 0.02 1 1 -ej 0.03 2 1 -eN 0.10 0.2 -eN 1 1

19. Models for studying influence factors of FitCoal (Figure S2)

We simulated variations of the Exponential growth model V by increasing or decreasing the number of sequences, length of sequences and recombination rate.

19.1. Number of sequence $n$ = 170, sequence length $L$ = 1 Mb, recombination rate $\rho = 0.8\mu$ (Figure S2A). The corresponding ms command for the simulation is

ms 170 1 -t 1440 -r 1152 1000000 -G 96.56627474604602 -eN 0.016666666666666666 0.2 -eN 0.03333333333333333 0.5

19.2. Number of sequence $n$ = 170, sequence length $L$ = 10 Mb, recombination rate $\rho = 0.8\mu$ (Figure S2B). The corresponding ms command for the simulation is

ms 170 10 -t 1440 -r 11520 1000000 -G 96.56627474604602 -eN 0.016666666666666666 0.2 -eN 0.03333333333333333 0.5

19.3. Number of sequence $n$ = 170, sequence length $L$ = 100 Mb, recombination rate $\rho = 0.8\mu$ (Figure S2C, S2F and S2H). The corresponding ms command for the simulation is

ms 170 100 -t 14400 -r 11520 10000000 -G 96.56627474604602 -eN 0.016666666666666666 0.2 -eN 0.03333333333333333 0.5

19.4. Number of sequence $n$ = 10, sequence length $L$ = 100 Mb, recombination rate $\rho = 0.8\mu$ (Figure S2D). The corresponding ms command for the simulation is

ms 10 100 -t 14400 -r 11520 10000000 -G 96.56627474604602 -eN 0.016666666666666666 0.2 -eN 0.03333333333333333 0.5

19.5. Number of sequence $n$ = 100, sequence length $L$ = 100 Mb, recombination rate $\rho = 0.8\mu$ (Figure S2E). The corresponding ms command for the simulation is

ms 100 100 -t 14400 -r 11520 10000000 -G 96.56627474604602 -eN 0.016666666666666666 0.2 -eN 0.03333333333333333 0.5

19.6. Number of sequence $n$ = 170, sequence length $L$ = 100 Mb, recombination rate $\rho = 0.1\mu$ (Figure S2G). The corresponding ms command for the simulation is

ms 170 100 -t 14400 -r 1440 10000000 -G 96.56627474604602 -eN 0.016666666666666666 0.2 -eN 0.03333333333333333 0.5

19.7. Number of sequence $n$ = 170, sequence length $L$ = 100 Mb, recombination rate $\rho = 10\mu$ (Figure S2I). The corresponding ms command for the simulation is

ms 170 100 -t 14400 -r 144000 10000000 -G 96.56627474604602 -eN 0.016666666666666666 0.2 -eN 0.03333333333333333 0.5

20. Models for studying the super bottleneck

For 1000GP, we used recent population expansion parameters of YRI and CHB to represent the recent population expansion for African and non-African population, respectively. Then, we used approximate average value of parameters inferred from African and non-African populations to represent other parameters of corresponding models.

For HGDP-CEPH, we used recent population expansion parameters of Yoruba and Han to represent the recent population expansion for African and non-African population, respectively. Then, we used approximate average value of parameters inferred from African except Biaka and non-African populations to represent other parameters of corresponding models. Because of the difference of ancient demography between Biaka and other African populations, we did not take it into consideration.

For idealized models, we fine-tuned parameters so that all parameters had a standard coalescent time smaller than 2.0.

20.1. Bottleneck I model for 1000GP (Figure 4A)

We used this model to mimic the demography of African populations. We assumed the ancient population size of 96,000. The population experienced an instantaneous decrease to 1,600 at 39,000 generations ago, an instantaneous growth to 27,000 at 32,500 generations ago and began exponential growth to 160,000 at 200 generations ago. 202

sequences of 800 Mb were simulated. The corresponding ms command for the simulation is

ms 202 80000 -t 76.8 -r 61.43 10000 -G 5693.878237534393 -eN 3.125E-4 0.16875 -eN 0.05078125 0.01 -eN 0.0609375 0.6

20.2. Bottleneck II model for 1000GP (Figure 4B)

We used this model to mimic the estimated demography of non-African populations. We assumed the ancient population size of 20,000. The population experienced an exponential decrease to 6,000 at 20,000 generations ago and began exponential growth to 250,000 at 1,000 generations ago. 202 sequences of 800 Mb were simulated. The corresponding ms command for the simulation is

ms 200 80000 -t 120 -r 96 10000 -G 3729.7014486341914 -eN 0.001 0.024 -eG 0.001 -63.36698970136505 -eN 0.02 0.08

20.3. Bottleneck III model for 1000GP (Figure 4C)

We used this model to mimic the real demography of non-African populations. We assumed the ancient population size of 96,000. The population experienced an instantaneous decrease to 1,600 at 39,000 generations ago, an instantaneous growth to 27,000 at 32,500 generations ago. Then, the population began exponential decline to 6,000 at 24,735 generations ago and instantaneously increased to 250,000 at 1,000 generations ago. 200 sequences of 800 Mb were simulated. The corresponding ms command for the simulation is

ms 200 80000 -t 120 -r 96 10000 -G 3729.7014486341914 -eN 0.001 0.024 -eG 0.001 -63.36959750479352 -eN 0.024735 0.108 -eN 0.0325 0.0064 -eN 0.039 0.384

20.4. Bottleneck IV model for HGDP-CEPH (Figure S8A)

We used this model to mimic the demography of African populations. We assumed the ancient population size of 120,000. The population experienced an instantaneous decrease to 1,400 at 41,000 generations ago, an instantaneous growth to 28,000 at 35,000 generations ago and began instantaneous growth to 50,000 at 500 generations ago. 44 sequences of 800 Mb were simulated. The corresponding ms command for the simulation is

ms 44 80000 -t 24 -r 19.19 10000 -eN 0.0025 0.56 -eN 0.17500000000000002 0.028 -eN 0.205 2.4

#### 20.5. Bottleneck V model for HGDP-CEPH (Figure S8B)

We used this model to mimic the estimated demography of non-African populations. We assumed the ancient population size of 21,000. The population experienced an exponential decrease to 6,000 at 20,000 generations ago and began exponential growth to 300,000 at 1,000 generations ago. 56 sequences of 800 Mb were simulated. The corresponding ms command for the simulation is

ms 56 80000 -t 144 -r 115.19 10000 -G 4694.427606513776 -eN 8.333333333333333E-4 0.02 -eG 8.333333333333333E-4 -79.1218716944443 -eN 0.016666666666666666 0.07

#### 20.6. Bottleneck VI model for HGDP-CEPH (Figure S8C)

We used this model to mimic the real demography of non-African populations. We assumed the ancient population size of 120,000. The population experienced an instantaneous decrease to 1,400 at 41,000 generations ago, an instantaneous growth to 25,000 at 35,000 generations ago. Then, the population began exponential decline to 6,000 at 24,363 generations ago and instantaneously increased to 300,000 at 1,000 generations ago. 56 sequences of 800 Mb were simulated. The corresponding ms command for the simulation is

ms 56 80000 -t 144 -r 115.19 10000 -G 4694.427606513776 -eN 8.333333333333333E-4 0.02 -eG 8.333333333333333E-4 -79.12228948065652 -eN 0.0203025 0.09333333333333334 -eN 0.029166666666666667 0.004666666666666667 -eN 0.034166666666666665 0.4

#### 20.7. Bottleneck VII model for idealized models (Figure S9A)

We used this model to mimic the demography of African populations. We assumed the ancient population size of 30,000. The population experienced an instantaneous decrease to 3,000 at 35,000 generations ago, an instantaneous growth to 30,000 at 30,000 generations ago. 170 sequences of 800 Mb were simulated. The corresponding ms (Hudson, 2002) command for the simulation is

ms 170 800 -t 1440 -r 1152 1000000 -eN 0.25 0.1 -eN 0.291667 1

#### 20.8. Bottleneck VIII model for idealized models (Figure S9B)

We used this model to mimic the estimated demography of non-African populations. We assumed the ancient population size of 20,000. The population experienced an exponential decrease to 6,000 at 20,000 generations ago and began exponential growth to

300,000 at 1,000 generations ago. 170 sequences of 800 Mb were simulated. The corresponding ms command for the simulation is

ms 170 800 -t 14400 -r 11520 1000000 -G 4694.427606513776 -eN 0.00083333 0.02 -eG 0.00083333 -101.6487102589958 -eN 0.016666666666666666 0.1

#### 20.9. Bottleneck IX model for idealized models (Figure S9C)

We used this model to mimic the real demography of non-African populations. We assumed the ancient population size of 30,000. The population experienced an instantaneous decrease to 3,000 at 35,000 generations ago, an instantaneous growth to 30,000 at 30,000 generations ago. Then, the population began exponential decline to 6,000 at 20,000 generations ago and instantaneously increased to 300,000 at 1,000 generations ago. 170 sequences of 800 Mb were simulated. The corresponding ms command for the simulation is

ms 170 800 -t 14400 -r 11520 1000000 -G 4694.427606513776 -eN 0.00083333 0.02 -eG 0.00083333 -101.6487102589958 -eN 0.016666666666666666 0.1 -eN 0.025 0.01 -eN 0.0291667 0.1

### 21. Models for estimating confidence interval

1000GP populations (Figure S22 – S26):

ACB:

ms 192 80000 -t 37.725120000000004 -r 30.180096000000002 10000 -G 307.97400639367373 -eN 0.0033742698361377437 0.35374201592996923 -eN 0.10237507717670723 0.012990813548107999 -eN 0.1132253281877044 0.9123469984986132

ASW:

ms 122 80000 -t 26.50224 -r 21.201792 10000 -G 56.10026737541272 -eN 0.013909291796913290 0.4582616412801333 -eN 0.1538498233982067 0.013891655950591346 -eN 0.16560167778269375 1.3718870555847353

BEB:

ms 172 80000 -t 62.65536 -r 50.124288 10000 -G 1215.8971532452542 -eN 0.0023403111314833293 0.058100695614868386 -eG 0.0023403111314833293 -30.907477873075123 -eN 0.03371108514231231 0.15320381209205405

CDX:

ms 186 80000 -t 25.248 -r 20.1984 10000 -G 495.40037108442965 -eN
0.004604646549934852 0.10216730038022814 -eG 0.004604646549934852 -
17.489419084798488 -eN 0.0835118490009135 0.4061216730038023
CEU:
ms 198 80000 -t 76.50768000000001 -r 61.206144 10000 -G 3486.9981711123683 -eN
8.189396733027973E-4 0.057518931432765964 -eN 0.01896877424771246
0.1280749854132291
CHB:
ms 206 80000 -t 148.45488 -r 118.763904 10000 -G 7477.99792685936 -eN
5.168378866508531E-4 0.020964753735276336 -eN 0.006336402867008054
0.06404208470614102
CHS:
ms 210 80000 -t 87.9024 -r 70.32192 10000 -G 4112.853998070424 -eN
8.149360963103421E-4 0.03502429967782449 -eN 0.011161187141954527
0.1114290394801507
CLM:
ms 188 80000 -t 26.05104 -r 20.840832 10000 -eN 0.006899708256449607
0.17470933981906286 -eN 0.04654813479653858 0.35472518563558303
ESN:
ms 198 80000 -t 32.99376 -r 26.395008 10000 -G 1399.7970544542743 -eN
6.732730258474137E-4 0.38967368375110933 -eN 0.11907424073256923
0.025241136505812008 -eN 0.14645778743979765 1.6907342479305179
FIN:
ms 198 80000 -t 21.88368 -r 17.506944 10000 -G 1545.9307685858762 -eN
0.0010069648928517855 0.21083108508258208 -eN 0.079627651288076
0.4667149218047421
GBR:
ms 182 80000 -t 68.96448 -r 55.171584 10000 -G 4780.2465803320365 -eN
4.8660205464359964E-4 0.09767810907876055 -eG 4.8660205464359964E-4
299.83121839376906 -eN 0.0026469357311387233 0.051108048665051926 -eG

0.0026469357311387233 -38.53061553796807 -eN 0.02777287031733699 0.1345666638826248

GIH:

ms 206 80000 -t 22.5384 -r 18.03072 10000 -G 319.3657263289911 -eN 0.005331789062104143 0.18217442231924183 -eG 0.005331789062104143 -7.108545561599834 -eN 0.1312801904940001 0.4459801938025769

GWD:

ms 226 80000 -t 71.53488 -r 57.227904 10000 -G 2480.4594823364796 -eN 6.955466408858352E-4 0.17812401446678877 -eN 0.055221144353749266 0.008796827505686736 -eN 0.06446476441787029 0.7480725486643718

IBS:

ms 214 80000 -t 101.40288 -r 81.122304 10000 -G 3879.3899843730937 -eN 7.379412692860351E-4 0.05711080395349718 -eN 0.001639355380662806 0.03441322376642557 -eG 0.001639355380662806 -55.51532072601329 -eN 0.019312961005874058 0.09179857613511569

ITU:

ms 204 80000 -t 45.81072 -r 36.648576 10000 -G 942.7096023053382 -eN 0.002494089782323459 0.09525456050461552 -eN 0.026664885830212617 0.2086358826056434

JPT:

ms 208 80000 -t 55.95024 -r 44.760192 10000 -G 3128.7376262633884 -eN 8.114087494514677E-4 0.07897017063733774 -eN 0.002182047525463559 0.04600945411494213 -eG 0.002182047525463559 -42.38616087573135 -eN 0.03372169017138904 0.17515849797963332

KHV:

ms 198 80000 -t 57.9072 -r 46.32576 10000 -G 1588.3658354039515 -eN 0.0019264561851523535 0.04689157824933687 -eG 0.0019264561851523535 -36.60290382061524 -eN 0.03792001517437307 0.17509118037135277

LWK:

ms 198 80000 -t 21.33888 -r 17.071104000000002 10000 -G 34.458048510849686 -eN 0.014319371341156495 0.6105362605722512 -eN 0.18803659662764166 0.02145942055065683 -eN 0.20879409785416014 2.0908763721432426

MSL:

ms 170 80000 -t 21.25872 -r 17.006976 10000 -G 127.20491855898698 -eN 0.003279354257615819 0.6589220799747115 -eN 0.18195660752091394 0.04585788796315112 -eN 0.23579811421655142 2.9051457472510105

MXL:

ms 128 80000 -t 72.2232 -r 57.77856 10000 -G 1185.9983336956855 -eN 0.0025552519587115612 0.04829030006978367 -eG 0.0025552519587115612 -35.997766981810344 -eN 0.030770096213141897 0.13333997939720202

PEL:

ms 170 80000 -t 104.52432 -r 83.619456 10000 -G 2449.33558491349 -eN 0.0014410414433505075 0.029316813541575778 -eN 0.01066574560965878 0.09490308092891683

PJL:

ms 192 80000 -t 40.10544 -r 32.084352 10000 -G 658.1508823918153 -eN 0.0035165490125809392 0.09882350125070315 -eG 0.0035165490125809392 -14.935034208218648 -eN 0.06458715193043525 0.2460234821011813

PUR:

ms 208 80000 -t 23.60832 -r 18.886656 10000 -eN 0.0038497802945414656 0.6750365972674041 -eN 0.014388104698224359 0.1546437865972674 -eG 0.014388104698224359 -15.582101828279491 -eN 0.07286054886643582 0.3846169486011711

STU:

ms 204 80000 -t 34.572 -r 27.657600000000002 10000 -G 510.43071934664556 -eN 0.004267252624293807 0.11325234293648039 -eN 0.02525451461678291 0.2616869142658799

TSI:

ms 214 80000 -t 75.82608 -r 60.660864000000004 10000 -G 2569.467980386571 -eN 0.0011371059177742527 0.05383899576504548 -eG 0.0011371059177742527 -19.704497360903815 -eN 0.04986270865479367 0.14062707712175018

YRI:

ms 216 80000 -t 64.91088 -r 51.928704 10000 -G 4776.570655987927 -eN 3.3888995289779744E-4 0.19814983250881826 -eN 0.0625962626099122 0.007882807936050167 -eN 0.06835272805758985 0.4330589879539455

HGDP-CEPH populations (Figure S27 – S32):

Adygei:

ms 32 80000 -t 19.19712 -r 15.357696 10000 -eN 0.005250411630207267 0.19837975696354454 -eG 0.005250411630207267 -8.596593090355556 -eN 0.11548669953457971 0.5117517627644147

Balochi:

ms 48 80000 -t 50.949600000000004 -r 40.75968 10000 -G 623.264941916954 -eN 0.004223024540924805 0.0719299072024118 -eG 0.004223024540924805 -32.99343443398211 -eN 0.032825823542035906 0.18482264826416694

Basque:

ms 46 80000 -t 26.153760000000002 -r 20.923008 10000 -G 751.4700408942055 -eN 0.0025016313707249523 0.15260520858186355 -eG 0.0025016313707249523 -8.224106050612061 -eN 0.1199670757097625 0.40097637968689775

Bedouin:

ms 92 80000 -t 36.622080000000004 -r 29.297664 10000 -G 244.38587570241546 -eN 0.008841200644961786 0.11524850581944007 -eG 0.008841200644961786 -16.31003863953942 -eN 0.05738997874929217 0.2544039005976722

Biaka:

ms 44 80000 -t 4.5912 -r 3.6729600000000002 10000 -eN 0.009592435429723692 3.6935703084161005 -eN 0.9319760745921495 0.09492943021432305 -eG 0.9319760745921495 -11.520947944349727 -eN 1.3098356260554107 7.379299529534762

Brahui:

ms 50 80000 -t 36.26112 -r 29.008896 10000 -G 372.4275509762127 -eN 0.006063547988128193 0.10453510536905644 -eG 0.006063547988128193 -20.411278092113488 -eN 0.05182198032003567 0.26600391824632

Burusho:

ms 48 80000 -t 11.5464 -r 9.23712 10000 -eN 0.015366836965198416 0.29627935980045733 -eG 0.015366836965198416 -8.16350928074288 -eN 0.1408168183093884 0.8250259821242986

Druze:

ms 84 80000 -t 16.18944 -r 12.951552 10000 -G 109.07256817628162 -eN 0.01288215226934864 0.24534511385199242 -eG 0.01288215226934864 -7.35809820577746 -eN 0.13234063031350007 0.5909037001897534

French:

ms 56 80000 -t 79.18416 -r 63.347328000000005 10000 -G 2198.9359820452387 -eN 0.001348902384433904 0.051501209332775646 -eG 0.001348902384433904 -24.36290322377743 -eN 0.03939915284814332 0.13014117975110173

Han:

ms 86 80000 -t 159.24816 -r 127.398528 10000 -G 5363.756025025552 -eN 7.370989032405503E-4 0.0191851510246649 -eN 0.005252449909354633 0.05743488653181299

Hazara:

ms 38 80000 -t 12.69792 -r 10.158336 10000 -eN 0.014217614003750299 0.25629394420503515 -eG 0.014217614003750299 -9.884907873225844 -eN 0.12245692270731047 0.7471459892643834

Japanese:

ms 54 80000 -t 33.88464 -r 27.107712 10000 -G 636.5261153273832 -eN 0.004000435354218995 0.07836471038204922 -eG 0.004000435354218995 -26.629980477176918 -eN 0.0528258276856906 0.2876064198999901

Kalash:

ms 44 80000 -t 3.04608 -r 2.436864 10000 -G -2.415743522809092 -eN 0.47028502523586235 3.1145603529782537

Makrani:

ms 50 80000 -t 24.11088 -r 19.288704 10000 -eN 0.00980388240380426 0.14526885787661006 -eG 0.00980388240380426 -17.111271203295853 -eN 0.06723955104118398 0.388146761959746

Mandenka:

ms 44 80000 -t 13.10688 -r 10.485504 10000 -eN 0.3296277261231593 0.048304401962938545 -eG 0.3296277261231593 -17.85185708722175 -eN 0.5825077171252933 4.411191679484362

Maya:

ms 42 80000 -t 16.673280000000002 -r 13.338624000000001 10000 -eN 0.008589230112361738 0.09687356057116536 -eG 0.008589230112361738 -25.14802627151756 -eN 0.08111110798113147 0.600155458314141

Mozabite:

ms 54 80000 -t 17.42304 -r 13.938432 10000 -G 34.034495761040134 -eN 0.03302787085100088 0.32494903300457323 -eG 0.03302787085100088 -4.504618045303915 -eN 0.1472698871889101 0.543638767976197

Palestinian:

ms 92 80000 -t 22.33968 -r 17.871744 10000 -eN 0.008844463035145432 0.1913151844610129 -eG 0.008844463035145432 -7.866119095624621 -eN 0.11128708520369861 0.4282675490427795

Pathan:

ms 48 80000 -t 42.96432 -r 34.371456 10000 -G 497.59849646988573 -eN 0.005312881946315924 0.07109899563172419 -eG 0.005312881946315924 -47.5703288285576 -eN 0.02861662601453915 0.21543085052899708

Russian:

ms 50 80000 -t 22.14048 -r 17.712384 10000 -G 258.609263568537 -eN 0.006473721471777433 0.1874647704114816 -eN 0.047392730886604 0.4300394571391406

Sardinian:

ms 56 80000 -t 19.90512 -r 15.924096 10000 -G 209.67966756103402 -eN 0.008361464215634221 0.17321372591574427 -eG 0.008361464215634221 -11.622404927783043 -eN 0.09674763618341507 0.48385541006535

Sindhi:

ms 48 80000 -t 28.07088 -r 22.456704000000002 10000 -eN 0.006697787857121225 0.12812708401019132 -eG 0.006697787857121225 -17.75301871013936 -eN 0.06131324436460199 0.33785331988167094

Yakut:

ms 50 80000 -t 10.99008 -r 8.792064 10000 -G 107.01050381191519 -eN 0.010397096462590117 0.32870370370370366 -eN 0.09403222060369128 0.8740391334730957

Yoruba:

ms 44 80000 -t 30.6816 -r 24.54528 10000 -G 298.1592873494309 -eN 0.0028204398926000218 0.4313047559449311 -eN 0.1396195363438095 0.026126408010012515 -eN 0.16980043486634022 2.1113579474342927

**Supplementary figures**

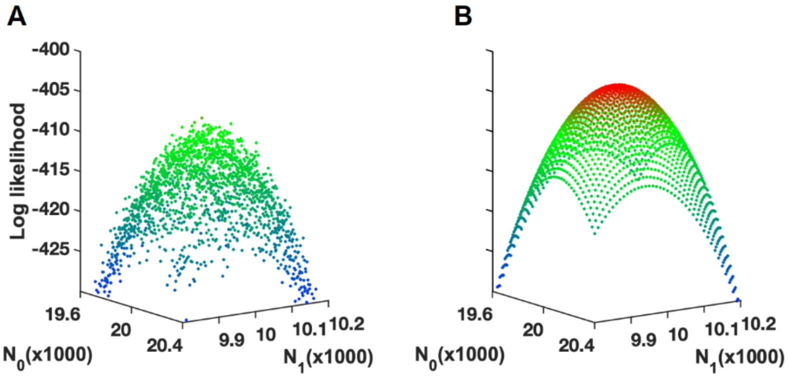

**Figure S1. Comparison of likelihood surfaces based on simulations and FitCoal.** (**A**) Likelihood surface of a SFS based on simulation approach. (**B**) Likelihood surface of the SFS based on FitCoal. The sample size is 100. The SFS is obtained under a demography that the population size increases from 10,000 to 20,000 at standard coalescent time 0.2. The surfaces were obtained conditional on standard coalescent time 0.2 while the current ($N_0$) and the ancestral population size ($N_1$) varied in the instantaneous growth model. Red dots indicate large likelihoods, and blue dots indicate small likelihoods.

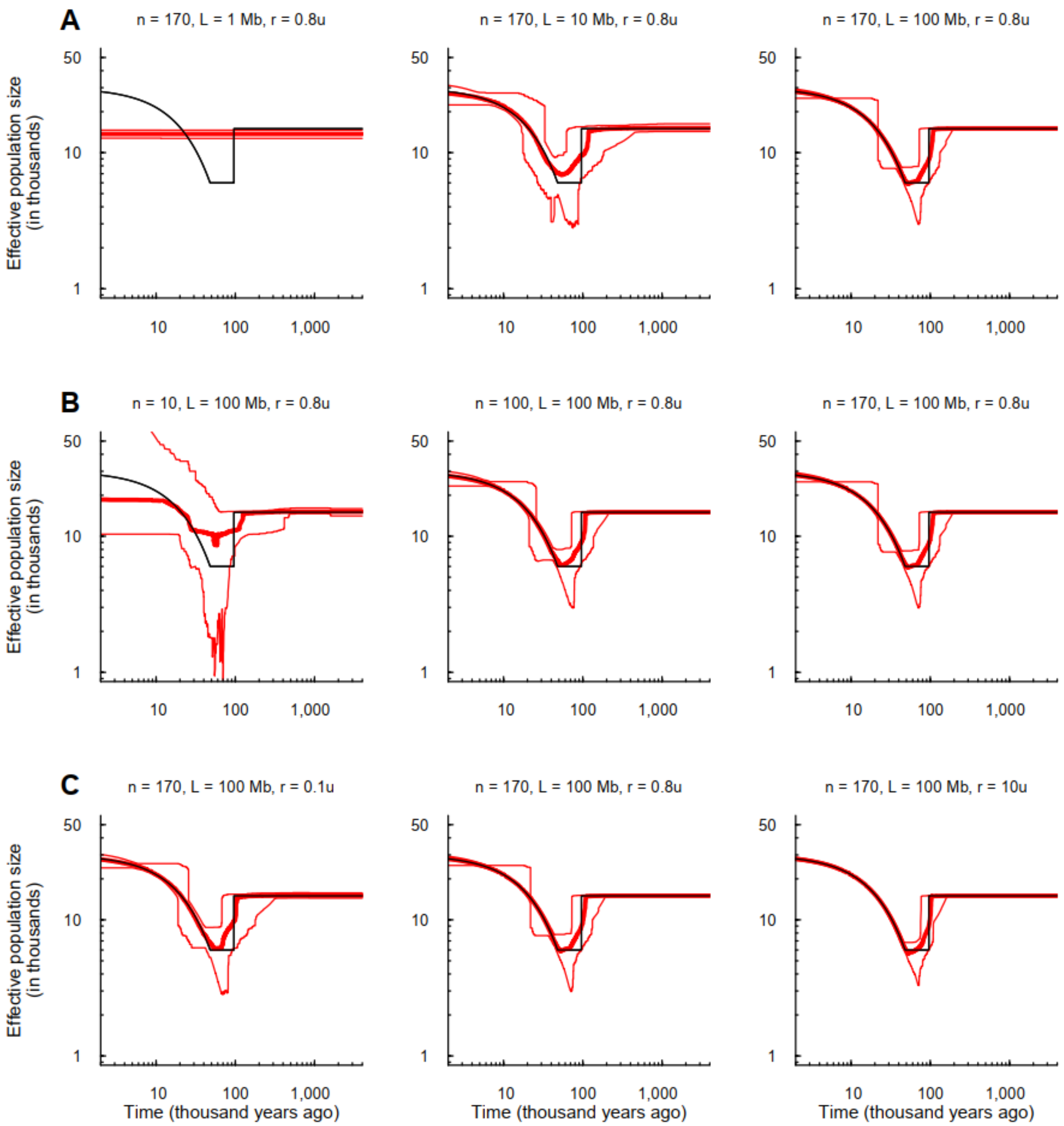

**Figure S2. Effects of sequence length (A), sample size (B), and recombination rate (C) in the FitCoal inference.** Thin black lines indicate true models. Thick red lines are the medians of the estimated histories of FitCoal; thin red lines are 2.5 and 97.5 percentiles of the estimated histories of FitCoal. *n* is the number of simulated sequences, *L* the length of the simulated sequences, and *r* the recombination rate relative to the mutation rate ($\mu$). Other parameters are the same with Figure 2. The corresponding commands for simulations are described above.

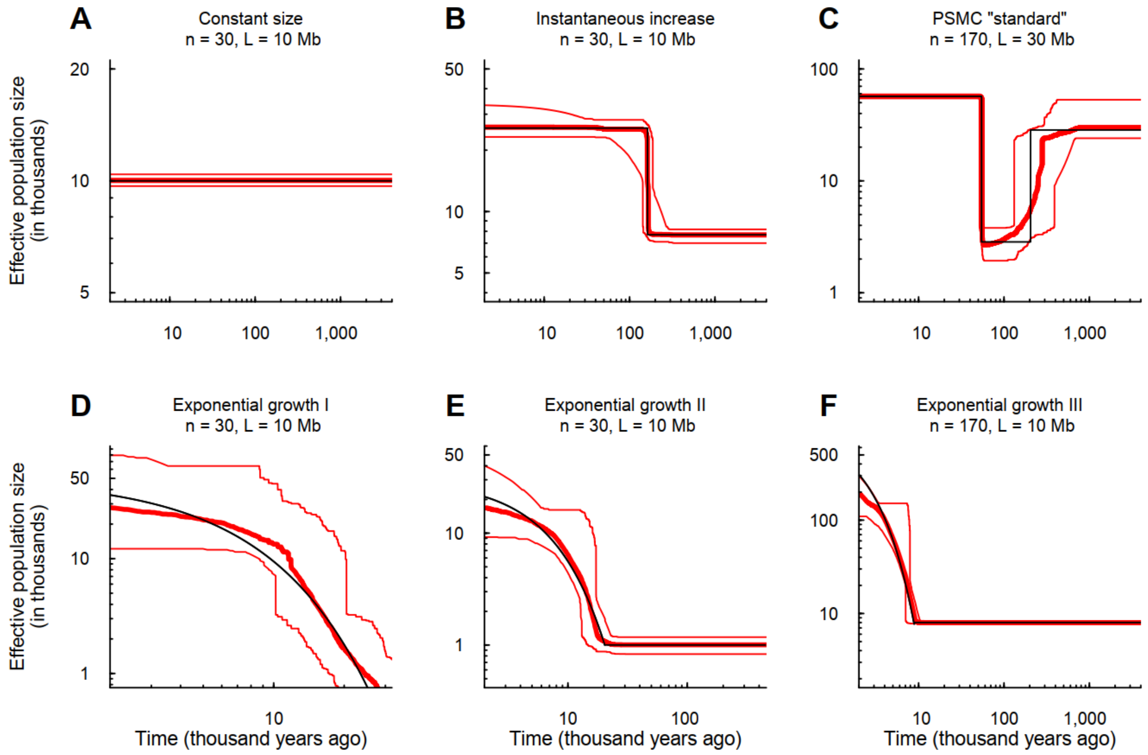

**Figure S3. Verification of FitCoal accuracy with truncated SFS.** (**A**) Constant size model. (**B**) Instantaneous increase model. (**C**) PSMC "standard" model. (**D**) Exponential growth I model. (**E**) Exponential growth II model. (**F**) Exponential growth III model. The six models are the same as Figure 2. Thin solid black lines indicate true models. Thick red lines are the medians of the estimated histories of FitCoal; thin red lines are 2.5 and 97.5 percentiles of the estimated histories of FitCoal. *n* is the number of simulated sequences and *L* the length of the simulated sequences. 10% SFS types (high frequency mutations) were discarded. The corresponding commands for simulations are described above.

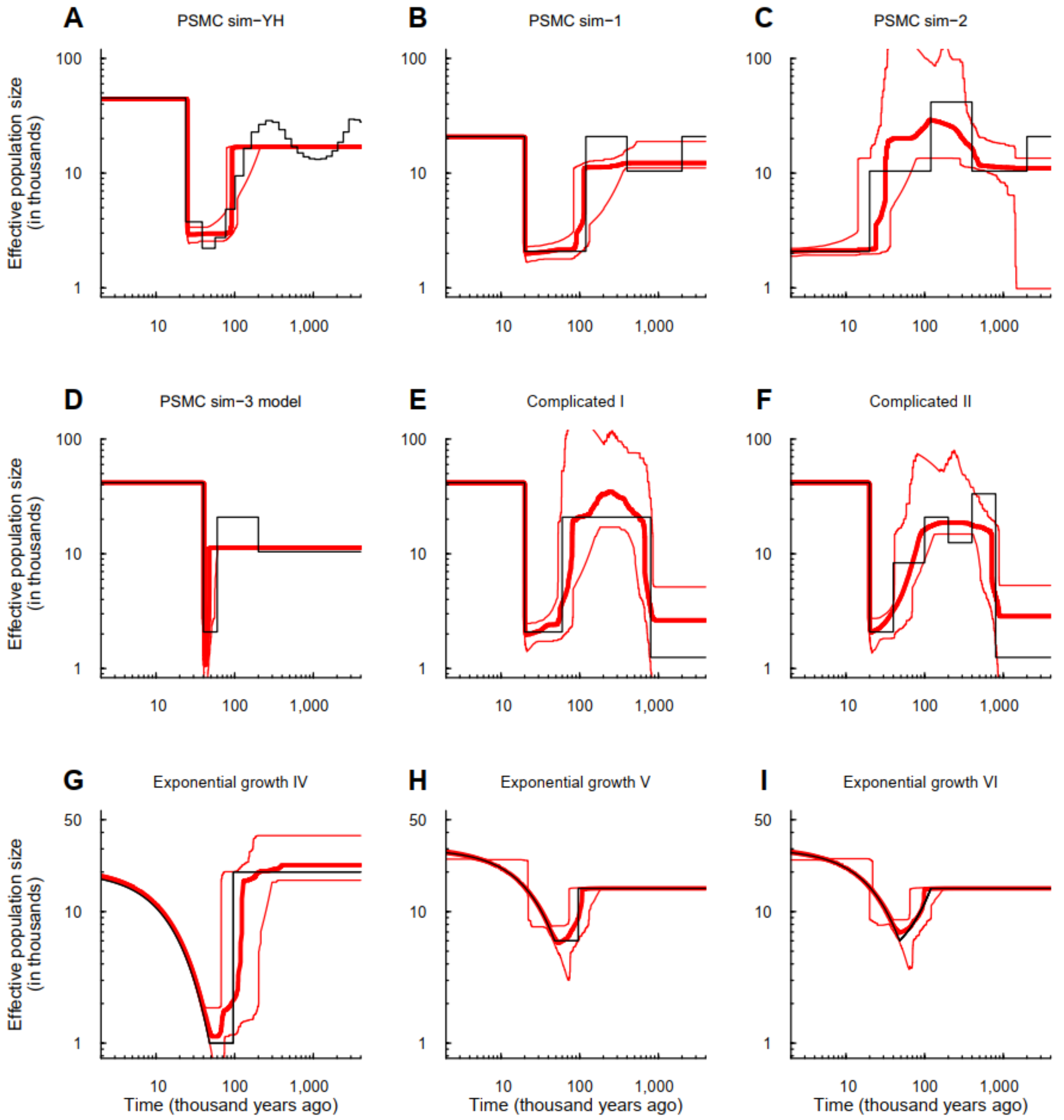

**Figure S4. Verification of FitCoal accuracy with truncated SFS under more complexed models.** (**A**) PSMC sim-YH model. (**B**) PSMC sim-1 model. (**C**) PSMC sim-2 model. (**D**) PSMC sim-3 model. (**E**) Complicated I model. (**F**) Complicated II model. (**G**) Exponential growth IV model. (**H**) Exponential growth V model. (**I**) Exponential growth VI model. Thin solid black lines indicate true models. Thick red lines are the medians of the estimated histories of FitCoal; thin red lines are 2.5 and 97.5 percentiles of the estimated histories of FitCoal. The number of simulated sequences is 170 and the length of the simulated sequences is100Mb. 10% SFS types (high frequency mutations) were discarded. The corresponding commands for simulations are described above.

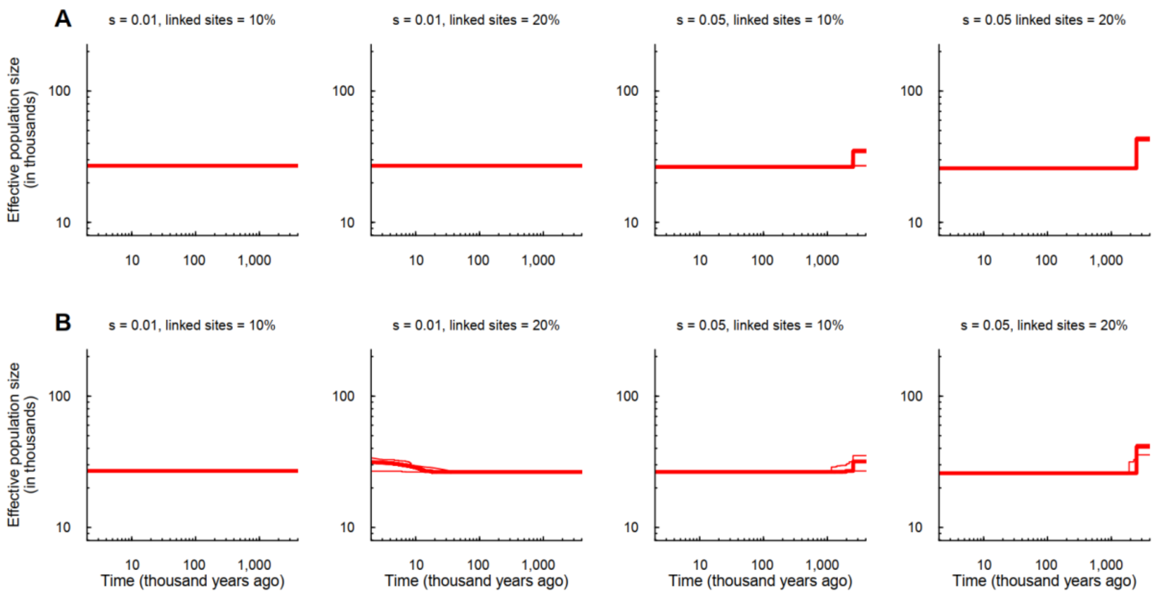

**Figure S5. Effects of positive selection on demographic inference.** (**A**) Demographic histories inferred by using the full size SFSs. (**B**) Demographic histories inferred by using the truncated SFSs. The constant size model was considered with different selection strength (s) and the percentage of loci affected by positive selection. $n = 202$. Thick red lines are the medians of the estimated histories of FitCoal; thin red lines are 2.5 and 97.5 percentiles of the estimated histories of FitCoal.

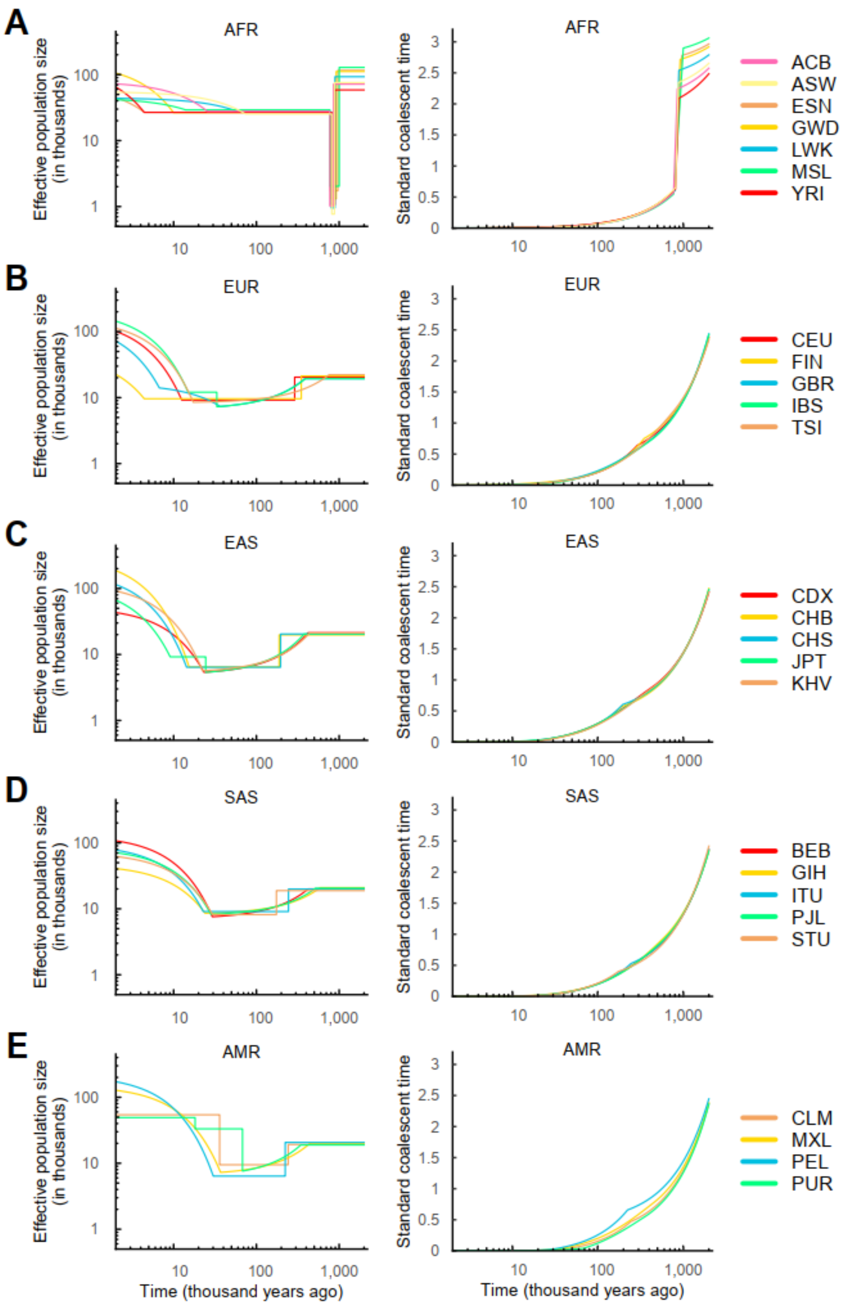

**Figure S6. Inferred demographic histories and standard coalescent times of 1000GP populations.** (**A**) African populations. (**B**) European populations. (**C**) East Asian populations. (**D**) South Asian populations. (**E**) American populations. The left column is the inferred demographic histories, and the right column is calendar time *vs* standard coalescent time. The results are the same with Figure 3.

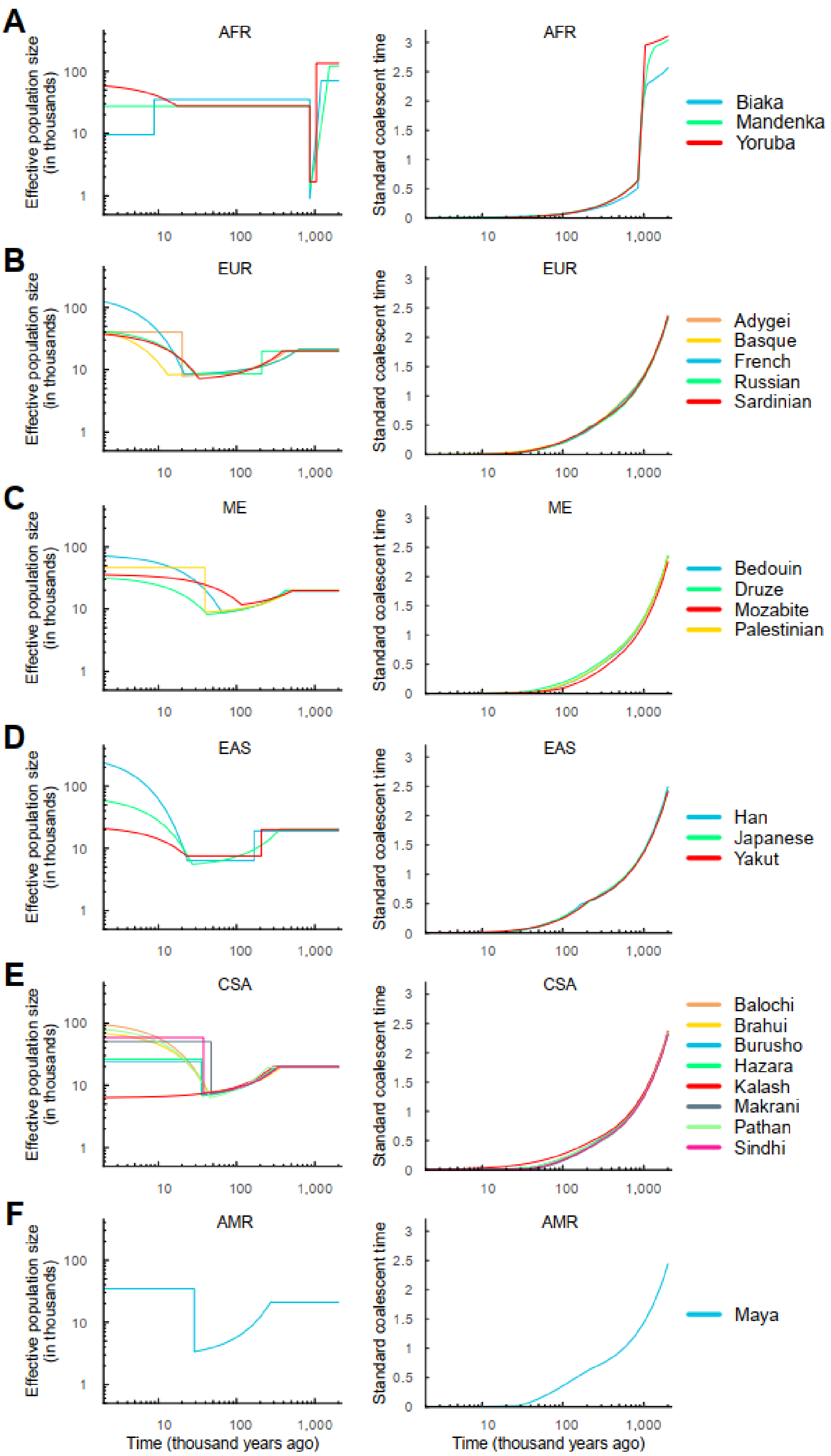

**Figure S7. Inferred demographic histories and standard coalescent times of HGPD-CEPH populations.** (**A**) African populations. (**B**) European populations. (**C**) Middle East populations. (**D**) East Asian populations. (**E**) Central & South Asian populations. (**F**) American population. The left column is the inferred demographic histories, and the right column is calendar time *vs* standard coalescent time. The results are the same with Figure 3.

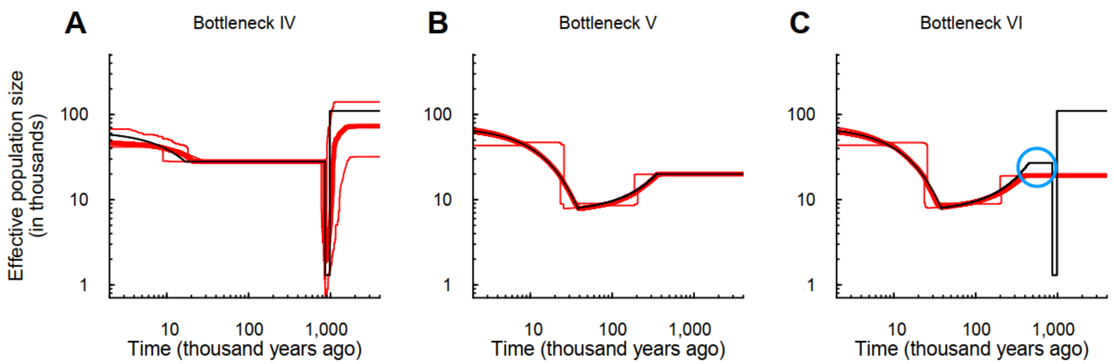

**Figure S8. Verification of the HGDP-CEPH inferred super bottleneck.** (**A**) Bottleneck IV model and the estimated histories. The bottleneck IV mimics the estimated demography of HGDP-CEPH African population. (**B**) Bottleneck V model and the estimated histories. The bottleneck V mimics the estimated demography of HGDP-CEPH non-African population. (**C**) Bottleneck VI model and the estimated histories. The bottleneck VI mimics the true demography of HGDP-CEPH non-African population. Thin black lines indicate three models. Thick red lines are the medians of the estimated histories of FitCoal; thin red lines are 2.5 and 97.5 percentiles of the estimated histories of FitCoal. Blue circles indicate the population size gap. 10% SFS types (high frequency mutations) were discarded. The number of simulated sequences is 44 in Bottleneck IV, and 56 in Bottleneck V and VI, as the average sampled sequences in the HGDP-CEPH African and non-African populations. The length of simulated sequences is 800 Mb.

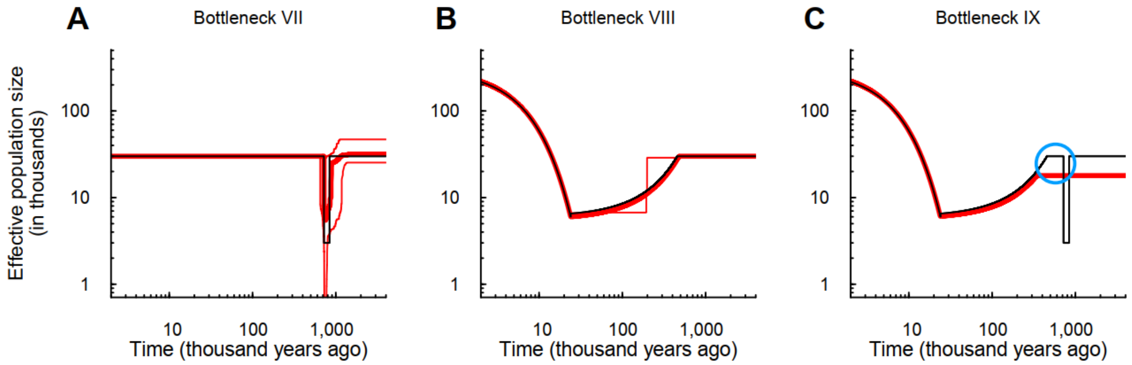

**Figure S9. Verification of the super bottleneck in artificial models.** (**A**) Bottleneck VII model and the estimated histories. The artificial bottleneck VII represents the authors-altered demography of African population. (**B**) Bottleneck VIII model and the estimated histories. The artificial bottleneck VIII represents the authors-altered estimated demography of non-African population. (**C**) Bottleneck IX model and the estimated histories. The bottleneck IX represents the authors-altered true demography of non-African population. Thin black lines indicate models. Thick red lines are the medians of the estimated histories of FitCoal; thin red lines are 2.5 and 97.5 percentiles of the estimated histories of FitCoal. Blue circle indicates the population size gap. 10% SFS types (high frequency mutations) were discarded. The number of simulated sequences is 170, and the length of simulated sequences is 800 Mb.

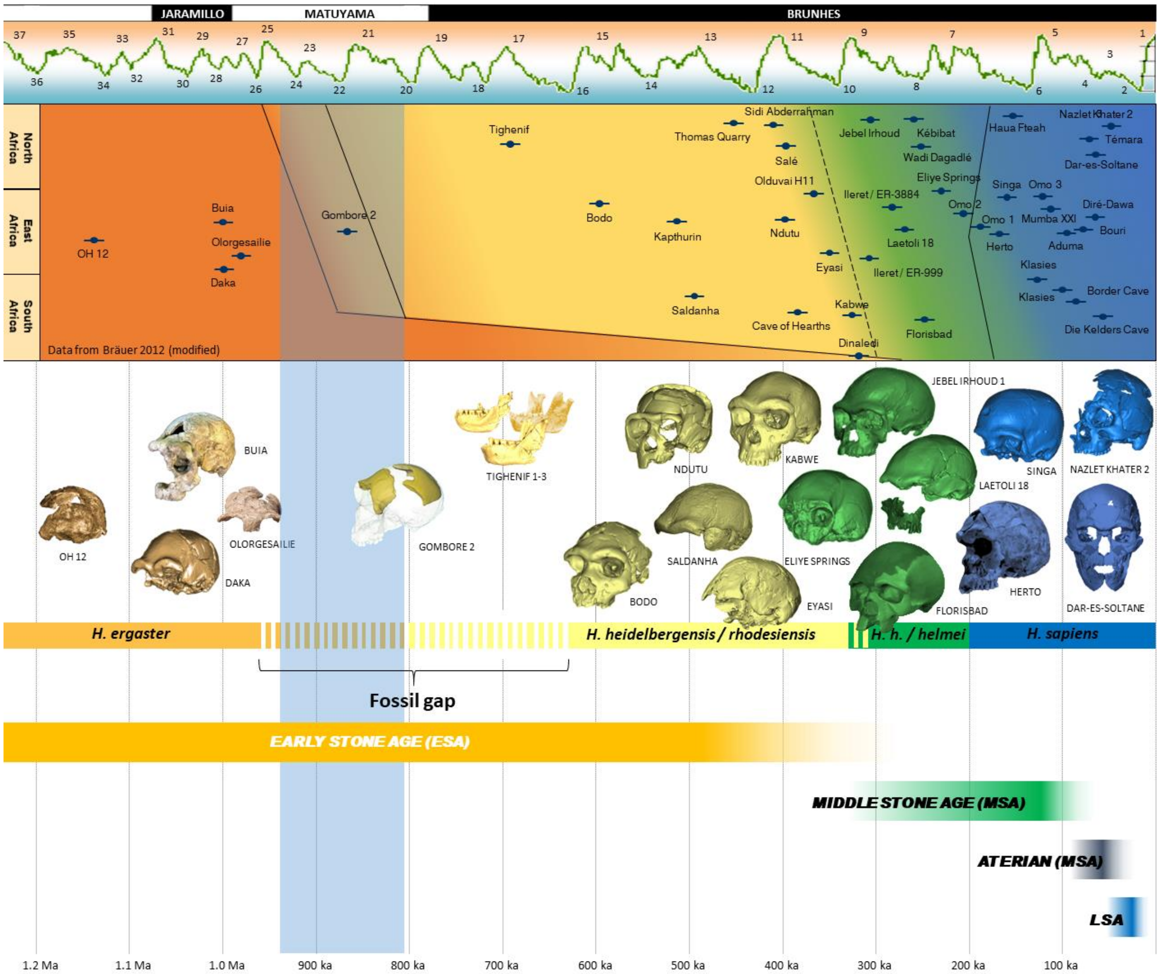

**Figure S10. Distribution of the human fossil record in Africa from the late Early Pleistocene to present.** It is compared to paleomagnetic chrons and a schematic climatic curve (above), as well as to the major stages of the African Palaeolithic (below). A gap in the human fossil record between 900 and 600 kyr ago is apparent. It is coincident, in turn, with the super bottleneck of 930-813 kyr ago (highlighted in pale blue). Data modified from Rightmire (1996, 2008); Mounier et al. (2011); Bräuer (2012); Manzi (2016) (ref. (Bräuer, 2012; Manzi, 2016; Mounier et al., 2011; Rightmire, 1996, 2008).

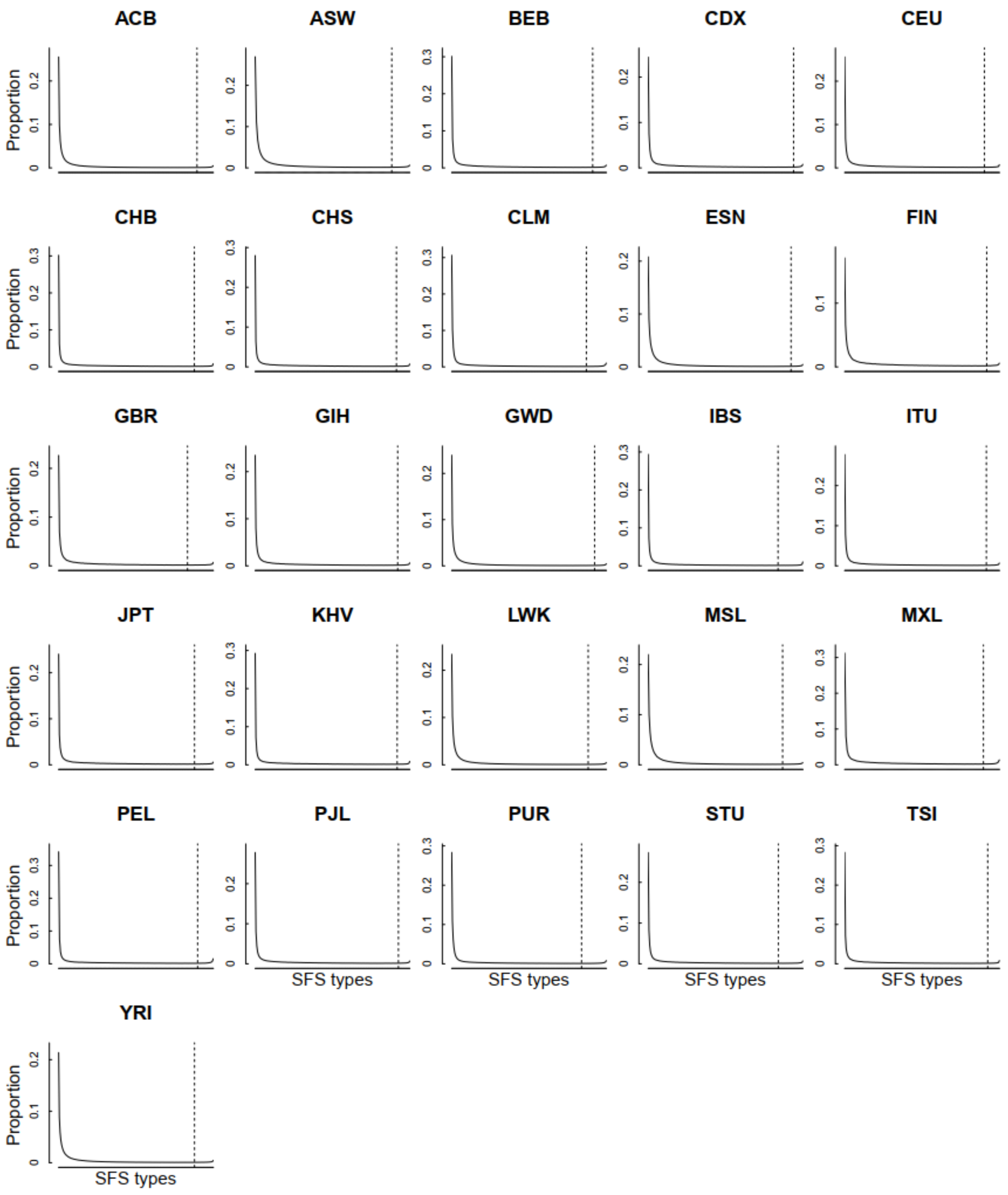

**Figure S11. The observed SFSs of 1000GP populations.** Solid lines indicate the observed SFS, and the $x$-axis is the SFS types, ranging from 1 to $(n-1)$. Dash lines indicate the threshold of truncating SFS.

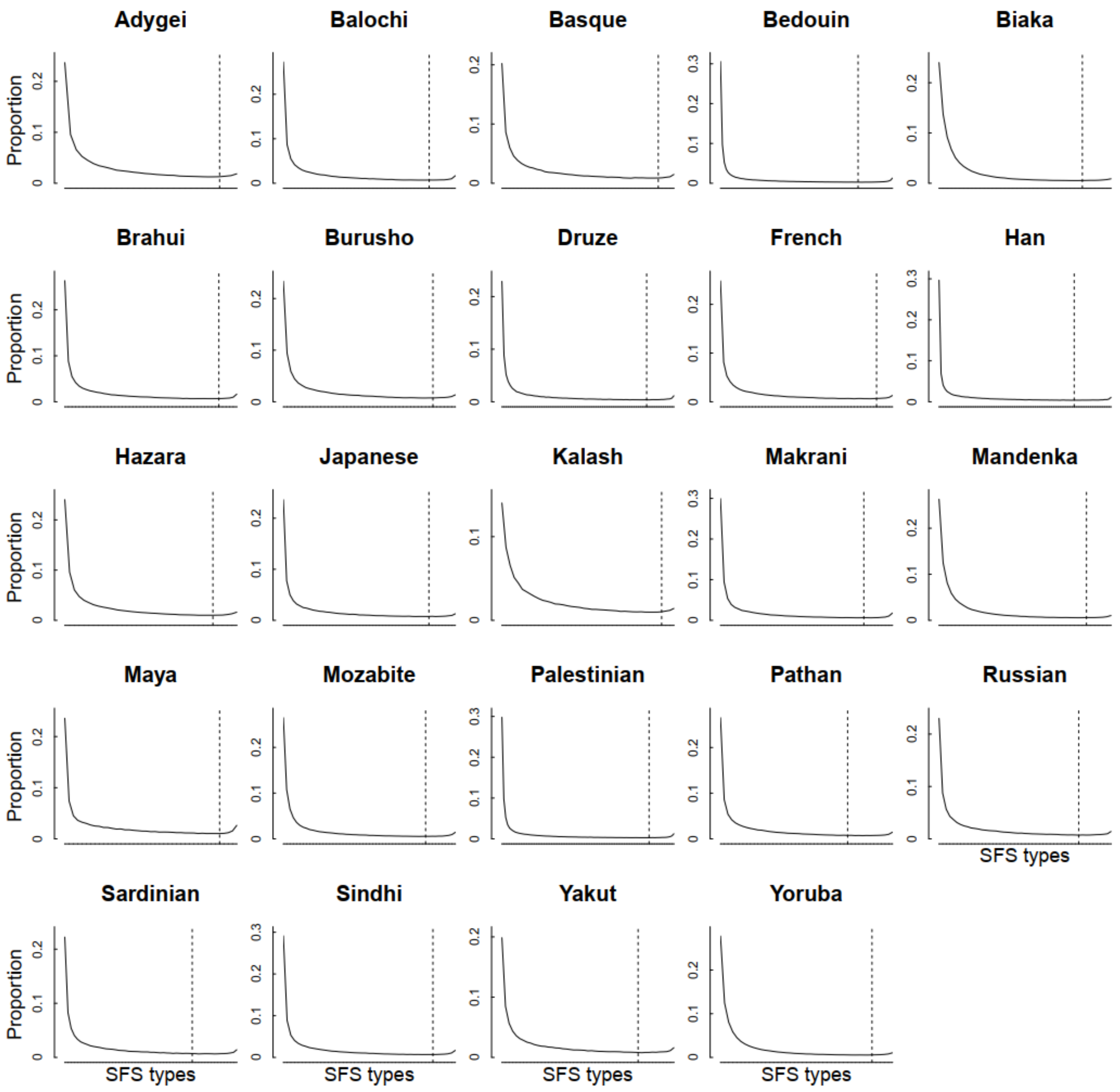

**Figure S12. The observed SFSs without missing data of HGDP-CEPH populations.** Solid lines indicate the observed SFS, and the $x$-axis is the SFS types, ranging from 1 to $(n-1)$. Dash lines indicate the threshold of truncating SFS.

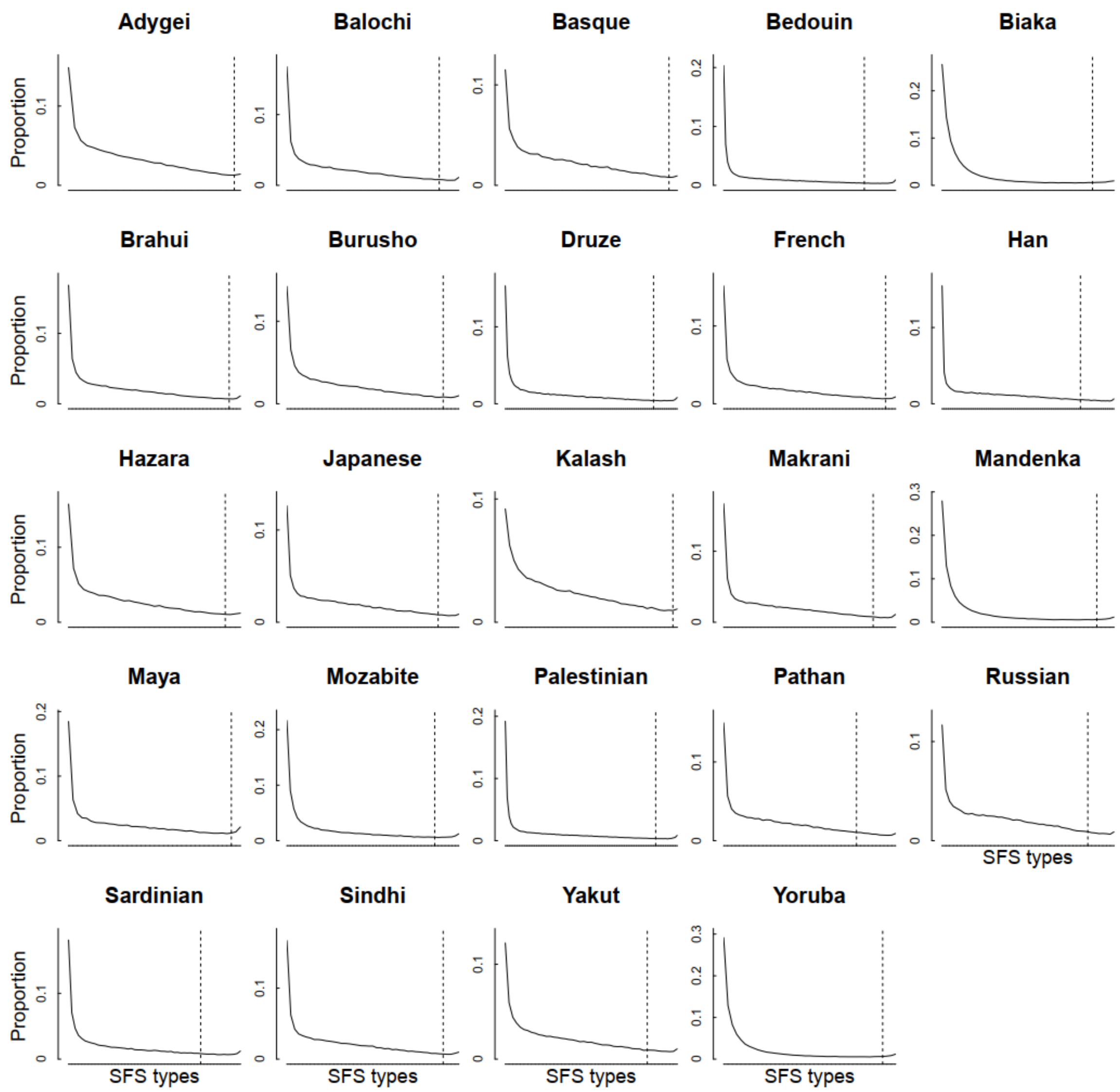

**Figure S13. The observed SFSs with one missing individual of HGDP-CEPH populations.** Solid lines indicate the observed SFS, and the *x*-axis is the SFS types, ranging from 1 to $(n-3)$. Dash lines indicate the threshold of truncating SFS.

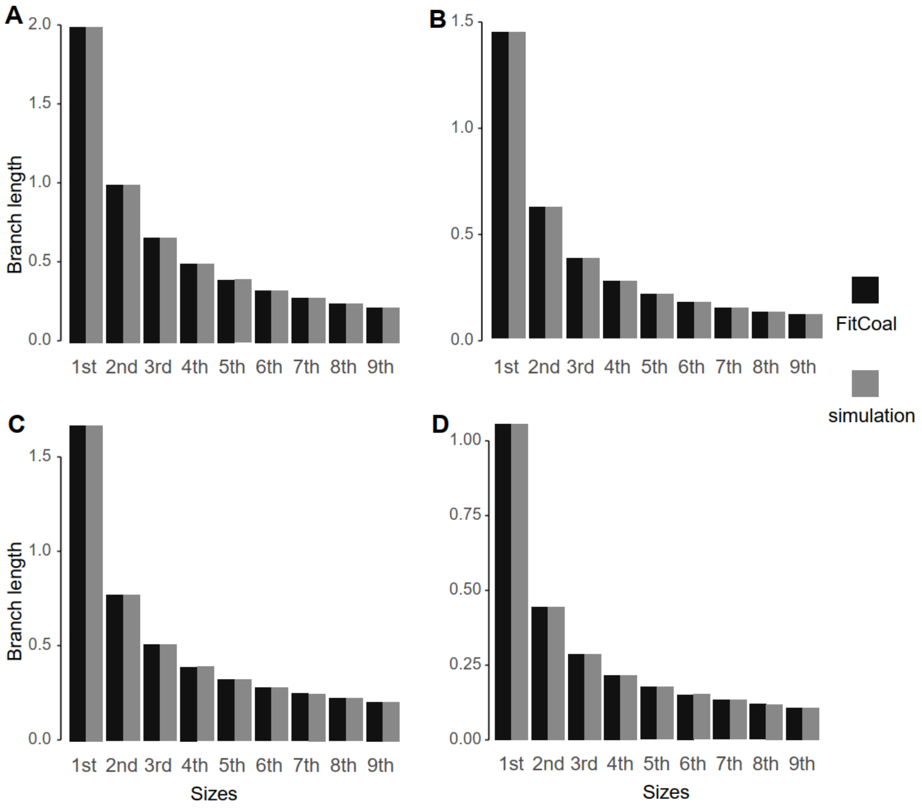

**Figure S14. Comparison of the expected branch lengths of FitCoal and the average branch lengths of coalescent simulations.** (**A**) Constant size model. (**B**) Exponential growth model. (**C**) Bottleneck model. (**D**) Complex model. The models, the related parameters, and the ms command lines are described above. $n = 10$. To calculate the average, the number of iterations is $10^6$ for coalescent simulations.

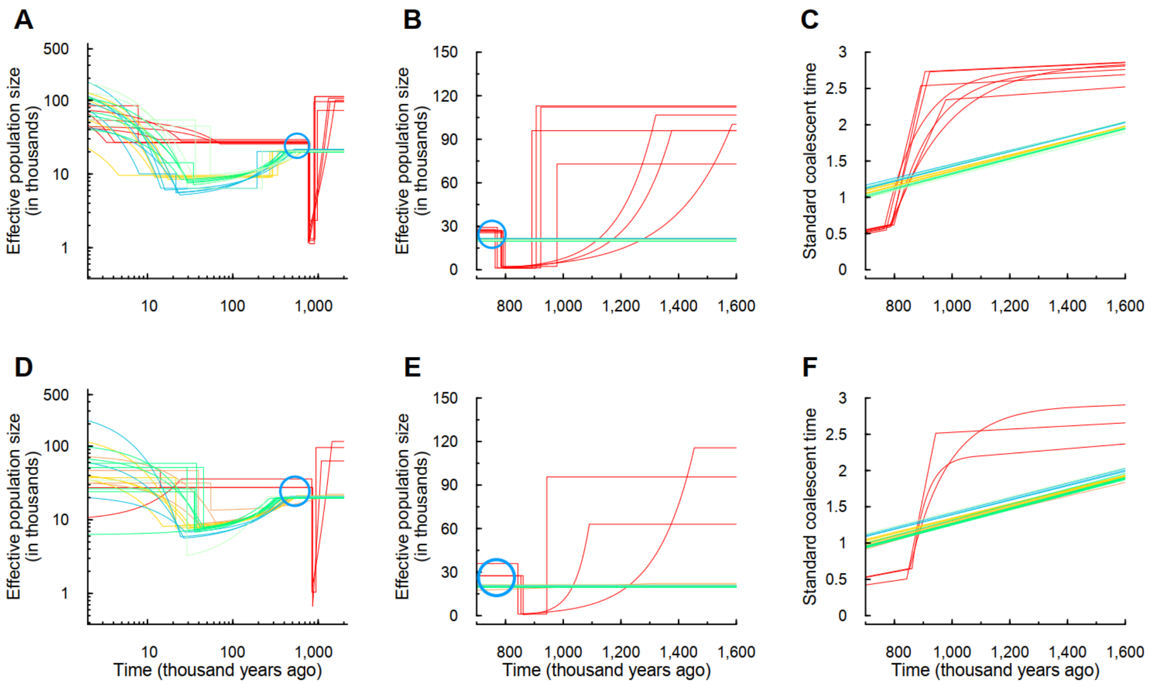

**Figure S15. Inferred demographic histories of 1000GP and HGPD-CEPH populations using the same truncating SFS standard for each data set.** (**A**) Estimated histories of 26 1000GP populations. (**B**) Linear-scaled estimated histories of 1000GP populations during the super bottleneck. (**C**) Calendar time *vs* standard coalescent time conditional on the estimated histories of 1000GP populations. (**D**) Estimated histories of 24 HGPD-CEPH populations. (**E**) Linear-scaled estimated histories of HGPD-CEPH populations during the super bottleneck. (**F**) Calendar time *vs* standard coalescent time conditional on the estimated histories of HGPD-CEPH populations. Red lines are the estimated histories of African populations; yellow lines stand for the European populations; brown lines the Middle East populations; blue lines the East Asian populations; green lines the Central or South Asian populations; dark sea green lines the American populations. Blue circle indicates the population size gap. We truncated 10% SFS types for 1000GP populations, and 15% for HGDP-CEPH populations. We assumed a mutation rate of $1.2 \times 10^{-8}$ per base per generation and a generation time of 24 years, the same as Figure 2.

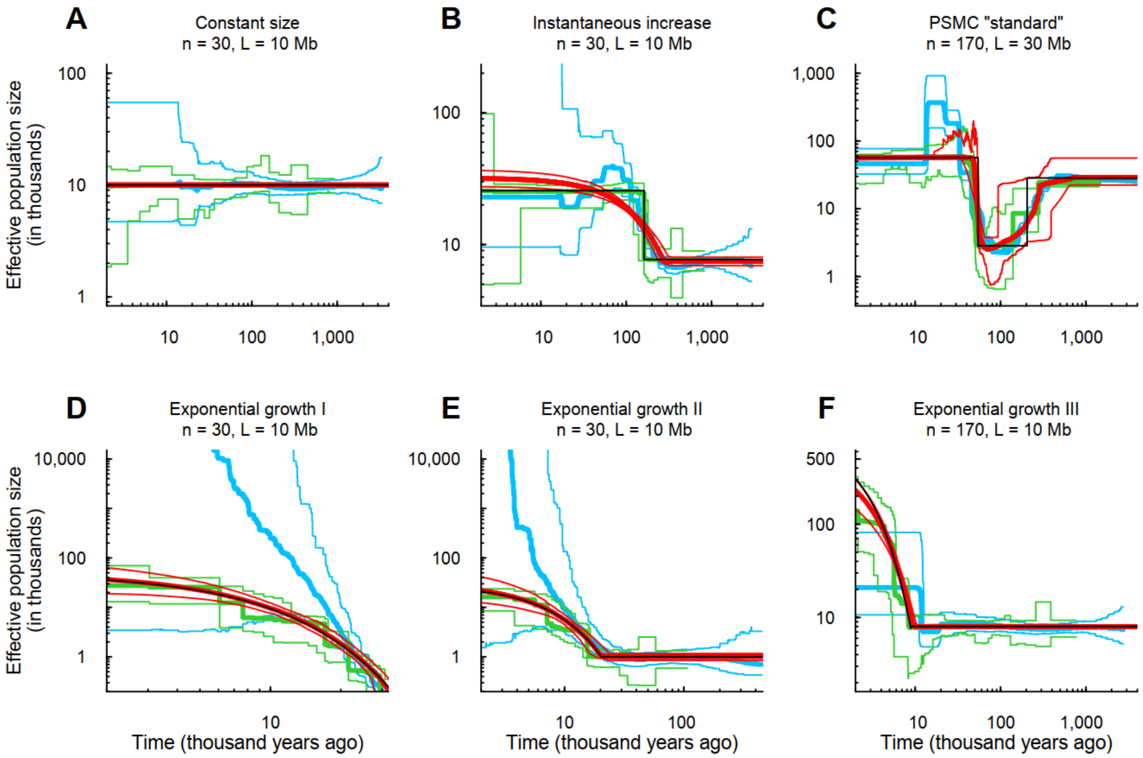

**Figure S16. Estimated demographic histories of FitCoal conditional on exponential change and instantaneous decline, stairway plot, and PSMC using simulated samples.** (**A**) Constant size model. (**B**) Instantaneous increase model. (**C**) PSMC "standard" model. (**D**) Exponential growth I model. (**E**) Exponential growth II model. (**F**) Exponential growth III model. Thin black lines indicate true models. Thick red lines are the medians of FitCoal histories estimated conditional on exponential change. Green and blue lines indicate the results of stairway plot and PSMC, respectively, which are obtained from the previous study (Liu and Fu, 2015). Other parameters are the same with Figure 2.

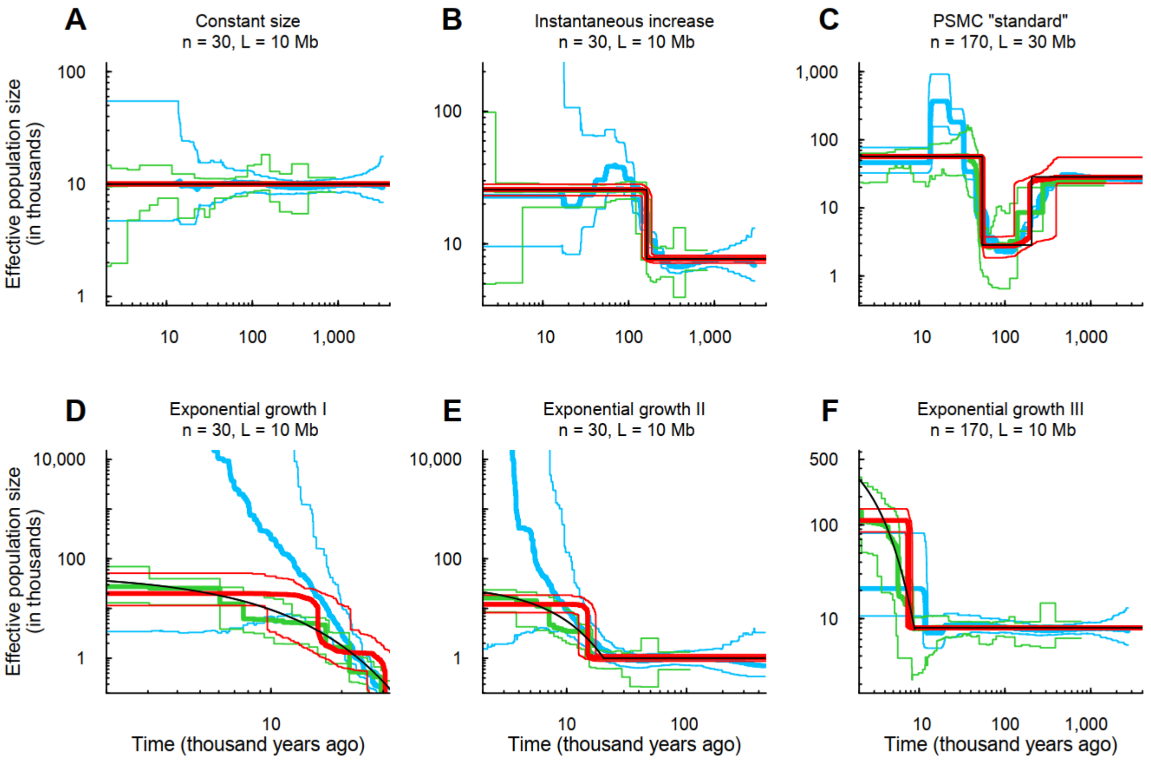

**Figure S17. Estimated demographic histories of FitCoal conditional on instantaneous change, stairway plot, and PSMC using simulated samples.** (**A**) Constant size model. (**B**) Instantaneous increase model. (**C**) PSMC "standard" model. (**D**) Exponential growth I model. (**E**) Exponential growth II model. (**F**) Exponential growth III model. FitCoal inference was performed conditional on instantaneous change. Other parameters are the same with Figure 2.

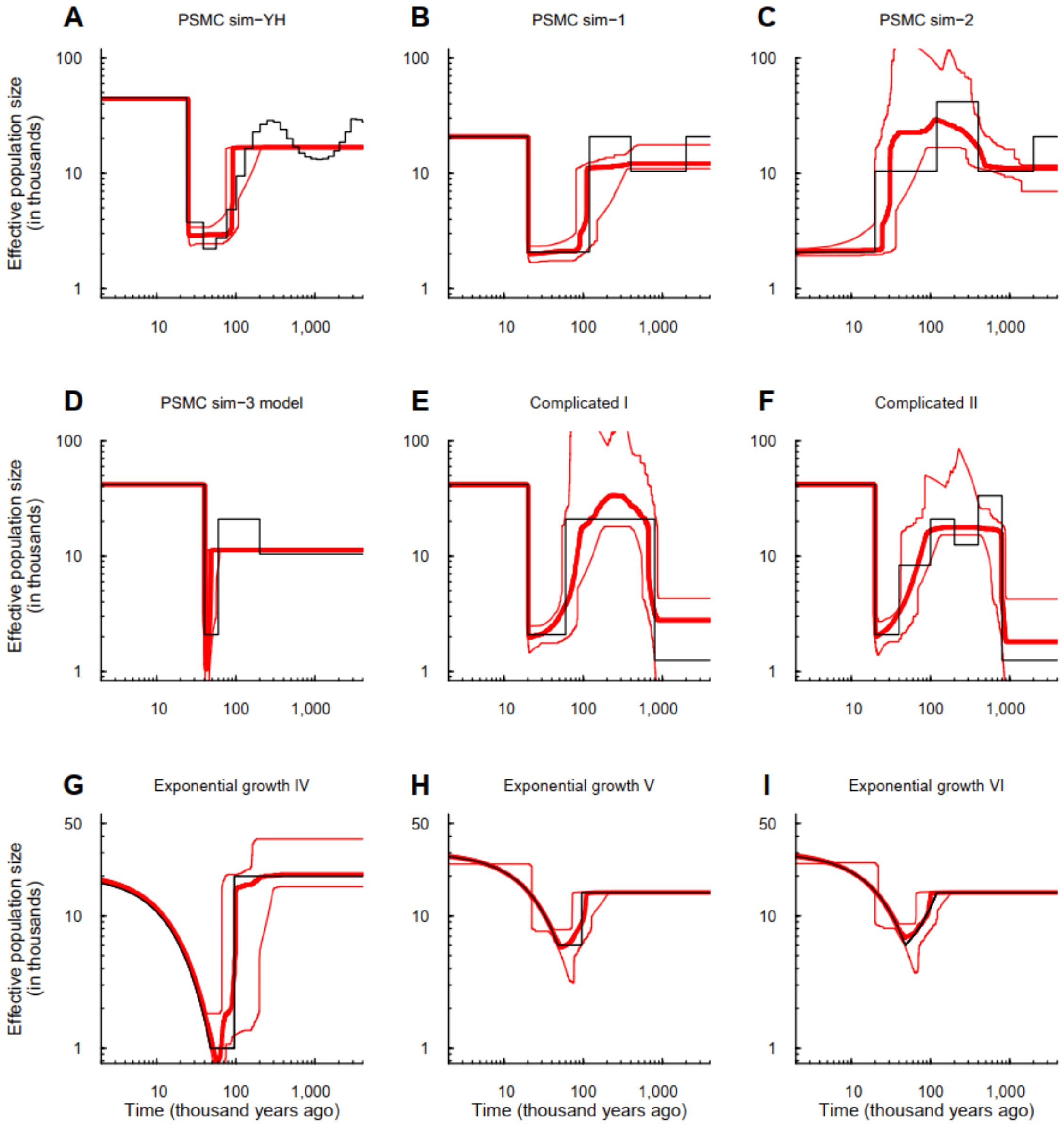

**Figure S18. Verification of the accuracy of FitCoal using simulated samples under complex models.** (**A**) PSMC sim-YH model. (**B**) PSMC sim-1 model. (**C**) PSMC sim-2 model. (**D**) PSMC sim-3 model. (**E**) Complicated I model. (**F**) Complicated II model. (**G**) Exponential growth IV model. (**H**) Exponential growth V model. (**I**) Exponential growth VI model. Thin black lines indicate true models. Thick red lines are the medians of the estimated histories of FitCoal; thin red lines are 2.5 and 97.5 percentiles of the estimated histories of FitCoal. The number of simulated sequences is 170 and the length of the simulated sequences is 100Mb in all models. Other parameters are the same with Figure 2. The corresponding commands for simulations are described above.

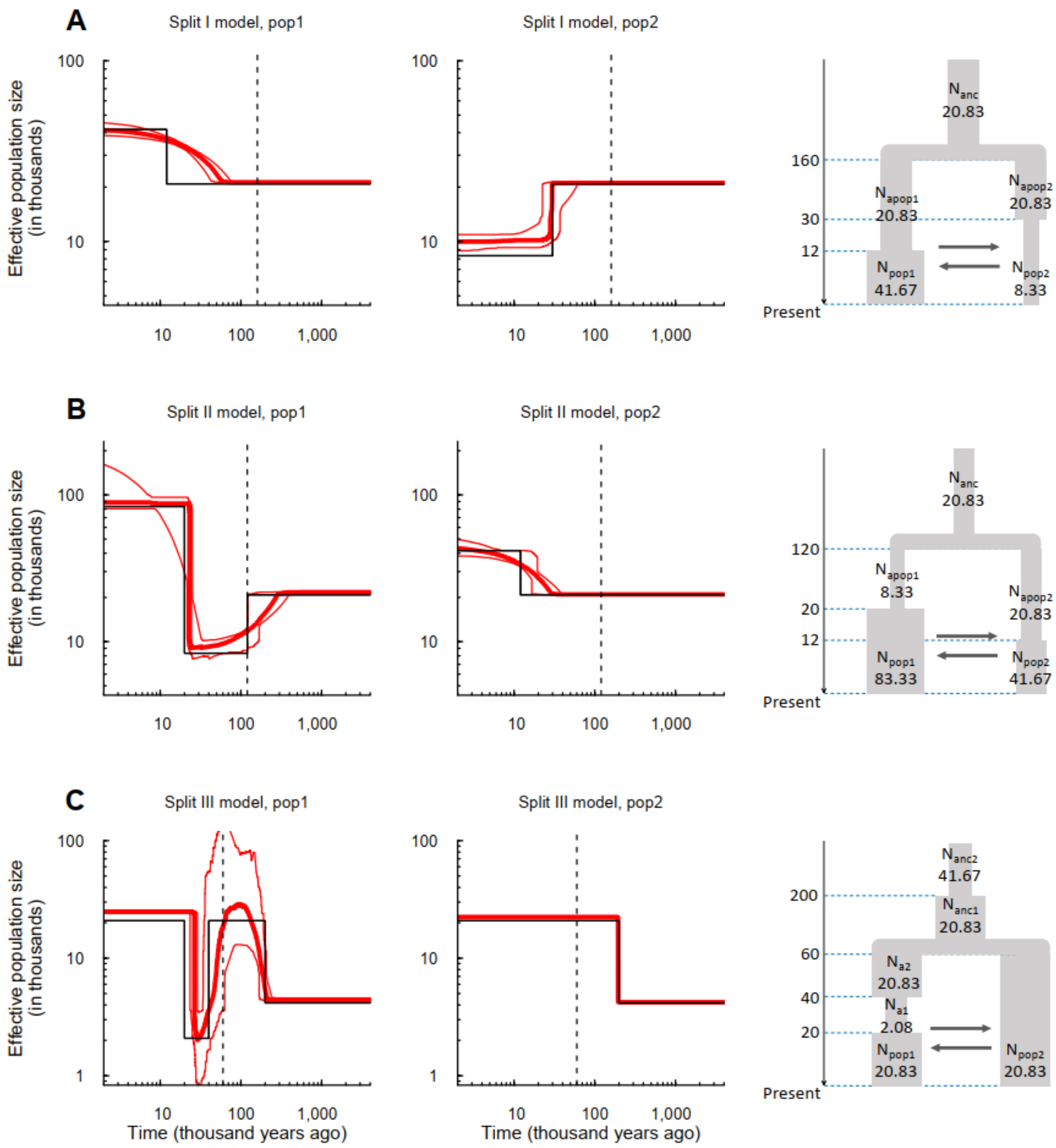

**Figure S19. Verification of FitCoal accuracy under three migration models.** (**A**) Inferred histories of two populations under Split I model. The model assumes that the two populations split at 160 kyr ago, and the first population had an instantaneous growth, and the second one had a population size decline. Migration occurred populations. The population size during each stage is shown in the model (the right panel). (**B**) Inferred histories of two populations under Split II model. (**C**) Inferred histories of two populations under Split III model. Thin solid black lines indicate true models. Thin dash lines indicate split times. Thick red lines are the medians of the estimated histories of FitCoal; thin red lines are 2.5 and 97.5 percentiles of the estimated histories of FitCoal. In three rightmost figures, the unit of time is 1,000 years and the unit of population size is 1,000. The number of simulated sequences is 170 and the migration rate ($4Nm$) is 4. The length of the simulated sequences is 30Mb in the first two models and 100MB in the last model. Other parameters are the same with Figure 2. The corresponding commands for simulations are described above.

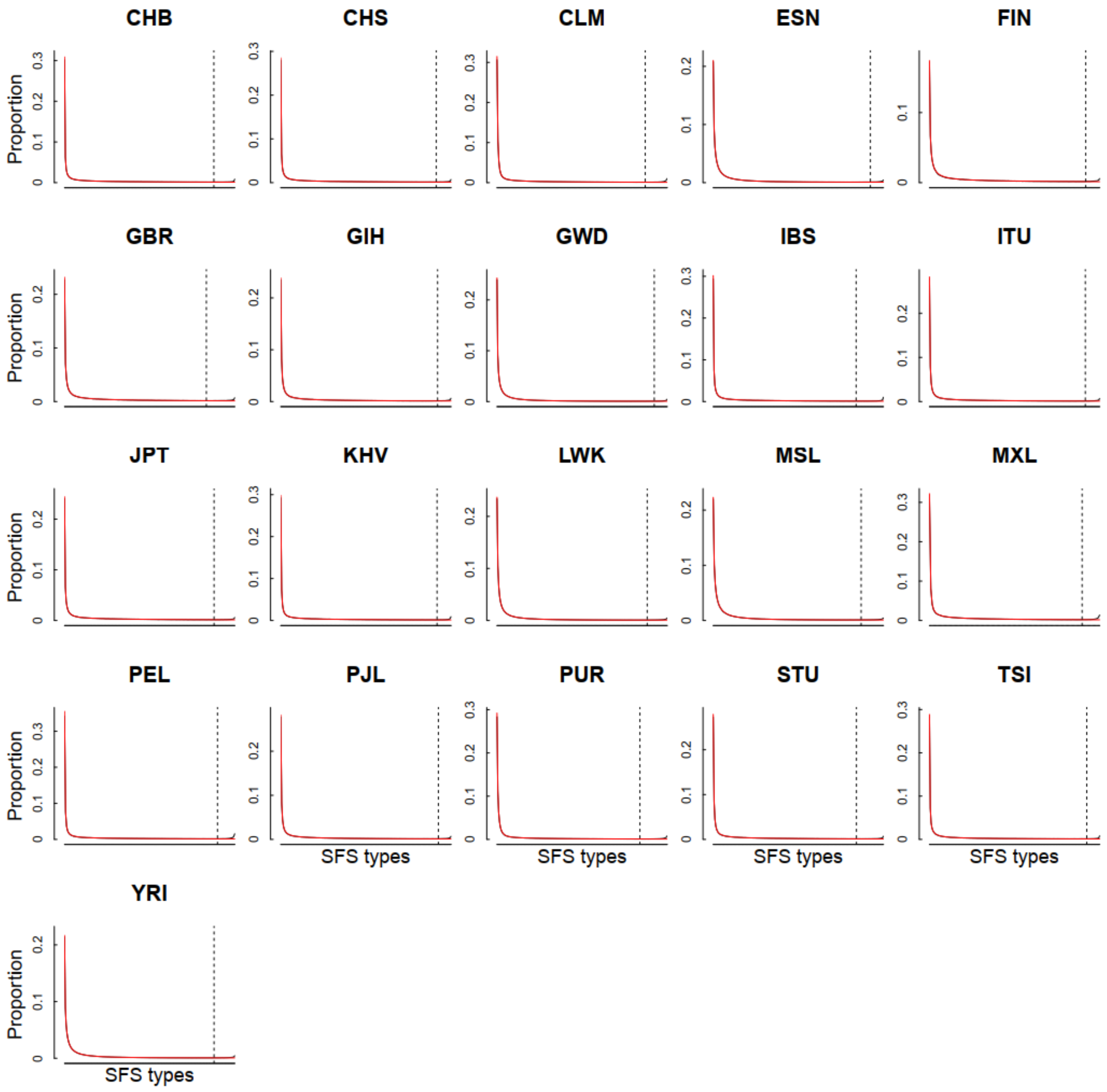

**Figure S20. The observed SFS and simulated SFS of 1000GP populations.** Black solid lines indicate the observed SFS, and the x-axis is the SFS types, ranging from 1 to (n-1). Red solid lines indicate the mean SFS of the 200 data sets simulated under the inferred demographic history for each population. Dash lines indicate the threshold of truncating SFS.

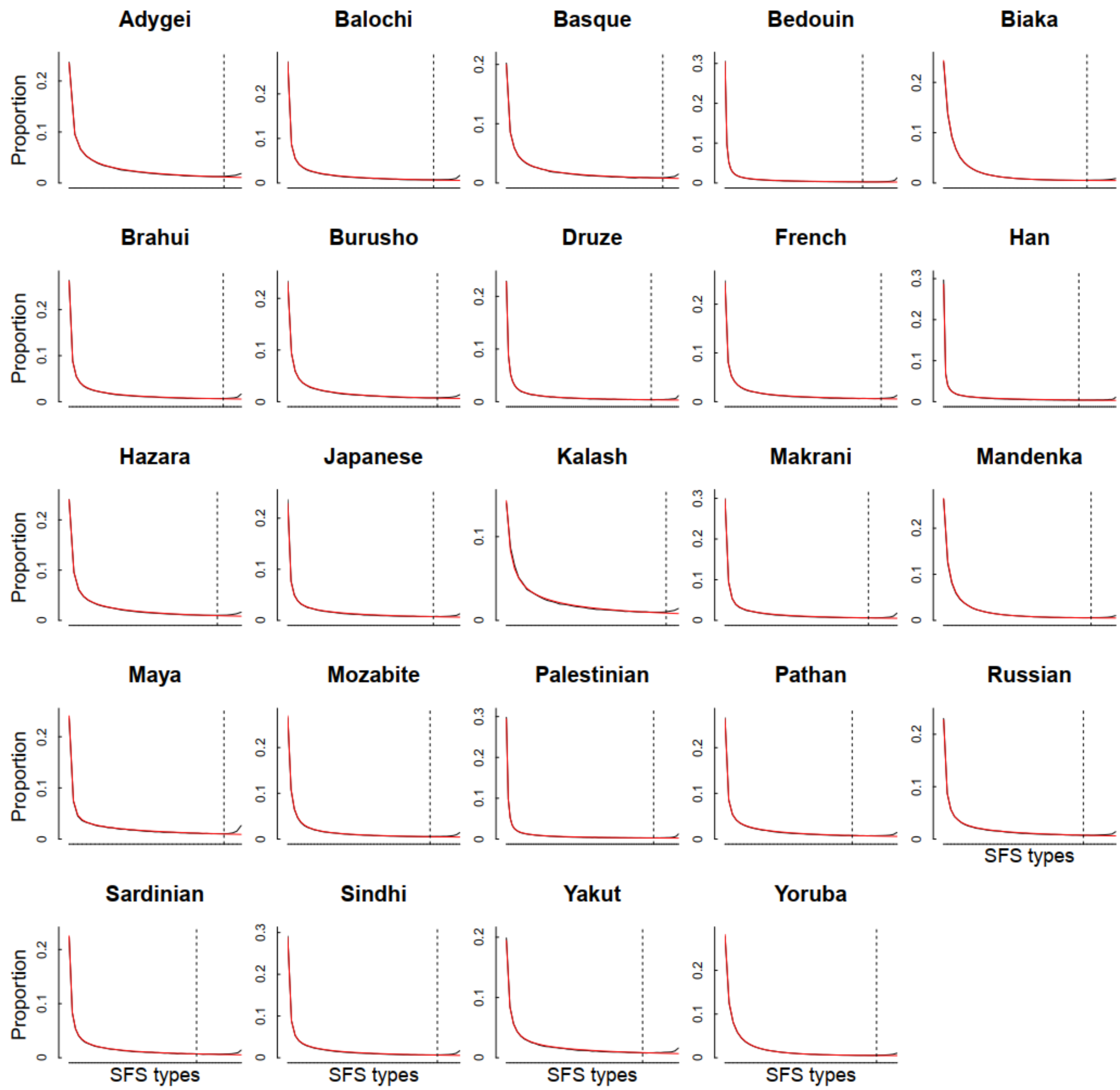

**Figure S21. The observed SFS without missing data and simulated SFS of HGDP-CEPH populations.** Black solid lines indicate the observed SFS, and the x-axis is the SFS types, ranging from 1 to (n-1). Red solid lines indicate the mean SFS of the 200 data sets simulated under the inferred demographic history for each population. Dash lines indicate the threshold of truncating SFS.

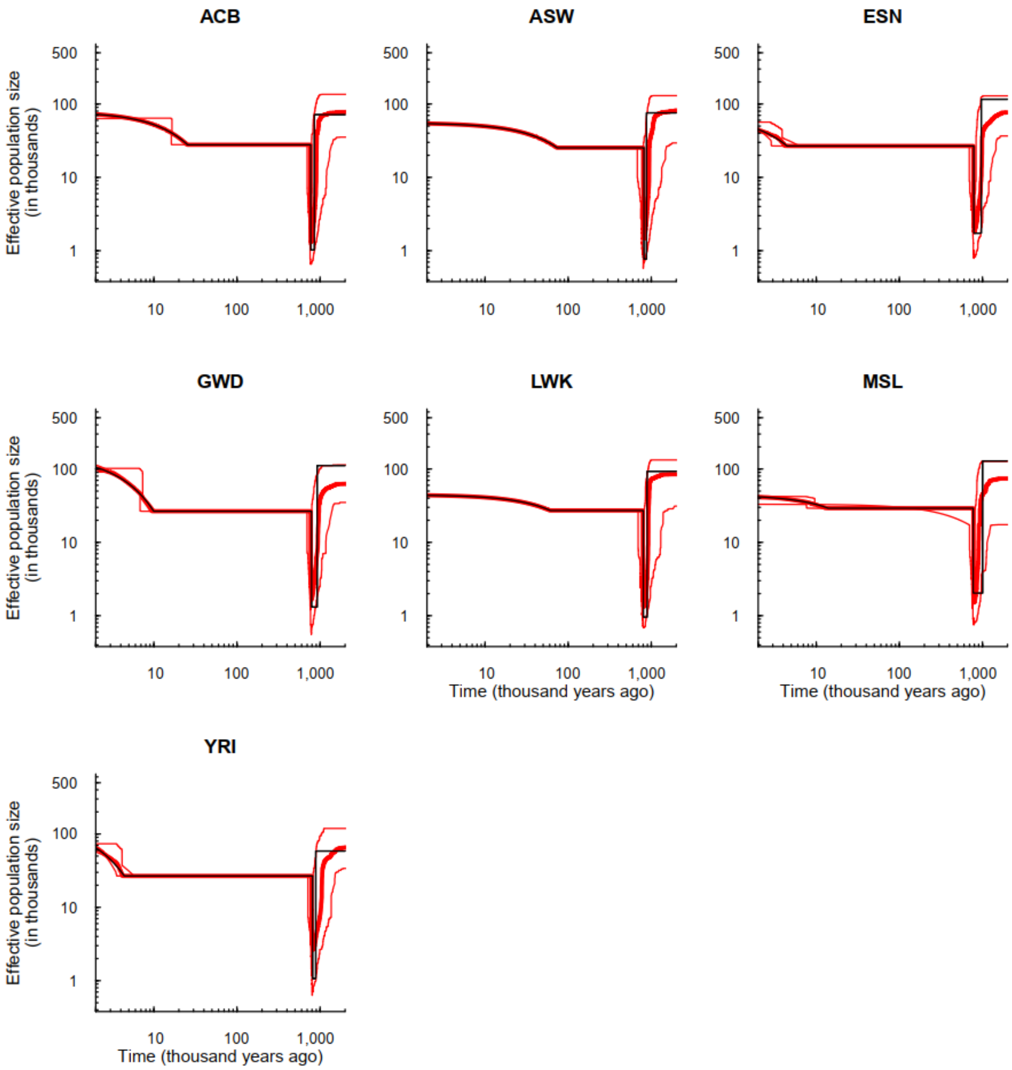

**Figure S22. 95% confidence intervals of 1000GP African populations.** ACB: African Caribbeans in Barbados; ASW: Americans of African Ancestry in SW USA; ESN: Esan in Nigeria; GWD: Gambian in Western Divisions in the Gambia; LWK: Luhya in Webuye, Kenya; MSL: Mende in Sierra Leone; YRI: Yoruba in Ibadan, Nigeria. The inferred demographic histories in Figure 3 are considered as true models, and 200 simulated samples are obtained for each true model. Black lines indicate true models. Thick red lines are the medians of the estimated histories of FitCoal; thin red lines are 2.5 and 97.5 percentiles of the estimated histories of FitCoal. Truncated SFSs were used. The corresponding commands for simulations are described above.

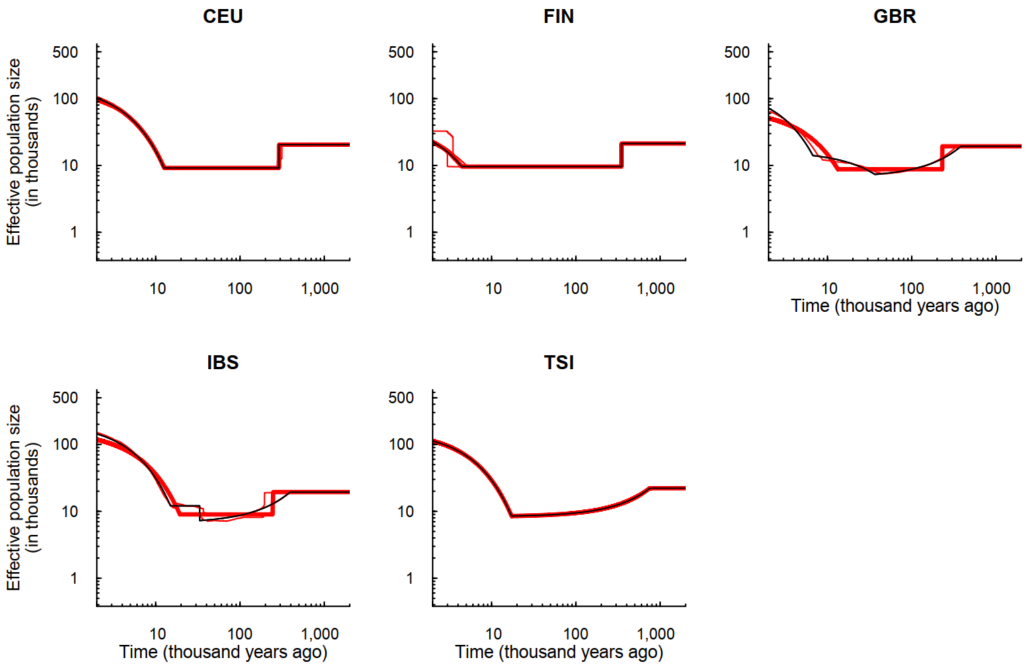

**Figure S23. 95% confidence intervals of 1000GP European populations.** CEU: Utah Residents (CEPH) with Northern and Western European Ancestry; FIN: Finnish in Finland; GBR: British in England and Scotland; IBS: Iberian Population in Spain; TSI: Toscani in Italia. The inferred demographic histories in Figure 3 are considered as true models, and 200 simulated samples are obtained for each true model. Black lines indicate true models. Thick red lines are the medians of the estimated histories of FitCoal; thin red lines are 2.5 and 97.5 percentiles of the estimated histories of FitCoal. Truncated SFSs were used. The corresponding commands for simulations are described above.

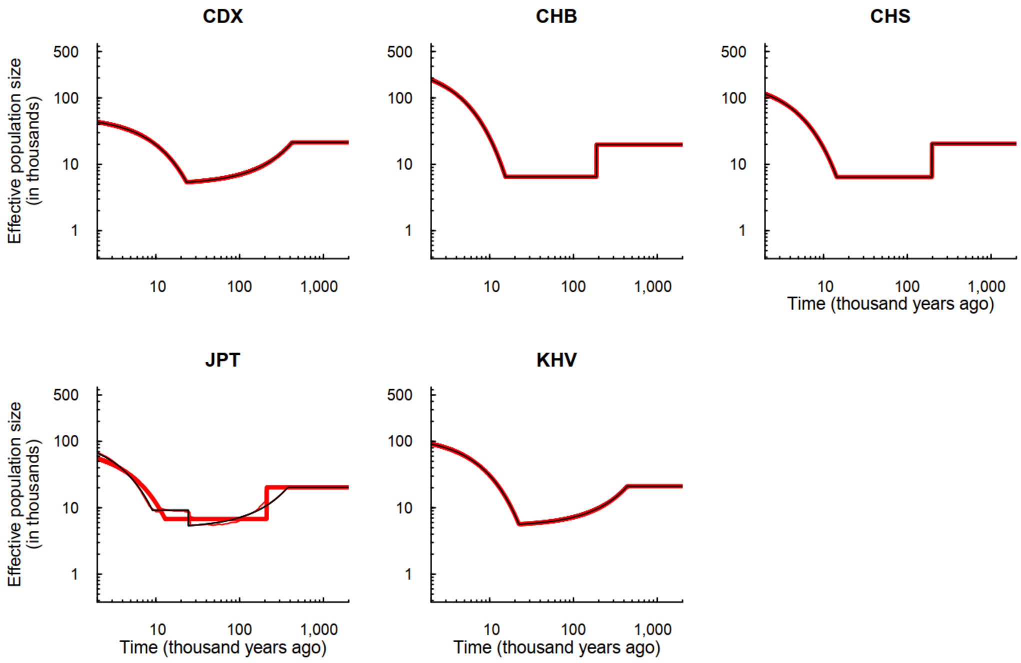

**Figure S24. 95% confidence intervals of 1000GP East Asian populations.** CDX: Chinese Dai in Xishuangbanna, China; CHB: Han Chinese in Beijing, China; CHS: Southern Han Chinese; JPT: Japanese in Tokyo, Japan; KHV: Kinh in Ho Chi Minh City, Vietnam. The inferred demographic histories in Figure 3 are considered as true models, and 200 simulated samples are obtained for each true model. Black lines indicate true models. Thick red lines are the medians of the estimated histories of FitCoal; thin red lines are 2.5 and 97.5 percentiles of the estimated histories of FitCoal. Truncated SFSs were used. The corresponding commands for simulations are described above.

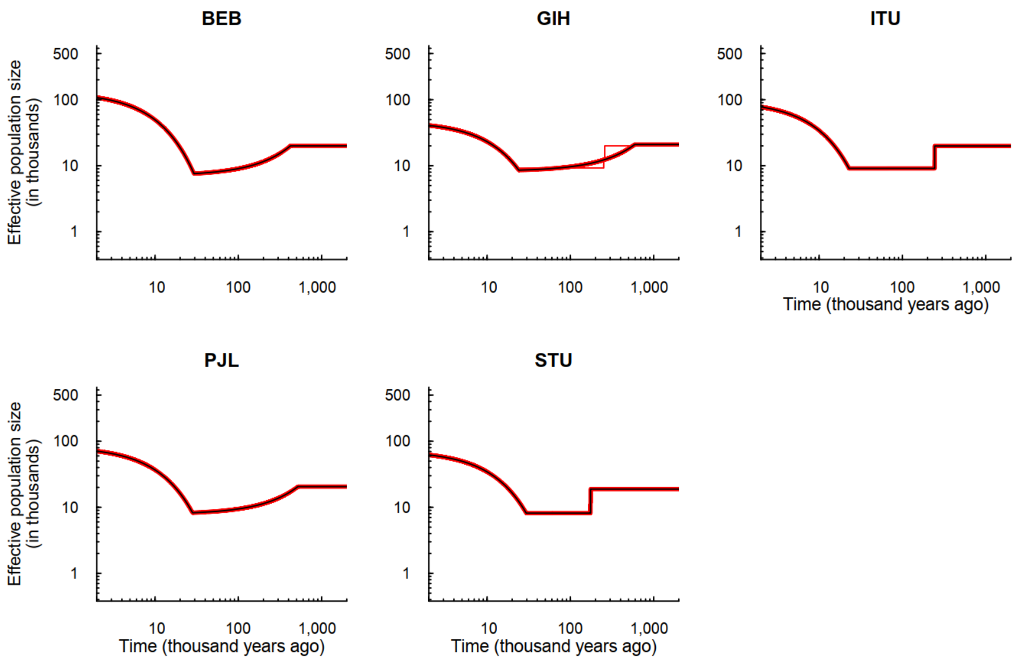

**Figure S25. 95% confidence intervals of 1000GP South Asian populations.** BEB: Bengali from Bangladesh; GIH: Gujarati Indian from Houston, Texas; ITU: Indian Telugu from the UK; PJL: Punjabi from Lahore, Pakistan; STU: Sri Lankan Tamil from the UK. The inferred demographic histories in Figure 3 are considered as true models, and 200 simulated samples are obtained for each true model. Black lines indicate true models. Thick red lines are the medians of the estimated histories of FitCoal; thin red lines are 2.5 and 97.5 percentiles of the estimated histories of FitCoal. Truncated SFSs were used. The corresponding commands for simulations are described above.

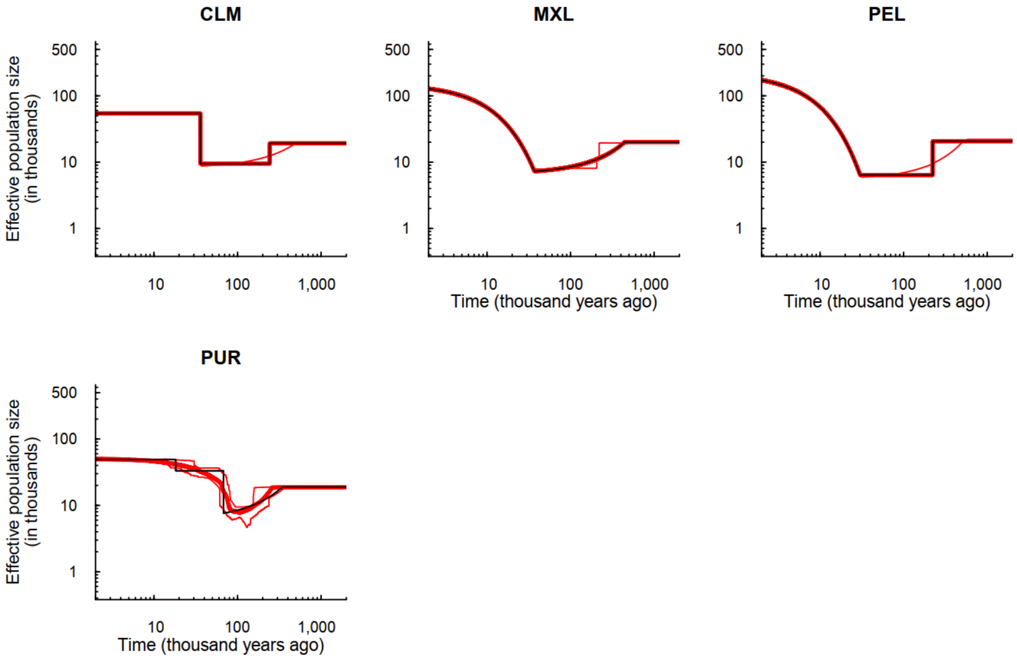

**Figure S26. 95% confidence intervals of 1000GP American populations.** CLM: Colombians from Medellin, Colombia; MXL: Mexican Ancestry from Los Angeles USA; PEL: Peruvians from Lima, Peru; PUR: Puerto Ricans from Puerto Rico. The inferred demographic histories in Figure 3 are considered as true models, and 200 simulated samples are obtained for each true model. Black lines indicate true models. Thick red lines are the medians of the estimated histories of FitCoal; thin red lines are 2.5 and 97.5 percentiles of the estimated histories of FitCoal. Truncated SFSs were used. The corresponding commands for simulations are described above.

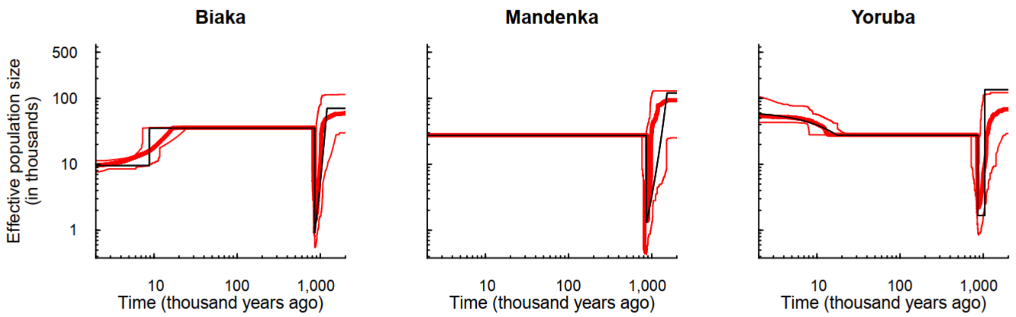

**Figure S27**. **95% confidence intervals of HGPD-CEPH African populations.** The inferred demographic histories in Figure 3 are considered as true models, and 200 simulated samples are obtained for each true model. Black lines indicate true models. Thick red lines are the medians of the estimated histories of FitCoal; thin red lines are 2.5 and 97.5 percentiles of the estimated histories of FitCoal. Truncated SFSs were used. The corresponding commands for simulations are described above.

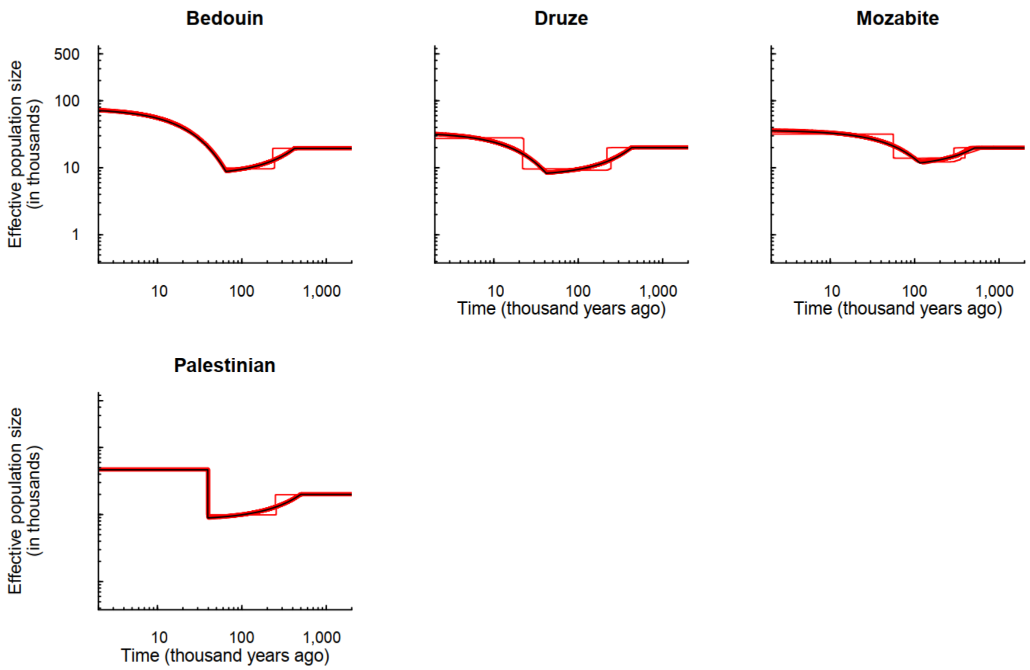

**Figure S28. 95% confidence intervals of HGPD-CEPH Middle East populations.** The inferred demographic histories in Figure 3 are considered as true models, and 200 simulated samples are obtained for each true model. Black lines indicate true models. Thick red lines are the medians of the estimated histories of FitCoal; thin red lines are 2.5 and 97.5 percentiles of the estimated histories of FitCoal. Truncated SFSs were used. The corresponding commands for simulations are described above.

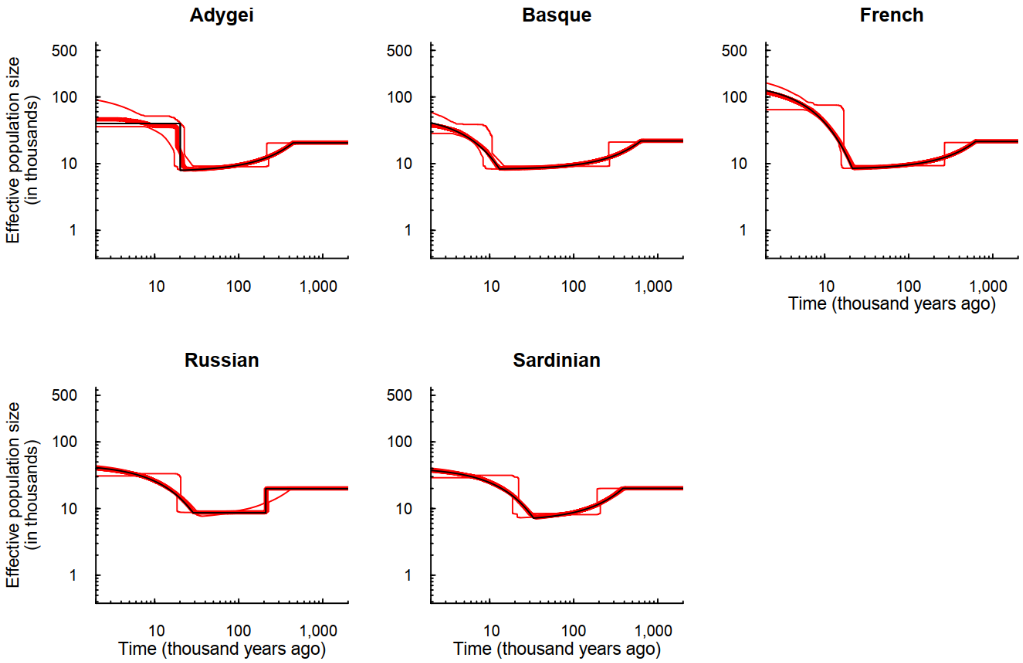

**Figure S29. 95% confidence intervals of HGPD-CEPH European populations.** The inferred demographic histories in Figure 3 are considered as true models, and 200 simulated samples are obtained for each true model. Black lines indicate true models. Thick red lines are the medians of the estimated histories of FitCoal; thin red lines are 2.5 and 97.5 percentiles of the estimated histories of FitCoal. Truncated SFSs were used. The corresponding commands for simulations are described above.

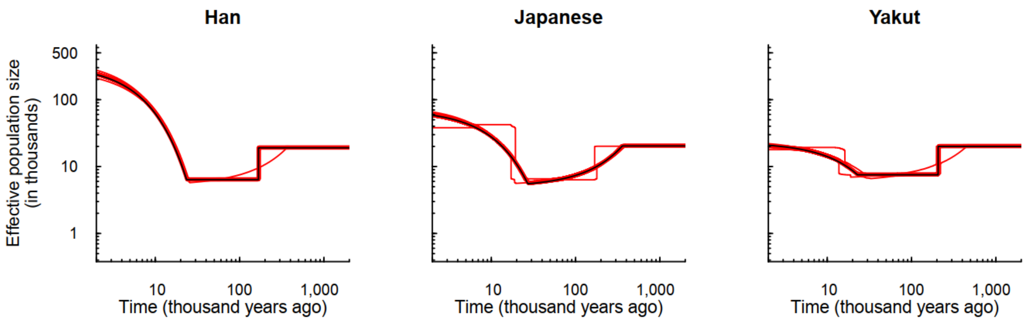

**Figure S30. 95% confidence intervals of HGPD-CEPH East Asian populations.** The inferred demographic histories in Figure 3 are considered as true models, and 200 simulated samples are obtained for each true model. Black lines indicate true models. Thick red lines are the medians of the estimated histories of FitCoal; thin red lines are 2.5 and 97.5 percentiles of the estimated histories of FitCoal. Truncated SFSs were used. The corresponding commands for simulations are described above.

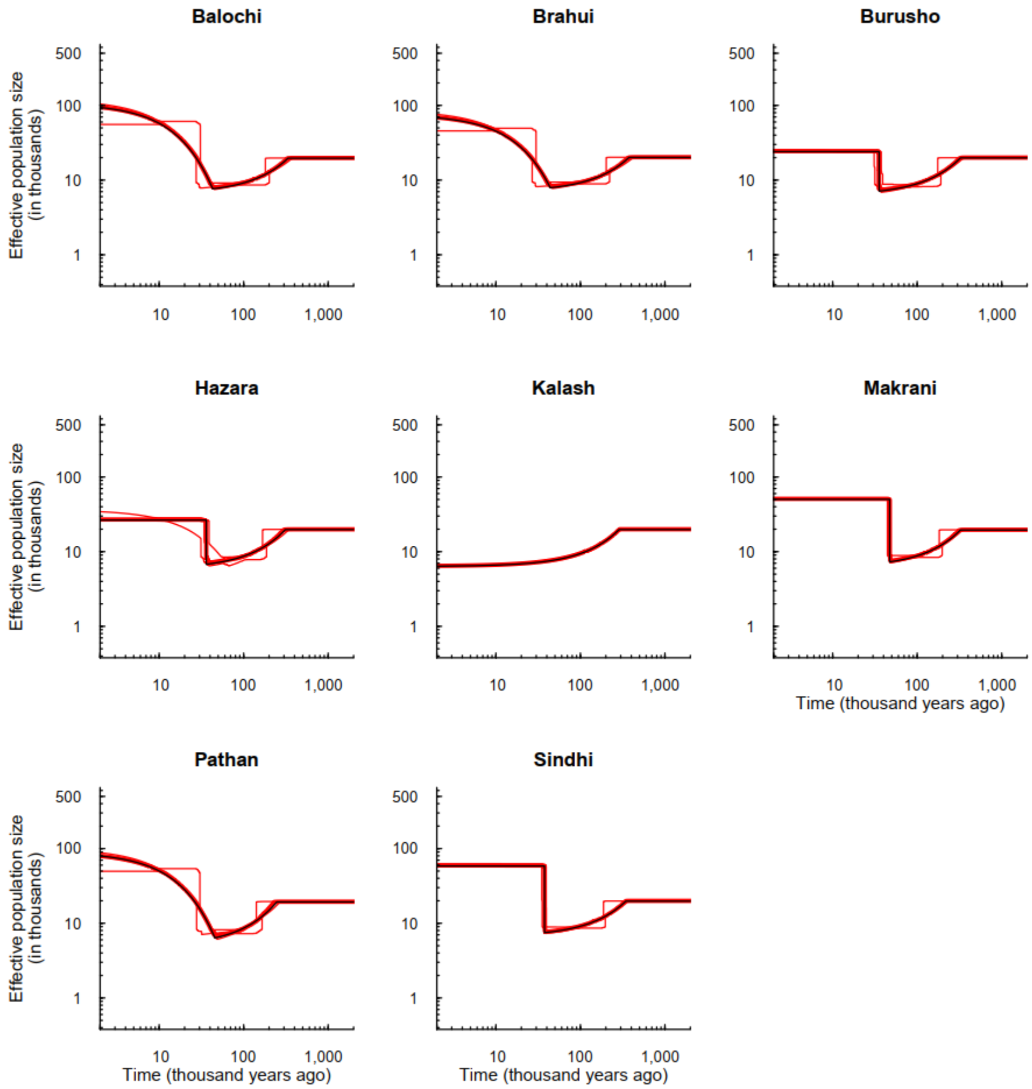

**Figure S31. 95% confidence intervals of HGPD-CEPH Central & South Asian populations.** The inferred demographic histories in Figure 3 are considered as true models, and 200 simulated samples are obtained for each true model. Black lines indicate true models. Thick red lines are the medians of the estimated histories of FitCoal; thin red lines are 2.5 and 97.5 percentiles of the estimated histories of FitCoal. Truncated SFSs were used. The corresponding commands for simulations are described above.

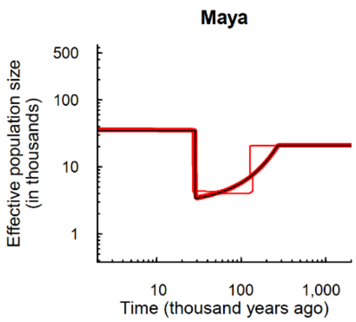

**Figure S32. 95% confidence intervals of HGPD-CEPH American population.** The inferred demographic history in Figure 3 is considered as the true model, and 200 simulated samples are obtained for the true model. Black line indicates the true model. Thick red line is the median of the estimated histories of FitCoal; thin red lines are 2.5 and 97.5 percentiles of the estimated histories of FitCoal. Truncated SFSs were used. The corresponding command for simulations is described above.

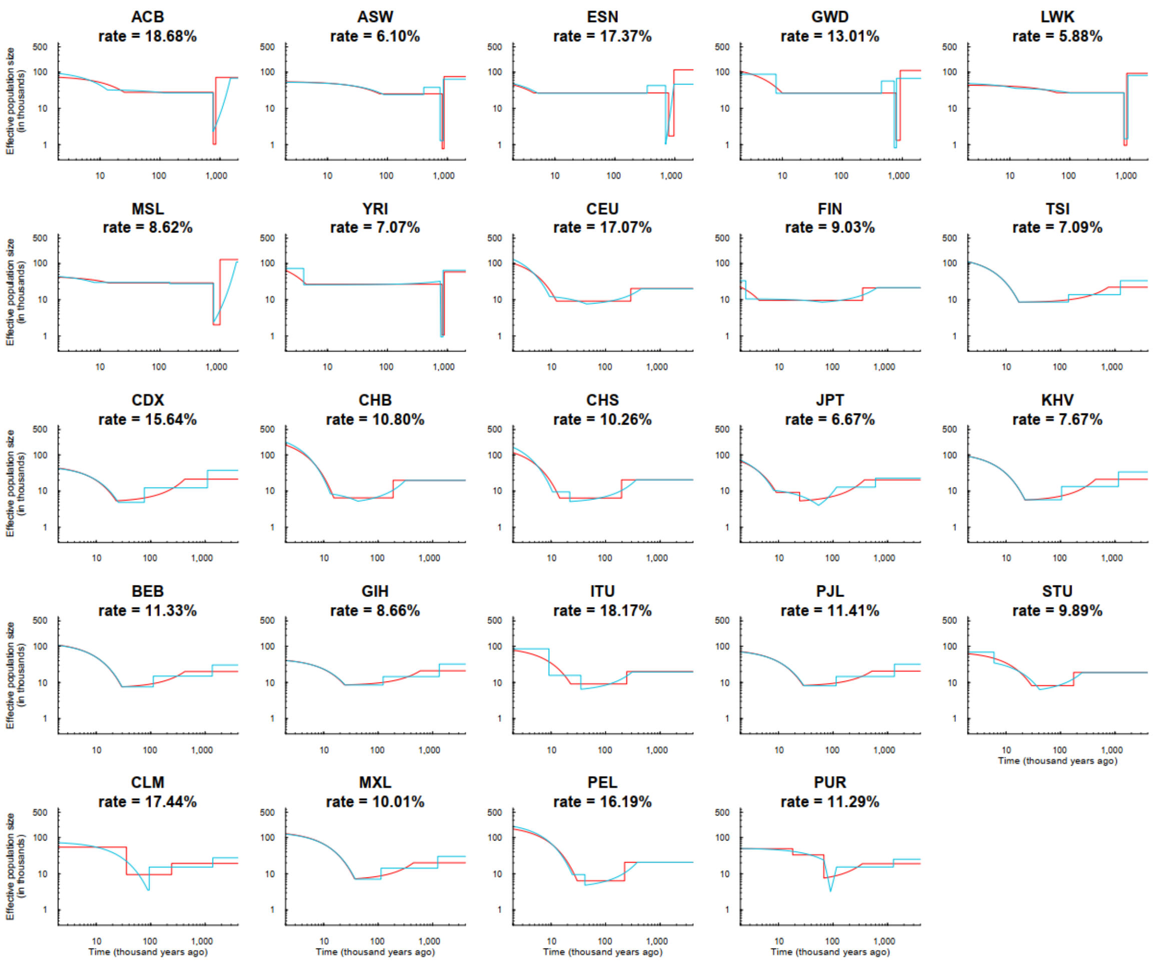

**Figure S33. Inferred demographic histories with different inference time intervals of 1000GP populations.** Red lines indicate the inferred demographic histories, and blue lines indicate demographic histories with one more inference time interval. The log-likelihood promotion rate for the blue-line-indicating history is presented below the name of population. The results of populations are shown in which log-likelihood promotion rate is at least 5% (Table S16).

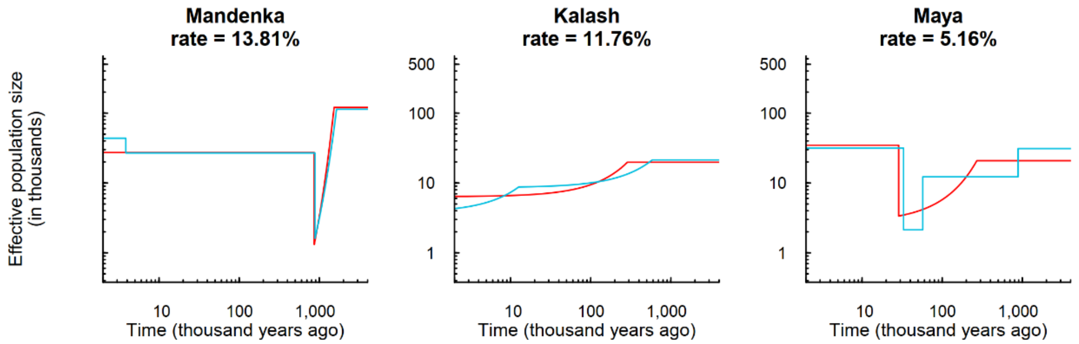

**Figure S34. Inferred demographic histories with different inference time intervals of HGPD-CEPH American populations.** Red lines indicate the inferred demographic histories, and blue lines indicate demographic histories with one more inference time interval. The log-likelihood promotion rate for the blue-line-indicating history is presented below the name of population. The results of populations are shown in which log-likelihood promotion rate is at least 5% (Table S17).

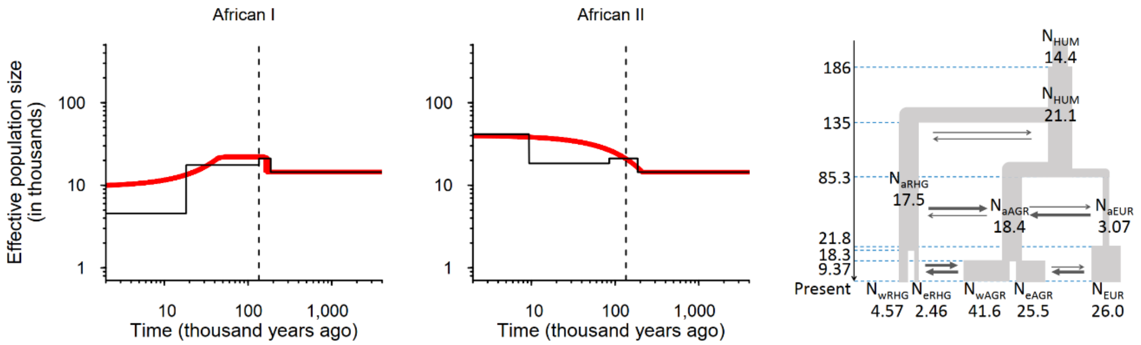

**Figure S35. Verification of inference accuracy under complex population structure model.** We used FitCoal to estimate the demography of imaginary wRHG (African I population) and wAGR (African II population). Split times and population sizes are show in the right panel. The unit of time is 1,000 years and the unit of population size is 1,000. In the left and middle panel, thin solid black lines indicate true models. Thick red lines are the medians of the estimated histories of FitCoal; thin red lines are 2.5 and 97.5 percentiles of the estimated histories of FitCoal. Truncated SFSs of simulated samples were used to infer the history. The corresponding command for simulations is described above. The number of simulated sequences is 170, and the length of simulated sequences is 800 Mb.

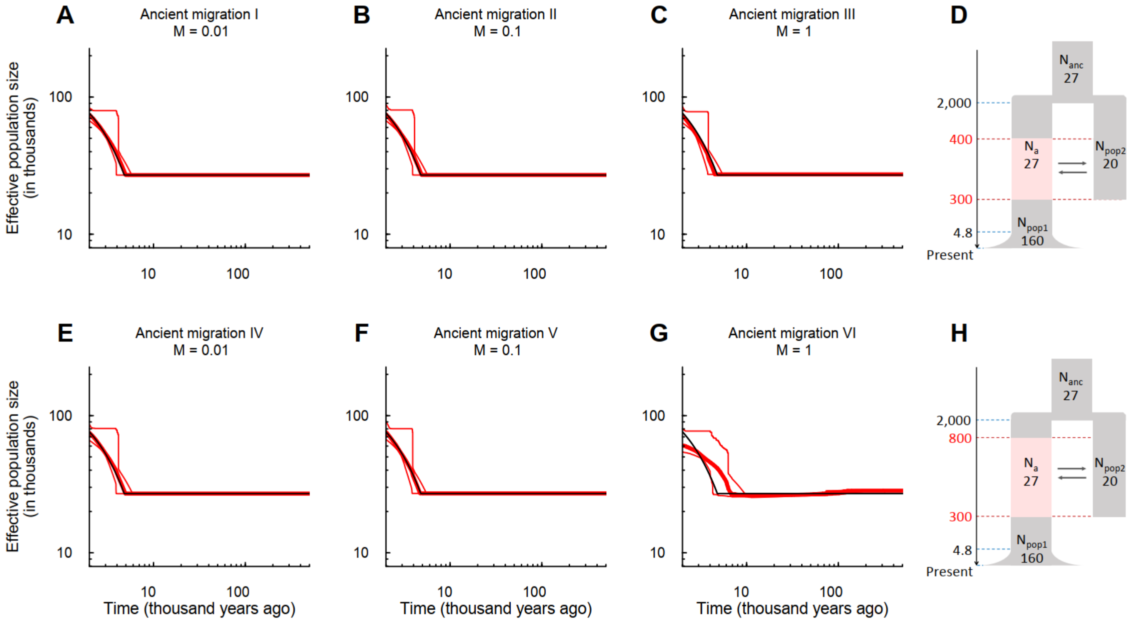

**Figure S36. Verification of inference accuracy when migration occurred with unknown hominin population.** Two models were considered with different time range of migration between the pop1 and an unknown hominin population (**D**, **H**), in which the unit of time is 1,000 years and the unit of population size is 1,000. The region with light red indicates the time range of migration in pop1. We assumed that the unknown population and modern human had common ancestors 2,000 kya. In (**D**), the unknown population began to migrate with modern human 400 kya, and stopped 300 kya. In (**H**), the unknown population began to migrate with modern human 800 kya, and stopped 300 kya. $M$ is the migration rate ($4Nm$). The inferred demographic histories of pop1 in (**D**) are shown in (**A** – **C**), and (**E** – **G**) are results of pop1 in the second model (**H**). The number of simulated sequences is 170, and the length of simulated sequences is 800 Mb. Truncated SFSs were used. Thin solid black lines indicate true models. Thick red lines are the medians of the estimated histories of FitCoal; thin red lines are 2.5 and 97.5 percentiles of the estimated histories of FitCoal.

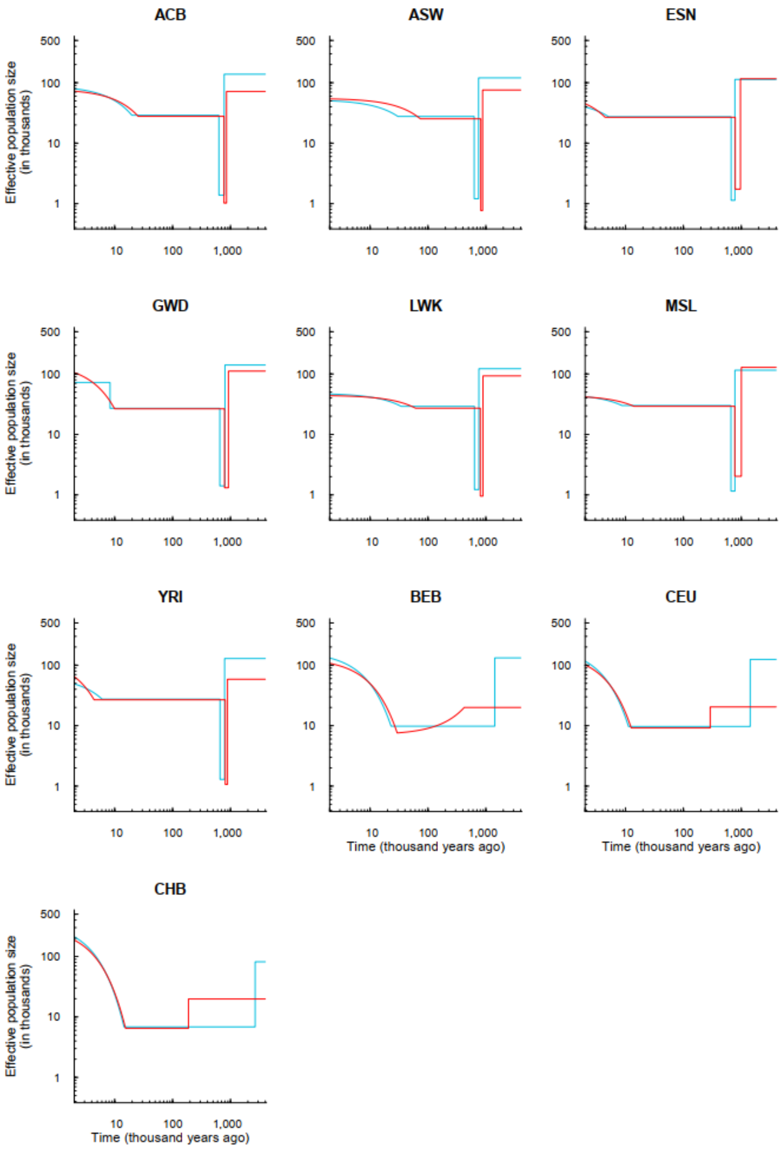

**Figure S37. Estimated demographic histories using the full and the truncated SFSs of 1000GP populations.** ACB: African Caribbeans in Barbados; ASW: Americans of African Ancestry in SW USA; ESN: Esan in Nigeria; GWD: Gambian in Western Divisions in the Gambia; LWK: Luhya in Webuye, Kenya; MSL: Mende in Sierra Leone; YRI: Yoruba in Ibadan, Nigeria, BEB: Bengali from Bangladesh; CEU: Utah Residents (CEPH) with Northern and Western European Ancestry; CHB: Han Chinese in Beijing, China. The comparison is conducted conditional on the same number of inference time intervals. Red lines indicate the demographic histories inferred using the truncated SFSs, the same as what were shown in Figure 3A and S6, and blue lines indicate these inferred using the full SFS.

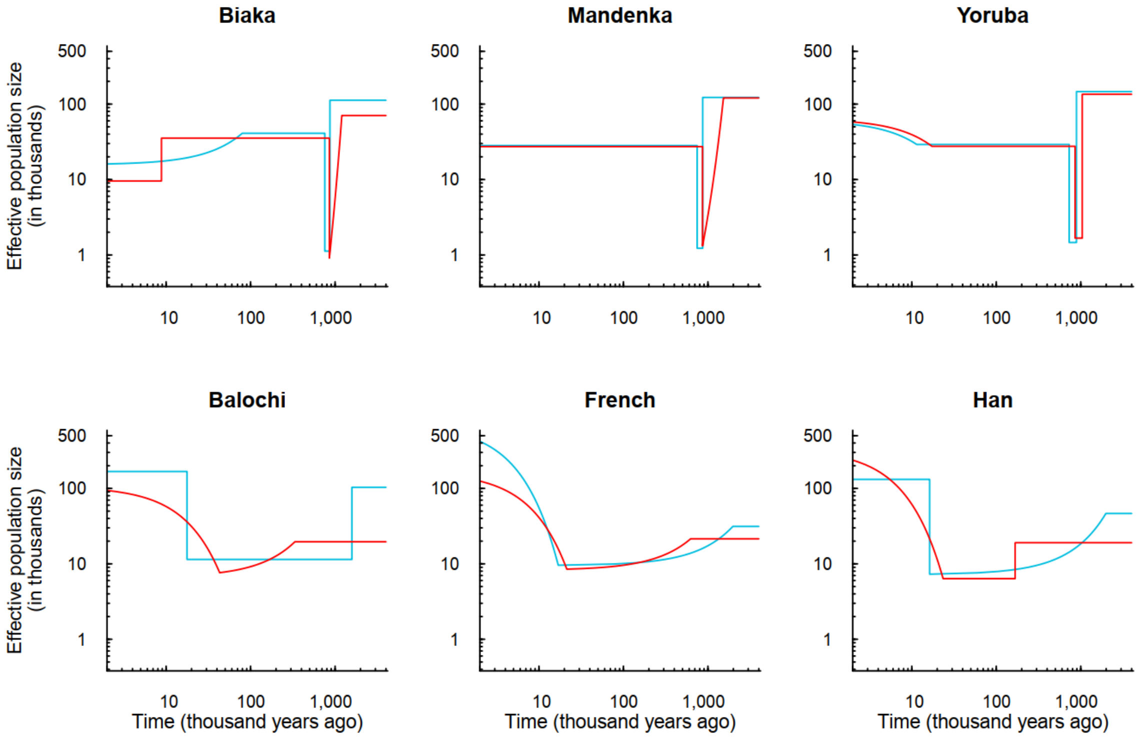

**Figure S38. Estimated demographic histories using the full and the truncated SFSs of HGDP-CEPH populations.** The comparison is conducted conditional on the same number of inference time intervals. Red lines indicate the demographic histories inferred using the truncated SFSs, the same as what were shown in Figure 3D and S7, and blue lines indicate these inferred using the full SFSs.

### Supplementary tables

**Table S1. Proportion of correctly-inferred change type for the most recent demographic event in six models.**

| Model | Proportion |
|---|---|
| Constant size | 100% |
| Instantaneous increase | 85.5% |
| PSMC “standard” | 100% |
| Exponential growth I | 41.5% |
| Exponential growth II | 61% |
| Exponential growth III | 66.5% |

**Table S2. Parameters of the super bottleneck in 1000GP African populations.**

| Population | Ancestral Ne | Start time of the bottleneck | | Ne during the bottleneck | End time of the bottleneck | | Ne immediate. after the bottleneck |
|---|---|---|---|---|---|---|---|
| | | Time[a] | Change type[b] | | Time[a] | Change type[b] | |
| ACB | 71,705 | 854,288 | I | 1,021 | 772,422 | I | 27,802 |
| ASW | 75,746 | 877,763 | I | 767 | 815,473 | I | 25,302 |
| ESN | 116,216 | 966,439 | I | 1,735 | 785,741 | I | 26,785 |
| GWD | 111,486 | 922,296 | I | 1,311 | 790,048 | I | 26,546 |
| LWK | 92,952 | 891,086 | I | 954 | 802,498 | I | 27,142 |
| MSL | 128,666 | 1,002,553 | I | 2,031 | 773,633 | I | 29,183 |
| YRI | 58,563 | 887,367 | I | 1,066 | 812,636 | I | 26,796 |

**Note**: a: Time in years. b: I represents instantaneous change, and E represents exponential change.

**Table S3. Parameters of the super bottleneck in HGDP-CEPH African populations.**

| Population | Ancestral Ne | Start time of the bottleneck | | Ne during the bottleneck | End time of the bottleneck | | Ne immediately after the bottleneck |
|---|---|---|---|---|---|---|---|
| | | Time[a] | Change type[b] | | Time[a] | Change type[b] | |
| Biaka | 70,583 | 1,202,743 | E | 908 | 855,778 | I | 35,329 |
| Mandenka | 120,452 | 1,526,972 | E | 1,319 | 864,078 | I | 27,306 |
| Yoruba | 134,958 | 1,041,950 | I | 1,670 | 856,750 | I | 27,569 |

**Note**: a: Time in years. b: I represents instantaneous change, and E represents exponential change.

**Table S4. Parameters of the out-of-Africa bottleneck in 1000GP non-African populations.**

| Super population | Population | Ancestral Ne | Start time of the bottleneck | | Ne during the bottleneck |
|---|---|---|---|---|---|
| | | | Time[a] | Change type[b] | |
| EUR | CEU | 20,414 | 290,251 | I | 9,168 |
| | GBR | 19,334 | 383,068 | E | 7,343 |
| | FIN | 21,278 | 348,509 | I | 9,612 |
| | IBS | 19,393 | 391,678 | E | 7,270 |
| | TSI | 22,215 | 756,179 | E | 8,505 |
| EAS | CDX | 20,414 | 421,701 | I | 9,168 |
| | CHB | 19,334 | 188,134 | E | 7,343 |
| | CHS | 21,278 | 196,219 | I | 9,612 |
| | JPT | 19,393 | 377,347 | E | 7,270 |
| | KHV | 22,215 | 439,168 | E | 8,505 |
| SAS | BEB | 19,998 | 422,436 | E | 7,584 |
| | GIH | 20,941 | 591,769 | E | 8,554 |
| | ITU | 19,912 | 244,308 | I | 9,091 |
| | PJL | 20,556 | 518,059 | E | 8,257 |
| | STU | 18,848 | 174,620 | I | 8,157 |
| AMR | CLM | 19,252 | 242,526 | I | 9,482 |
| | MXL | 20,063 | 444,463 | E | 7,266 |
| | PEL | 20,666 | 222,966 | I | 6,384 |
| | PUR | 18,917 | 344,023 | E | 7,606 |

**Note**: a: Time in years. b: I represents instantaneous change, and E represents exponential change.

**Table S5. Parameters of the out-of-Africa bottleneck in HGDP-CEPH non-African populations.**

| Super population | Population | Ancestral Ne | Start time of the bottleneck | | Ne during the bottleneck |
|---|---|---|---|---|---|
| | | | Time[a] | Change type[b] | |
| Middle East | Bedouin | 19,410 | 420,348 | E | 8,793 |
| | Druze | 19,930 | 428,504 | E | 8,275 |
| | Mozabite | 19,932 | 497,224 | E | 8,904 |
| | Palestinian | 19,733 | 513,178 | E | 11,795 |
| European | Adygei | 20,467 | 443,402 | E | 7,934 |
| | Basque | 21,848 | 627,518 | E | 8,315 |
| | French | 21,469 | 623,958 | E | 8,496 |
| | Russian | 19,836 | 209,860 | I | 8,647 |
| | Sardinian | 20,065 | 385,155 | E | 7,183 |
| East Asian | Han | 19,055 | 167,289 | I | 6,365 |
| | Japanese | 20,303 | 357,997 | E | 5,532 |
| | Yakut | 20,012 | 206,684 | I | 7,526 |
| Central & South Asian | Balochi | 19,618 | 334,493 | E | 7,635 |
| | Brahui | 20,095 | 375,825 | E | 7,897 |
| | Burusho | 19,846 | 325,186 | E | 7,127 |
| | Hazara | 19,765 | 310,990 | E | 6,780 |
| | Kalash | 19,766 | 286,505 | E | 6,346 |
| | Makrani | 19,497 | 324,241 | E | 7,297 |
| | Pathan | 19,283 | 245,899 | E | 6,364 |
| | Sindhi | 19,758 | 344,223 | E | 7,493 |
| American | Maya | 20,847 | 270,478 | E | 3,365 |

**Note**: a: Time in years. b: I represents instantaneous change, and E represents exponential change.

**Table S6. The super bottleneck parameters of Bottleneck I model.**

| Parameter | Value | Median of estimations | Lower bound of 95% CI | Upper bound of 95% CI |
|---|---|---|---|---|
| Start time of the bottleneck | 912,000 | 978,024 | 790,776 | 1,890,528 |
| End time of the bottleneck | 792,000 | 764,376 | 685,656 | 794,784 |
| Population size before the bottleneck | 93,000 | 62,224 | 32,334 | 130,445 |
| Population size during the bottleneck | 1,300 | 1,907 | 546 | 5,838 |
| Population size after the bottleneck | 27,000 | 27,069 | 26,936 | 27,230 |

**Table S7. The super bottleneck parameters of Bottleneck IV model.**

| Parameter | Value | Median of estimations | Lower bound of 95% CI | Upper bound of 95% CI |
|---|---|---|---|---|
| Start time of the bottleneck | 984,000 | 1,072,368 | 894,648 | 1,899,288 |
| End time of the bottleneck | 864,000 | 854,256 | 774,360 | 873,672 |
| Population size before the bottleneck | 110,000 | 72,408 | 31,934 | 140,463 |
| Population size during the bottleneck | 1,300 | 1,677 | 638 | 5,536 |
| Population size after the bottleneck | 28,000 | 27,965 | 27,817 | 28,203 |

**Table S8. The super bottleneck parameters of Bottleneck VII model.**

| Parameter | Value | Median of estimations | Lower bound of 95% CI | Upper bound of 95% CI |
|---|---|---|---|---|
| Start time of the bottleneck | 840,000 | 887,088 | 736,560 | 1,419,480 |
| End time of the bottleneck | 720,000 | 719,712 | 650,688 | 1,126,176 |
| Population size before the bottleneck | 30,000 | 31,513 | 25,356 | 132,364 |
| Population size during the bottleneck | 3,000 | 2,892 | 328 | 7,682 |
| Population size after the bottleneck | 30,000 | 3,0002 | 29,924 | 30,086 |

**Table S9. Influence of different log-likelihood promotion thresholds.**

| Model | 30% | | | 20% | | | 10% | | |
|---|---|---|---|---|---|---|---|---|---|
| | Underfitting | Correct | Overfitting | Underfitting | Correct | Overfitting | Underfitting | Correct | Overfitting |
| Constant size | … | 100% | 0 | … | 100% | 0 | … | 93% | 2% |
| Instantaneous increase | 0 | 100% | 0 | 0 | 100% | 0 | 0 | 99.5% | 0.5% |
| PSMC “standard” | 0 | 100% | 0 | 0 | 100% | 0 | 0 | 100% | 0 |
| Exponential growth II | 0 | 100% | 0 | 0 | 100% | 0 | 0 | 99.5% | 0.5% |
| Exponential growth III | 0 | 100% | 0 | 0 | 100% | 0 | 0 | 100% | 0 |
| PSMC sim-YH | 0 | 100% | 0 | 0 | 100% | 0 | 0 | 99.5% | 0.5% |
| PSMC sim-1 | 0 | 100% | 0 | 0 | 100% | 0 | 0 | 100% | 0 |
| PSMC sim-2 | 20% | 80% | 0 | 0 | 100% | 0 | 0 | 100% | 0 |
| PSMC sim-3 | 0 | 100% | 0 | 0 | 100% | 0 | 0 | 98% | 2% |
| Complicated I | 0 | 100% | 0 | 0 | 100% | 0 | 0 | 100% | 0 |
| Complicated II | 0 | 100% | 0 | 0 | 100% | 0 | 0 | 100% | 0 |
| Exponential growth IV | 0 | 100% | 0 | 0 | 100% | 0 | 0 | 100% | 0 |
| Exponential growth V | 0 | 100% | 0 | 0 | 100% | 0 | 0 | 100% | 0 |
| Exponential growth VI | 0 | 100% | 0 | 0 | 100% | 0 | 0 | 100% | 0 |
| Split I pop 1 | 0 | 100% | 0 | 0 | 100% | 0 | 0 | 99.5% | 0.5% |
| Split I pop 2 | 0 | 100% | 0 | 0 | 100% | 0 | 0 | 99.5% | 0.5% |
| Split II pop 1 | 0 | 100% | 0 | 0 | 100% | 0 | 0 | 100% | 0 |
| Split II pop 2 | 0 | 100% | 0 | 0 | 100% | 0 | 0 | 100% | 0 |
| Split III pop 1 | 0 | 100% | 0 | 0 | 100% | 0 | 0 | 100% | 0 |
| Split III pop 2 | 0 | 100% | 0 | 0 | 100% | 0 | 0 | 100% | 0 |

**Table S10. Information of truncated SFS of 1000GP populations.**

| Population | Size of SFS ($n-1$) | Truncated sizes of SFS | Proportion of truncated sizes (%) | Proportion of truncated SNPs (%) |
|---|---|---|---|---|
| ACB | 191 | 171 – 191 | 10.99 | 3.17 |
| ASW | 121 | 107 – 121 | 12.40 | 3.66 |
| BEB | 171 | 156 – 171 | 9.36 | 4.00 |
| CDX | 185 | 174 – 185 | 6.49 | 3.56 |
| CEU | 197 | 178 – 197 | 10.15 | 4.39 |
| CHB | 205 | 180 – 205 | 12.68 | 5.34 |
| CHS | 209 | 191 – 209 | 9.09 | 4.17 |
| CLM | 187 | 163 – 187 | 13.37 | 5.49 |
| ESN | 197 | 182 – 197 | 8.12 | 2.67 |
| FIN | 197 | 181 – 197 | 8.63 | 4.19 |
| GBR | 181 | 151 – 181 | 17.13 | 6.60 |
| GIH | 205 | 189 – 205 | 8.29 | 3.75 |
| GWD | 225 | 208 – 225 | 8.00 | 2.59 |
| IBS | 213 | 179 – 213 | 16.43 | 6.29 |
| ITU | 203 | 186 – 203 | 8.87 | 3.88 |
| JPT | 207 | 182 – 207 | 12.56 | 5.26 |
| KHV | 197 | 181 – 197 | 8.63 | 4.21 |
| LWK | 197 | 174 – 197 | 12.18 | 3.39 |
| MSL | 169 | 147 – 169 | 13.61 | 3.63 |
| MXL | 127 | 114 – 127 | 11.02 | 5.49 |
| PEL | 169 | 152 – 169 | 10.65 | 6.21 |
| PJL | 191 | 177 – 191 | 7.85 | 3.54 |
| PUR | 207 | 174 – 207 | 16.43 | 5.77 |
| STU | 203 | 171 – 203 | 16.26 | 5.79 |
| TSI | 213 | 197 – 213 | 7.98 | 3.78 |
| YRI | 215 | 189 – 215 | 12.56 | 3.50 |

**Table S11. Proportion of SNPs without or with missing data of HGDP-CEPH populations.**

| Population | No missing data (%) | Missing samples <= 2 chromosomes (%) |
|---|---|---|
| Adygei | 92.58 | 98.80 |
| Balochi | 91.01 | 98.37 |
| Basque | 92.66 | 98.83 |
| Bedouin | 87.00 | 97.30 |
| Biaka | 93.25 | 98.03 |
| Brahui | 90.76 | 98.37 |
| Burusho | 91.30 | 98.55 |
| Druze | 89.60 | 97.86 |
| French | 88.78 | 98.14 |
| Han | 84.54 | 97.14 |
| Hazara | 92.53 | 98.76 |
| Japanese | 87.45 | 97.96 |
| Kalash | 90.57 | 98.48 |
| Makrani | 91.10 | 98.29 |
| Mandenka | 91.93 | 98.13 |
| Maya | 94.98 | 99.13 |
| Mozabite | 93.21 | 98.58 |
| Palestinian | 87.22 | 97.32 |
| Pathan | 91.74 | 98.60 |
| Russian | 92.68 | 98.68 |
| Sardinian | 93.04 | 98.81 |
| Sindhi | 90.73 | 98.43 |
| Yakut | 86.66 | 97.80 |
| Yoruba | 92.78 | 98.02 |

**Table S12. Information of truncated SFS of HGDP-CEPH populations.**

| Population | Size of SFS ($n-1$) | Truncated sizes of SFS | Proportion of truncated sizes (%) | Proportion of truncated SNPs (%) |
|---|---|---|---|---|
| Adygei | 31 | 28 – 31 | 12.90 | 5.65 |
| Balochi | 47 | 40 – 47 | 17.02 | 7.22 |
| Basque | 45 | 41 – 45 | 11.11 | 6.18 |
| Bedouin | 91 | 73 – 91 | 20.88 | 9.52 |
| Biaka | 43 | 36 – 43 | 18.60 | 4.93 |
| Brahui | 49 | 44 – 49 | 12.24 | 5.21 |
| Burusho | 47 | 41 – 47 | 14.89 | 6.57 |
| Druze | 83 | 70 – 83 | 16.87 | 8.24 |
| French | 55 | 50 – 56 | 10.91 | 5.59 |
| Han | 85 | 67 – 85 | 22.35 | 8.72 |
| Hazara | 37 | 32 – 37 | 16.22 | 6.72 |
| Japanese | 53 | 45 – 53 | 16.98 | 6.95 |
| Kalash | 43 | 40 – 43 | 9.30 | 5.22 |
| Makrani | 49 | 41 – 49 | 18.37 | 7.24 |
| Mandenka | 43 | 37 – 43 | 16.28 | 4.75 |
| Maya | 41 | 37 – 41 | 12.20 | 7.20 |
| Mozabite | 53 | 44 – 53 | 18.87 | 7.43 |
| Palestinian | 91 | 78 – 91 | 15.38 | 8.53 |
| Pathan | 47 | 35 – 49 | 27.66 | 10.12 |
| Russian | 49 | 40 – 49 | 20.41 | 8.06 |
| Sardinian | 55 | 51 – 55 | 27.27 | 11.65 |
| Sindhi | 47 | 41 – 47 | 14.89 | 6.35 |
| Yakut | 49 | 39 – 49 | 22.45 | 10.02 |
| Yoruba | 43 | 38 – 43 | 13.95 | 4.14 |

**Table S13. Comparison of branch length between theoretical values and FitCoal under the constant size model.**

| | $n = 5$ | |
|---|---|---|
| Type | Theoretical length | FitCoal length |
| 1 | 2.000000000000 | 1.999999995001 |
| 2 | 1.000000000000 | 0.999999995001 |
| 3 | 0.666666666667 | 0.666666661667 |
| 4 | 0.500000000000 | 0. 499999995000 |
| | $n = 1{,}000$ | |
| Type | Theoretical length | FitCoal length |
| 1 | 2.000000000000 | 1.999999999979 |
| 100 | 0.020000000000 | 0.019999999980 |
| 200 | 0.010000000000 | 0.009999999980 |
| 300 | 0.006666666667 | 0.006666666647 |
| 400 | 0.005000000000 | 0.004999999980 |
| 500 | 0.004000000000 | 0.003999999980 |
| 600 | 0.003333333333 | 0.003333333313 |
| 700 | 0.002857142857 | 0.002857142837 |
| 800 | 0.002500000000 | 0.002499999980 |
| 900 | 0.002222222222 | 0.002222222202 |
| 999 | 0.002002002002 | 0.002002001982 |

**Table S14. Comparison of accuracy between FitCoal and Z-W's method.**

| Constant size model | | | |
|---|---|---|---|
| size | Z-W's method | FitCoal | Tabulated FitCoal |
| 1 | 1.9999999999 | 1.9999999977 | 2.0000000000 |
| 2 | 0.9999999999 | 0.9999999977 | 1.0000000000 |
| 3 | 0.6666666666 | 0.6666666644 | 0.6666666666 |
| 4 | 0.4999999999 | 0.4999999977 | 0.5000000000 |
| 5 | 0.3999999999 | 0.3999999977 | 0.4000000000 |
| 6 | 0.3333333333 | 0.3333333311 | 0.3333333333 |
| 7 | 0.2857142857 | 0.2857142834 | 0.2857142857 |
| 8 | 0.2499999999 | 0.2499999977 | 0.2500000000 |
| 9 | 0.2222222222 | 0.2222222200 | 0.2222222222 |
| Instantaneous growth model | | | |
| size | Z-W's method | FitCoal | Tabulated FitCoal |
| 1 | 1.7547171613 | 1.7547178684 | 1.7547229380 |
| 2 | 0.7797340922 | 0.7797345132 | 0.7797375358 |
| 3 | 0.4686921135 | 0.4686923318 | 0.4686939043 |
| 4 | 0.3218480766 | 0.3218481580 | 0.3218487511 |
| 5 | 0.2394397929 | 0.2394397884 | 0.2394397658 |
| 6 | 0.1883537444 | 0.1883536920 | 0.1883533260 |
| 7 | 0.1545070379 | 0.1545069652 | 0.1545064537 |
| 8 | 0.1309436073 | 0.1309435332 | 0.1309430117 |
| 9 | 0.1138668845 | 0.1138668210 | 0.1138663751 |
| Bottleneck model | | | |
| size | Z-'s method | FitCoal | FitCoal with tabulated |
| 1 | 1.6737287481 | 1.6737298163 | 1.6737374623 |
| 2 | 0.7744529188 | 0.7744529447 | 0.7744538197 |
| 3 | 0.5109730563 | 0.5109727763 | 0.5109715354 |
| 4 | 0.3927621211 | 0.3927618350 | 0.3927603900 |
| 5 | 0.3264269184 | 0.3264267194 | 0.3264256821 |
| 6 | 0.2832102142 | 0.2832101093 | 0.2832095492 |
| 7 | 0.2519772704 | 0.2519772388 | 0.2519770488 |
| 8 | 0.2277455013 | 0.2277455192 | 0.2277455704 |
| 9 | 0.2080244642 | 0.2080245126 | 0.2080247051 |

**Table S15. Comparison of accuracy between FitCoal and simulations.**

| Constant size model | | | |
|---|---|---|---|
| Type | Simulation method | FitCoal | Tabulated FitCoal |
| 1 | 1.999529551 | 1.999999998 | 2.000000000 |
| 2 | 0.999720720 | 0.999999998 | 1.000000000 |
| 3 | 0.667102715 | 0.666666664 | 0.666666667 |
| 4 | 0.500240165 | 0.499999998 | 0.500000000 |
| 5 | 0.400967708 | 0.399999998 | 0.400000000 |
| 6 | 0.333256388 | 0.333333331 | 0.333333333 |
| 7 | 0.285465389 | 0.285714283 | 0.285714286 |
| 8 | 0.249471384 | 0.249999998 | 0.250000000 |
| 9 | 0.222272679 | 0.222222220 | 0.222222222 |
| Exponential growth model | | | |
| Type | Simulation method | FitCoal | Tabulated FitCoal |
| 1 | 1.441789518 | 1.441679286 | 1.441673129 |
| 2 | 0.616556332 | 0.616509230 | 0.616508244 |
| 3 | 0.376438453 | 0.376743719 | 0.376742859 |
| 4 | 0.269168704 | 0.268510898 | 0.268510338 |
| 5 | 0.207984340 | 0.208373433 | 0.208373096 |
| 6 | 0.170848693 | 0.170519845 | 0.170519652 |
| 7 | 0.144555420 | 0.144603196 | 0.144603091 |
| 8 | 0.125837330 | 0.125751961 | 0.125751908 |
| 9 | 0.111364011 | 0.111402468 | 0.111402445 |
| Bottleneck model | | | |
| Type | Simulation method | FitCoal | Tabulated FitCoal |
| 1 | 1.673321651 | 1.673732814 | 1.673741394 |
| 2 | 0.775051787 | 0.774454488 | 0.774457584 |
| 3 | 0.510960086 | 0.510973534 | 0.510974557 |
| 4 | 0.394262151 | 0.392762187 | 0.392762519 |
| 5 | 0.325116671 | 0.326426872 | 0.326426966 |
| 6 | 0.284039310 | 0.283210170 | 0.283210115 |
| 7 | 0.251717509 | 0.251977260 | 0.251977042 |
| 8 | 0.228240338 | 0.227745526 | 0.227745129 |
| 9 | 0.207539986 | 0.208024514 | 0.208023953 |

| Complex model | | | |
|---|---|---|---|
| Type | Simulation method | FitCoal | Tabulated FitCoal |
| 1 | 1.055607245 | 1.055618949 | 1.055592252 |
| 2 | 0.444340081 | 0.444811898 | 0.444812747 |
| 3 | 0.285522751 | 0.285500848 | 0.285501081 |
| 4 | 0.215995323 | 0.215622334 | 0.215622470 |
| 5 | 0.176447487 | 0.176540313 | 0.176540331 |
| 6 | 0.151750997 | 0.151231386 | 0.151231289 |
| 7 | 0.133173783 | 0.133109963 | 0.133109779 |
| 8 | 0.118845125 | 0.119172606 | 0.119172375 |
| 9 | 0.107810932 | 0.107899199 | 0.107898957 |

**Table S16. Likelihood promotion rate of inferred demographic histories with different inference time intervals of 1000GP populations.**

| Super population | Population | Rate compared with result of (*k*-1) inference time intervals (%) | | | |
|---|---|---|---|---|---|
| | | 2 | 3 | 4 | 5 |
| AFR | ACB | 1384.26 | 350.31 | 60.09 | 18.68 |
| | ASW | 3538.08 | 130.56 | 93.48 | 6.10 |
| | ESN | 714.05 | 35.23 | 126.18 | 17.37 |
| | GWD | 590.15 | 302.50 | 112.43 | 13.01 |
| | LWK | 2925.80 | 91.55 | 70.91 | 5.88 |
| | MSL | 2935.78 | 43.91 | 60.37 | 8.62 |
| | YRI | 861.84 | 166.36 | 69.09 | 7.07 |
| EUR | CEU | 135.51 | 2471.16 | 17.07 | … |
| | FIN | 341.55 | 541.28 | 9.03 | … |
| | GBR | 65.68 | 2900.65 | 22.20 | 2.50 |
| | IBS | 384.43 | 2289.75 | 25.85 | 4.34 |
| | TSI | 236.98 | 2286.49 | 7.09 | … |
| EAS | CDX | 49.99 | 4430.87 | 15.64 | … |
| | CHB | 175.03 | 5343.87 | 10.80 | … |
| | CHS | 111.49 | 4384.47 | 10.26 | … |
| | JPT | 56.37 | 4100.17 | 24.05 | 6.76 |
| | KHV | 153.13 | 4242.29 | 7.67 | … |
| SAS | BEB | 319.95 | 2491.42 | 11.33 | … |
| | GIH | 101.31 | 1982.75 | 8.66 | … |
| | ITU | 261.16 | 1836.98 | 18.17 | … |
| | PJL | 254.33 | 1884.50 | 11.41 | … |
| | STU | 284.42 | 2590.84 | 9.89 | … |
| AMR | CLM | 834.67 | 1867.06 | 17.44 | … |
| | MXL | 347.01 | 4258.47 | 10.01 | … |
| | PEL | 359.65 | 5549.64 | 16.19 | … |
| | PUR | 1335.08 | 967.44 | 22.39 | 11.29 |

**Table S17. Likelihood promotion rate of inferred demographic histories with different inference time intervals of HGDP-CEPH populations.**

| Super population | Population | Rate compared with result of ($k$-1) inference time intervals (%) | | | |
|---|---|---|---|---|---|
| | | 2 | 3 | 4 | 5 |
| AFR | Biaka | 387.67 | 931.28 | 58.4 | 0.42 |
| | Mandenka | 2350.47 | 31.56 | 13.81 | ... |
| | Yoruba | 3157.91 | 24.28 | 67.73 | 0.35 |
| ME | Bedouin | 282.02 | 87.97 | 1.89 | ... |
| | Druze | 33.15 | 218.35 | 1.68 | ... |
| | Mozabite | 275.41 | 130.64 | 2.99 | ... |
| | Palestinian | 195.99 | 102.86 | 1.2 | ... |
| EUR | Adygei | 179.37 | 98.23 | 0.37 | ... |
| | Basque | 336.87 | 42.2 | 0.78 | ... |
| | French | 48.25 | 201.88 | 0.58 | ... |
| | Russian | 75.68 | 97.53 | 0.27 | ... |
| | Sardinian | 105.69 | 649.27 | 1.04 | ... |
| EAS | Han | 59.53 | 201.39 | 0.95 | ... |
| | Japanese | 86.6 | 154.89 | 0.41 | ... |
| | Yakut | 287.46 | 43.19 | 0.23 | ... |
| CSA | Balochi | 40.25 | 244.48 | 0.78 | ... |
| | Brahui | 27.24 | 276.49 | 0.6 | ... |
| | Burusho | 75.72 | 115.73 | 0.21 | ... |
| | Hazara | 95.97 | 159.24 | 1.05 | ... |
| | Kalash | 1465.6 | 11.76 | ... | ... |
| | Makrani | 106.68 | 112.25 | 0.35 | ... |
| | Pathan | 28.68 | 218.83 | 0.31 | ... |
| | Sindhi | 69.31 | 170.14 | 0.29 | ... |
| AMR | Maya | 136.19 | 1813.04 | 5.16 | ... |

**Table S18. Probabilities of each state (the number of ancestral lineages remained) at time *t*.**

| Standard coalescent time | *n* | Number of ancestral lineages remained | | | | | | | |
|---|---|---|---|---|---|---|---|---|---|
| | | 1 | 2 | 3 | 4 | 5 | 6 | 7 | else |
| 0.6 | 30 | 1.92% | 17.22% | 38.16% | 30.57% | 10.39% | 1.62% | 0.12% | <0.01% |
| 0.6 | 40 | 1.64% | 15.62% | 36.9% | 31.85% | 11.78% | 2.03% | 0.17% | 0.01% |
| 0.6 | 50 | 1.49% | 14.67% | 36.06% | 32.55% | 12.68% | 2.32% | 0.21% | 0.01% |
| 0.6 | 180 | 1.11% | 12.05% | 33.27% | 34.28% | 15.53% | 3.37% | 0.37% | 0.02% |
| 0.6 | 200 | 1.09% | 11.96% | 33.15% | 34.34% | 15.65% | 3.42% | 0.38% | 0.02% |
| 0.6 | 220 | 1.08% | 11.88% | 33.05% | 34.38% | 15.74% | 3.46% | 0.38% | 0.02% |
| 0.8 | 30 | 7.32% | 36.36% | 40.14% | 14.22% | 1.86% | 0.1% | <0.01% | <0.01% |
| 0.8 | 40 | 6.73% | 34.93% | 40.68% | 15.36% | 2.17% | 0.12% | <0.01% | <0.01% |
| 0.8 | 50 | 6.38% | 34.05% | 40.96% | 16.08% | 2.38% | 0.14% | <0.01% | <0.01% |
| 0.8 | 180 | 5.44% | 31.41% | 41.56% | 18.27% | 3.09% | 0.22% | 0.01% | <0.01% |
| 0.8 | 200 | 5.4% | 31.31% | 41.58% | 18.36% | 3.12% | 0.22% | 0.01% | <0.01% |
| 0.8 | 220 | 5.37% | 31.22% | 41.59% | 18.43% | 3.15% | 0.22% | 0.01% | <0.01% |
| 1.0 | 30 | 15.99% | 48.88% | 29.75% | 5.09% | 0.28% | 0.01% | <0.01% | <0.01% |
| 1.0 | 40 | 15.18% | 48.17% | 30.73% | 5.58% | 0.33% | 0.01% | <0.01% | <0.01% |
| 1.0 | 50 | 14.7% | 47.72% | 31.32% | 5.89% | 0.36% | 0.01% | <0.01% | <0.01% |
| 1.0 | 180 | 13.35% | 46.29% | 33% | 6.88% | 0.48% | 0.01% | <0.01% | <0.01% |
| 1.0 | 200 | 13.29% | 46.23% | 33.06% | 6.92% | 0.48% | 0.01% | <0.01% | <0.01% |
| 1.0 | 220 | 13.25% | 46.18% | 33.12% | 6.95% | 0.49% | 0.01% | <0.01% | <0.01% |
| 1.2 | 30 | 26.31% | 53.05% | 18.95% | 1.65% | 0.04% | <0.01% | <0.01% | <0.01% |
| 1.2 | 40 | 25.42% | 52.97% | 19.75% | 1.82% | 0.05% | <0.01% | <0.01% | <0.01% |
| 1.2 | 50 | 24.89% | 52.89% | 20.23% | 1.93% | 0.05% | <0.01% | <0.01% | <0.01% |
| 1.2 | 180 | 23.37% | 52.6% | 21.69% | 2.27% | 0.07% | <0.01% | <0.01% | <0.01% |
| 1.2 | 200 | 23.31% | 52.59% | 21.74% | 2.29% | 0.07% | <0.01% | <0.01% | <0.01% |
| 1.2 | 220 | 23.26% | 52.58% | 21.79% | 2.3% | 0.07% | <0.01% | <0.01% | <0.01% |

Note: the shadowed area represents that, when $t \geq 1.0$, the number of ancestral lineages remained is no more than 3 in more than 90% cases.
